# Dating the bacterial tree of life based on ancient symbiosis

**DOI:** 10.1101/2023.06.18.545440

**Authors:** Sishuo Wang, Haiwei Luo

## Abstract

Obtaining a timescale for bacterial evolution is crucial to understand early life evolution but is difficult owing to the scarcity of bacterial fossils and absence of maximum age constraints of the available fossils. Here, we introduce multiple new time constraints to calibrate bacterial evolution based on ancient symbiosis. This idea is implemented using a bacterial tree constructed with mitochondria-originated genes where the mitochondrial lineage representing eukaryotes is embedded within Proteobacteria, such that the date constraints of eukaryotes established by their abundant fossils are propagated to ancient co-evolving bacterial symbionts and across the bacterial tree of life. Importantly, we formulate a new probabilistic framework that considers uncertainty in inference of the ancestral lifestyle of modern symbionts to apply 19 relative time constraints (RTC) each informed by host-symbiont association to constrain bacterial symbionts no older than their eukaryotic host. Moreover, we develop an approach to incorporating substitution mixture models that better accommodate substitutional saturation and compositional heterogeneity for dating deep phylogenies. Our analysis estimates that the last bacterial common ancestor (LBCA) occurred approximately 4.0-3.5 billion years ago (Ga), followed by rapid divergence of major bacterial clades. It is robust to alternative root ages, root positions, tree topologies, fossil ages, ancestral lifestyle reconstruction, gene sets, among other factors. The timetree obtained enables the validation of various hypotheses, such as the survival of life during the late heavy bombardment, the absence of a connection between ancient stromatolites and cyanobacteria, and the presence of aerobic enzymes before the oldest geochemical records of molecular oxygen.

**Significance Statement:** Bacteria, with their vast diversity and ancient history, play a crucial role in shaping Earth’s biogeochemistry. However, the scarcity of fossils complicates the determination of their evolution timescale and its link to Earth’s history. To address this issue, we have devised and implemented novel methods that utilize ancient symbiosis and eukaryotic fossils to calibrate bacterial evolution by molecular clock. We obtain a comprehensive genus-level evolutionary timeline of bacteria that sheds light on their profound influence on the development and diversity of life on our planet, as well as its environmental dynamics. This research greatly contributes to our understanding of microbial evolution and its implications for Earth’s past and present.

## Introduction

Bacteria are the dominating form of life in the first two thirds of the history of our planet, and their evolution and diversification have greatly shaped the geosphere of Earth (1). Many hypotheses have been proposed to interpret outstanding questions like how bacteria affected the evolution of biogeochemical cycles, and when major innovations of bacteria such as photosynthesis occurred and why they did not appear earlier (2–4). Interpreting these questions depends on temporal links between bacterial evolution and the corresponding geobiological events. However, efforts to date the early evolution of bacteria based on relaxed molecular clock (5–7) often fail to reach convergence. A major challenge is that bacteria have rarely left calibrations well suited for molecular dating, i.e., fossils attributable to or biomarkers diagnostic for certain clades (4). Further, because fossils or biomarkers by themselves provide only a lower minimum bound (8, 9), there are no reliable maximum time constraints to calibrate bacterial evolution. Accordingly, the time estimates heavily depend on the maximum age assigned to the root, i.e., the LBCA, whose origin time involves even more debate (10, 11). Moreover, previous studies in dating the early evolution of bacteria (3, 4, 11) are mostly based on site-homogeneous amino acid substitution models that do not well account for among-site compositional heterogeneity and substitutional saturation (12). In deep-time phylogenetics, this may lead to an underestimate of branch length (expected number of substitutions per site) (10), which potentially affects time estimation.

Recent studies have used eukaryotic fossils to date the evolution of Proteobacteria (11) and Cyanobacteria (13, 14) where mitochondrial and plastid-endosymbiosis are believed to have occurred, respectively. These successes imply the value of the abundant eukaryotic fossils in timing bacterial evolution by molecular clock, but more temporal information of eukaryote evolution is waiting to be harvested. One such source is bacteria-eukaryote symbiosis: as such symbioses occurred in multiple ancient bacterial lineages (15, 16), it is tempting to fit it into a recently developed framework called relative time constraint (RTC) dating (17, 18) which constrains the order of divergence by assuming the age of some nodes of the tree to be younger than others. Despite some encouraging progress, the current implementation of this method treats all RTCs as if they are known with full certainty. However, this assumption might be oversimplified because of the uncertainty in identifying the relative time order based on horizontal gene transfer (HGT) (19) or other evolutionary events. Here, we addressed the above challenges by leveraging combined evidence of sequences, fossils, and ancient symbiosis to calibrate an integrative timescale of bacterial evolution. We developed and implemented a framework to propagate date constraints of eukaryotes established by abundant fossils to ancient co-evolving bacterial symbionts and other bacteria by requiring that the origin of symbionts postdates their hosts’ while accommodating the uncertainty in ancestral host reconstruction. We established timelines of bacterial evolution at different taxonomy resolutions and discussed how they may affect our views of the evolution of early life and habitat.

## Results

### A new framework for dating deep-time bacterial evolution informed by host-symbiont association

We overcame the paucity of bacterial fossils in dating the bacterial tree by leveraging the temporal information of eukaryotes recorded in their abundant fossils. Briefly, by incorporating eukaryotes into bacterial phylogeny as mitochondrial lineages affiliated with α-proteobacteria based on mitochondrial endosymbiosis (20, 21), it is tempting to co-estimate the ages of both the eukaryotic host and bacterial symbiont (11) while applying the recently developed RTC-based molecular dating (17, 18, 22). In the context of host-symbiont association, this means constraining the origin time of the symbiont’s crown group to be no earlier than that of the host’s total group, thereby providing maximum time constraints to symbionts whose temporal information may further be propagated across bacteria.

It is well possible that the ancestor of modern symbionts adapted to a different host or even a free-living lifestyle. We formulate a three-step approach accounting for the possibility of host shift of symbionts through ancestral state reconstruction (ASR) and fit it into RTC-based dating in a probabilistic framework, which we call probability-based RTC (pRTC; see Methods for details). First, ASR is performed to estimate the probabilities that the ancestral nodes of modern symbionts are associated with different hosts or are free-living. Second, MCMC molecular dating is performed using mitochondria-originated genes with fossil calibrations but without RTC to generate posterior samples of timetrees where the ages of symbionts and eukaryotic hosts, which are placed as the mitochondrial clade sister to α-proteobacteria (11, 20, 21), are co-estimated. Third, the posterior samples of timetrees collected in Step 2 are examined to remove those where the host’s age is younger than the symbiont’s, weighted by the probability of different states (hosts) taken by the ancestor of modern symbionts estimated in Step 1, using a rejection sampling approach. The idea is further illustrated in Fig. 1A. Assume that ASR infers the ancestral lifestyles of “symbiont1” as *P*(*S*_1_ = *mammal*) = 0.9 and *P*(*S*_1_ = *freeliving*) = 0.1, and for “symbiont2” as *P*(*S*_1_ = *mammal*) = 0.5 and *P*(*S*_1_ = *freeliving*) = 0.5. For the two timetrees in the posterior samples shown in Fig. 1A, the probabilities of acceptance are (0.9 × 1 + 0.1 × 1) × (0.5 × 1 + 0.5 × 1) = 1, and (0.9 × 1 + 0.1 × 1) × (0.5 × 0 + 0.5 × 1) = 0.5, respectively (see Eqs. 3-5 in Methods). If the above procedure is performed for all posterior timetrees collected from Step 2, the posterior dates of all bacterial lineages, particularly the symbiont, might shift toward the present due to the use of RTC informed by host-symbiont association (Fig. 1A). As ancient bacterial symbionts are distributed in different deep-branching lineages (15, 23), this method has a great potential to date the bacterial tree of life.

**Figure 1.**
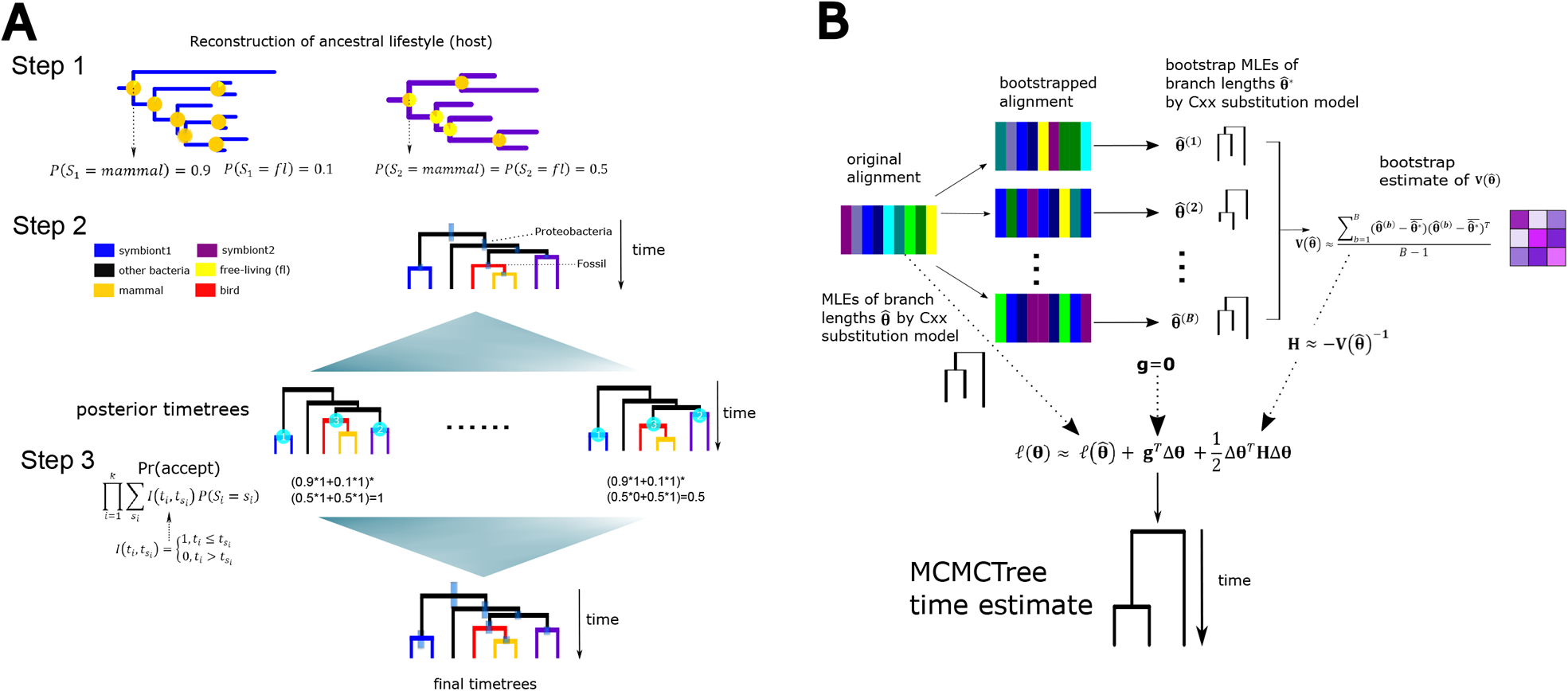
Graphical illustration of the two new methods in molecular dating developed in the present study. (A) Procedure of the pRTC dating informed by host-symbiont association. Suppose that there are two symbiotic lineages, Symbiont1 from non-Proteobacteria (blue) and Symbiont2 from α-Proteobacteria (purple), and most of their members are mammal symbionts (orange) but some are free-living (yellow). First, ASR is performed on the phylogram of 16S rRNA tree to obtain the probability of each ancestral state taken by the LCA of the two “symbiont” clades respectively, i.e., *P*(*S*_1_) and *P*(*S*_2_). Second, a phylogeny consisting of symbionts and mammals is constructed by mitochondria-originated genes, and MCMCTree is run with fossil calibrations but without RTCs. Third, rejection sampling is performed to remove posterior timetrees obtained in the previous step based on the requirement that symbiont’s origin must postdate the host. Specifically, we require that the age of the crown group of the symbionts (node1 and node2) no older than that of the total-group mammals (node3), weighted by *P*(*S*_*i*_ = *mammal*). The use of mammals’ total group is because there could be stem-group mammals that are unsampled or go extinct but that have Symbiont1 or Symbiont2 as symbiont. For the free-living state, we define *I*(*t*_*i*_, *t*_*fl*_) = 1 as the RTC is automatically satisfied for free-living bacteria (see Methods). In the final timetree, the divergence time of symbiont lineages becomes younger than that inferred before host-association is considered, suggesting that incorporating RTC informed by host-association improves the molecular clock inferences. (B) Overflow of the bootstrap-based approach to incorperating branch length estimates inferred under site-heterogeneous profile mixture substitution models (LG+G+Cxx). ***θ*** = (*θ*_1_, *θ*_2_, …, *θ*_*n*_)^*T*^ denotes the parameters to estimate (branch length, i.e., expected number of amino acid substitutions per site) and ℓ(***θ***) is the log-likelihood as a function of branch lengths. Briefly, the maximum-likelihood estimate (MLE) of branch length 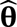 is obtained by LG+G+Cxx, and the bootstrap covariance matrix of branch lengths 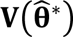 is calculated to approximate the hessian matrix (second-order derivative of the log-likelihood of branch length), which are further used for time estimation by MCMCTree.

### Integrating eukaryote temporal information by a Bayesian sequential approach

We reconstructed the phylogeny of bacteria using the 265 genomes and 60 genes carefully selected in a comprehensive phylogenomics study (24) using the best-fit site-heterogeneous profile mixture model LG+G+C60 (Table S1A; Note S1). We applied the time non-reversible model (25) which infers the root of the bacterial tree as part of maximum-likelihood phylogenetic reconstruction where different root positions may have different likelihoods. This method does not depend on any outgroup, thereby avoiding long branch attraction artefacts caused by using highly divergent sequences to root deep phylogenies. Our analyses rooted the bacterial phylogeny in the neighbourhood of a clade consisting of Fusobacteriota and DST (Deinococcota, Synergistota, and Thermotogota). The analyses also supported the division of most bacterial lineages into Gracilicutes and Terrabacteria, and placed the CPR (Patescibacteria) clade (Fig. S1), initially suggested as the earliest-split bacterial group (26, 27), as a late-split clade sister to Chloroflexi (Fig. S1; Table S2; Note S2). This is generally consistent with a recent study which roots the bacterial tree using an independent method based on gene-species tree reconciliation (24).

To apply pRTC based on host-symbiont co-evolution, we integrated eukaryotes and bacteria in a unified framework to co-estimate their times using the mitochondrial endosymbiosis-based dating strategy (11). Apparently, the ages of symbionts are most strongly constrained if the host date is inferred with a high precision, thus less uncertainty. We therefore took the advantages of the genome-scale data of eukaryotes by employing a Bayesian sequential approach in which the posterior dates of a small number of organisms with many genes are estimated to serve as the time priors of a second analysis with many organisms but a few genes (28, 29). Note that sequential Bayesian molecular dating is conceptually distinct from secondary calibrations (using time estimates from other studies as calibrations) by accommodating the uncertainty in time estimation in the first-step analysis within a Bayesian framework (Note S1.2). We estimated the divergence times of 29 representative eukaryotes with 11 fossil calibrations (Figs. S2-S3) and 320 orthologs conserved across eukaryotes (30). The posterior dates of eukaryote nodes were in agreement with most studies (11, 31–33). Next, we approximated the posterior times of each node in the eukaryote tree by a best-fit probability distribution (Fig. S4; Note S1.2.3), which were further used to calibrate the eukaryote nodes for dating the bacterial tree where eukaryotes were placed as a subtree sister to α-proteobacteria (11, 20, 21).

### A time-calibrated bacterial phylogeny based on ancient symbiosis

To apply the pRTC dating approach based on ancient symbiosis, we first performed comprehensive MCMC-based ASR to infer the ancestral lifestyles of seven well-known ancient clades where symbionts are widely distributed, four at a phylum level (Spirochaetota, Tenericutes, Chlamydiae, and Elusimicrobiota) and three at an order level (Rickettsiales, Holosporales, and Legionellales). The ASR was performed using the 16S rRNA gene, which best captures bacterial lifestyle diversity by including environmental and uncultured samples, based on the lifestyle information of modern members within each of the ancient bacterial clades mainly consisting of symbionts (Fig. S5). We additionally included two genus-level symbionts (*Buchnera* and *Blattabacterium*) which are classical host-symbiont co-evolution examples (16). A total of 19 ancestral nodes consisting of 28 symbiont lineages from the above nine symbiont clades were selected to represent the host-symbiont association-based RTCs in subsequent analyses (Figs. 2, S6; Note S3.1).

**Figure 2.**
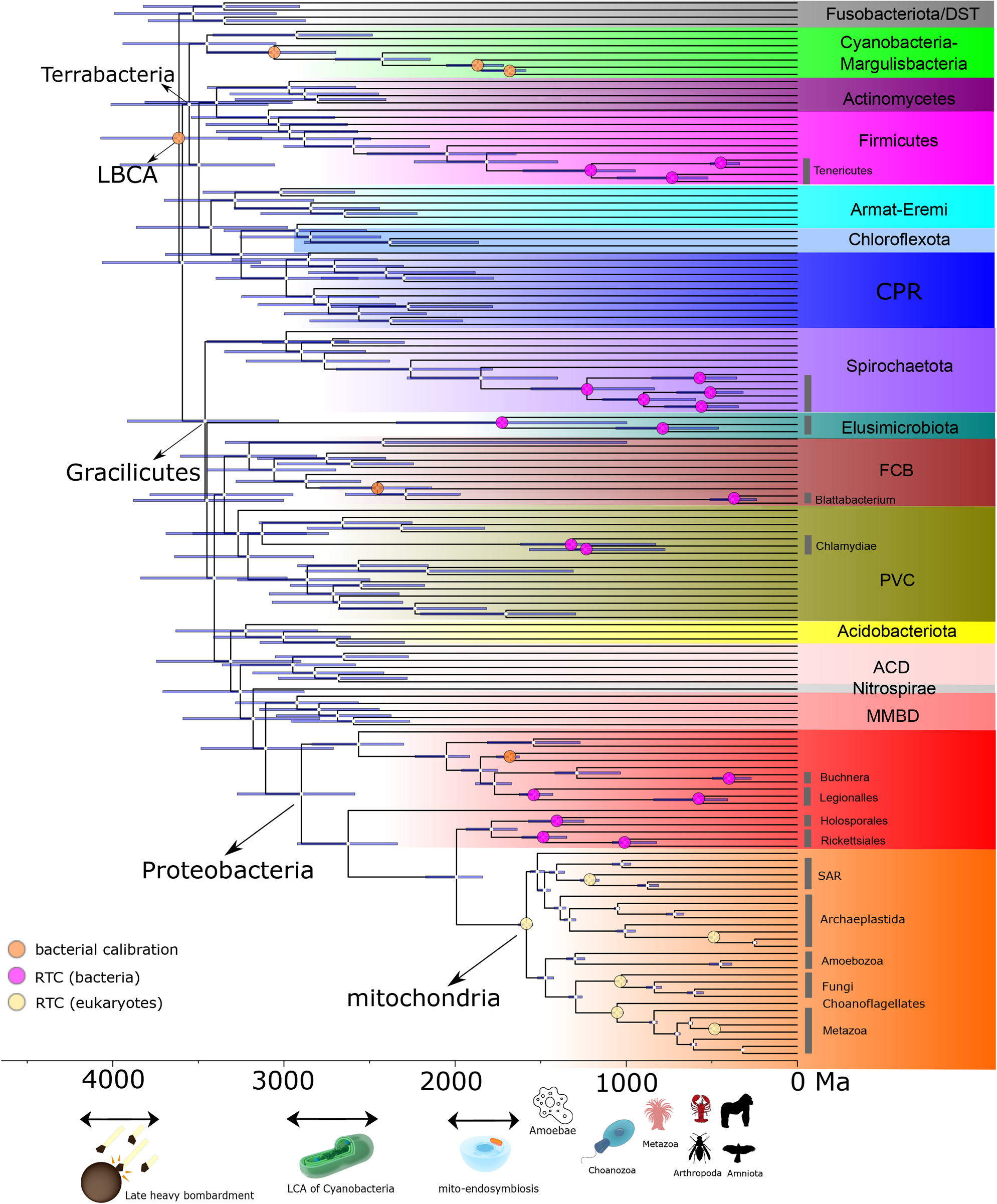
An evolutionary timeline of the bacterial tree of life established by the three-step pRTC procedure. The root position is determined using IQ-Tree’s non-reversible substitution model in maximum-likelihood phylogenomic tree reconstruction using the 19 mitochondria-originated genes. Node bars indicate the 95% HPD interval of age estimates. Nodes with orange circles indicate the bacterial calibration points (including the root). Eukaryotic calibrations obtained from sequential Bayesian dating are given in Fig. S4. RTCs of the 28 bacterial symbionts and five eukaryotic hosts (LCA of eukaryotes, total-group animals, total-group ciliates, total-group insects, crown-group fungi) are denoted by pink and yellow circles, respectively. Because the 28 symbionts are selected from “symbiont” clades, a total of 28-9=19 internal nodes serve as the RTCs for bacteria. The organism images are placed at the origin times of eukaryotes of interest co-estimated with bacterial lineages. Detailed posterior ages for clades of interest are summarised in Table S4. DST: Deinococcota, Synergistota, and Thermotogota; Armat-Eremi: Armatimonadetes and Eremiobacteraeota. CPR: Candidate phyla radiation; FCB: Fibrobacterota, Chlorobiota, Bacteroidota; PVC: Planctomycetota, Verrucomicrobiota, Chlamydiota, and related lineages; ACD: Aquificota, Campylobacterota, and Deferribacterota. MBDD (δ-Proteobacteria): Myxococcota, Bdellovibrionota, Desulfomonadota, and Desulfobacterota. The images of the choanozoan, and the cyanobacteria and mitochondria are credited to Urutseg and Kevin Song respectively under CC BY-SA 3.0 (https://creativecommons.org/licenses/by-sa/3.0/) with slight modifications. All other organism images are distributed under CC0.

Further, we identified 19 genes well conserved across bacteria (Figs. S7-S8) from 108 genes with a mitochondrial origin reported in a recent study (21). We performed molecular clock analysis with these 19 genes on a 148-genome focal dataset consisting of 29 eukaryotes, 28 symbionts, and 91 non-symbionts. The phylogeny was based on the tree reconstructed above using the non-reversible model and with eukaryotes placed as the sister to α-Proteobacteria. The posterior dates of eukaryotic lineages inferred in the first-step sequential analysis above, and five lineage-specific fossil- or biomarker-calibrations within bacteria (see Methods and Note S3.2), were used as time calibrations. We employed MCMCTree’s approximate likelihood method, which obtains a 1000× speed-up without loss of accuracy compared with the exact likelihood method (34), well-suited for our large-scale molecular clock analysis. However, the amino acid substitution models (e.g. LG+G) used to calculate branch lengths in MCMCTree (and many other molecular dating software such as BEAST and MrBayes) do not consider across-site compositional heterogeneity and often show poor model fit in reconstructing deep phylogenies (10, 12).

We therefore developed a bootstrap-based approach to incorperating branch length estimates inferred under the site-heterogeneous profile mixture substitution model LG+G+C60, which better accommodates issues in deep-time phylogenetics like substitution saturation (10), into MCMCTree’s approximate likelihood method (Fig. 1B; see also Methods). Briefly, maximum-likelihood trees were built for 1000 bootstrap alignments, and the inverse of the bootstrap covariance matrix of their branch lengths were calculated to approximate the hessian matrix which is necessary for MCMCTree’s approximate method. Simulations showed considerable improvements in the estimates of both ages and absolute rates by integrating site-heterogeneous models with this approach, partcularly in the absence of internal calibrations or if the divergence times are older than 3.0 Ga (Figs. S9-S10; see Note S4 for details).

Next, we followed Eq. (5) (Methods) to apply a rejection sampling approach to filter out the posterior timetrees obtained without applying RTC from Step 2, based on the probabilities of the ancestor of modern symbionts being associated with different eukaryotic hosts or free-living inferred from Step 1 (Figs. 1A, S8C). Collectively, using a carefully selected focal dating scheme (Data S3), we estimated that the LBCA occurred at around 3648 Ma (95% HPD [highest posterior density] 4111-3162 Ma) (Fig. 2), followed by the split between Terrabacteria and Gracilicutes estimated to be 3629 Ma (95% HPD 4103-3167 Ma). Most major clades, many of which are beyond a phylum level, diverged during 3500-3000 Ma (Fig. 2). All selected symbiotic bacteria lineages (purple nodes in Fig. 2) originated no earlier than 1700 Ma, approximately the appearance time of the LCA of eukaryotes.

We gradually included eukaryotic temporal constraints, the LG+G+C60 substitution model, and host-symbiont association-based RTCs to improve date estimates of bacteria (lower triangle in Fig. 3A). The inclusion of eukaryote’s timing (*Strategy2*) led to a ∼13% decrease in the posterior dates compared with *Strategy1* where the five bacterial calibrations were the only constraints (*Strategy1)*. When the improved branch length estimates based on the LG+G+C60 model was additionally included (*Strategy3*), those nodes between 3000 and 2000 Ma were estimated to be older by 7% (*Strategy2* vs. *Strategy3*). Further, when pRTC was applied on the basis of *Strategy3*, the posterior dates of almost all nodes systematically decreased ∼5% (*Strategy3* vs. *Focal* in Fig. 3A), suggesting that the constraint on the time order imposed by host-symbiont co-evolution was propagated through the bacterial tree of life. Collectively, the combination of the use of time constraints provided by eukaryote temporal information, site-heterogeneous profile mixture models, and host-symbiont association-based RTCs estimated the divergence times of analysed bacteria ∼15% younger than those obtained without any of these settings (*Strategy1* vs. *Focal* in Fig. 3A; this represents a net reduction and some effects cancelled out as illustrated above). As to the posterior rate for each branch, the use of C60 led to an almost doubling of rate estimate (upper triangle in Fig. 3A; *Strategy2* vs. *Strategy3*), indicating that the absolute rates are largely underestimated by the compositionally homogeneous LG+G model. Moreover, when we successively increased the maximum time prior of the root from 4500 Ma to 6000 Ma, the posterior dates of bacterial lineages increased by more than 20% with *Strategy1*, but remained almost unchanged with the focal scheme (Figs. 3B, S11A), indicating that the (maximum) time constraints imposed by eukaryotes effectively constrained the timing of bacterial evolution, such that their time estimates were robust to the root calibration where much debate has arisen (10, 35).

**Figure 3.**
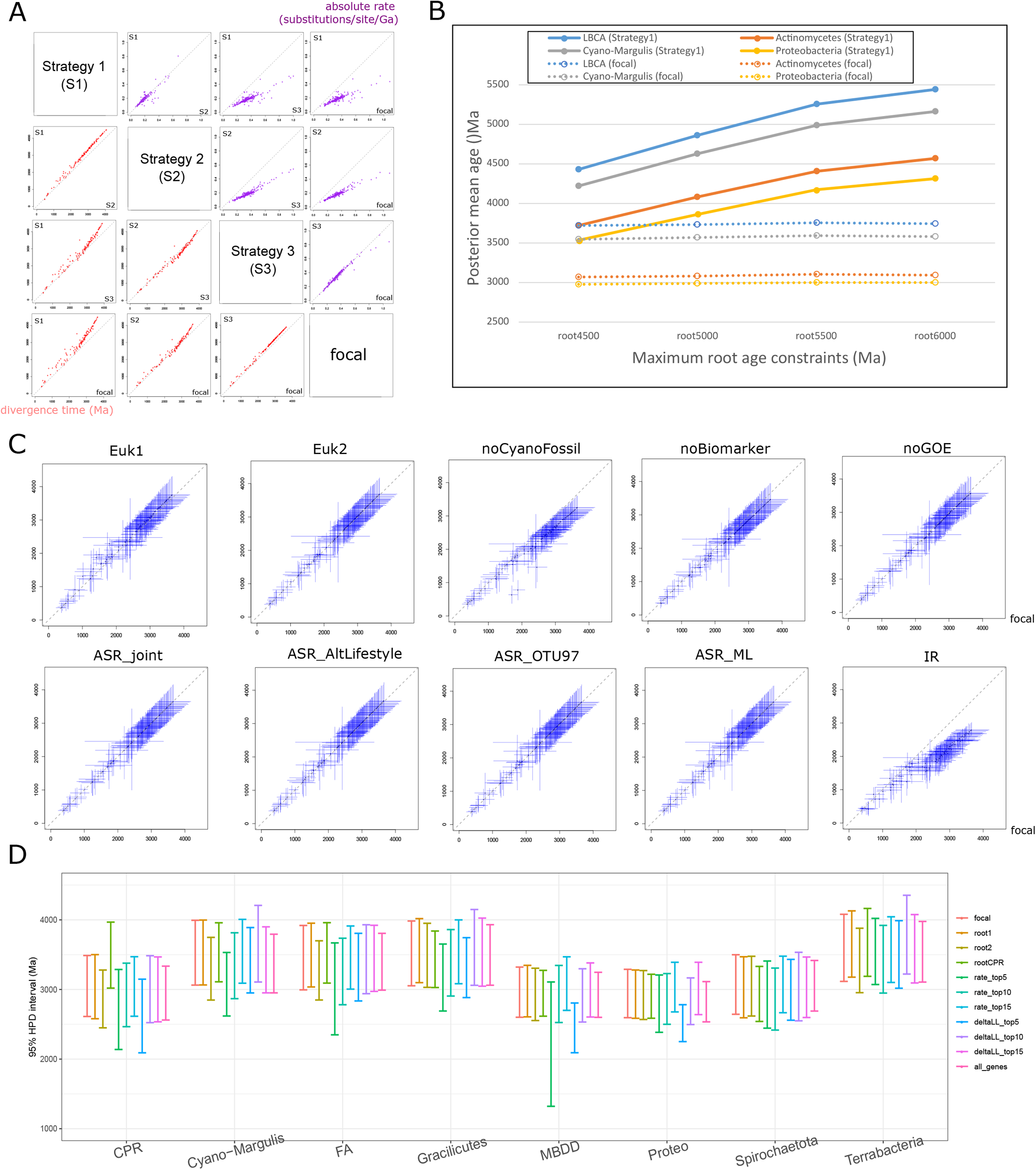
Assessing the uncertainty in Bayesian molecular clock analysis. (A) Comparison of the posterior mean ages (lower triangle; unit: Ma) and rates (upper triangle; unit: number of substitutions per site per Ga) of bacteria estimated with different strategies. *Strategy 1* (traditional strategy used in prior studies): five bacterial calibrations with hard minimum bounds and a soft maximum bound <4.5 Ga at LBCA, no eukaryote timing information, substitution model LG+G, no RTCs. *Strategy 2*: bacterial calibrations, eukaryote times by sequential molecular dating, substitution model LG+G, no RTCs. *Strategy 3*: bacterial calibrations, eukaryote times by sequential molecular dating, substitution model LG+G+C60, no RTCs. Focal: bacterial calibrations, eukaryote times by sequential molecular dating, substitution model LG+G+C60, with RTCs. (B) Posterior mean ages of the selected clades calculated under root maximum prior ages from 4500 to 6000 Ma with the focal strategy (dashed line) and Strategy 1 (solid line). (C) Divergence times of bacteria estimated by alternative schemes (*y*-axis; Data S3) versus the one in the focal strategy (*x*-axis). The blue bars denote the 95% HPD. *Euk1*: the minimum time bound of crown-group red algae alternatively set as 1.6 Ga based on *Rafatazmia*; *Euk2*: the minimum time bound of animal crown group alternatively set as 0.89 Ga (Note S3.2); *noCyanoFossil*: the two cyanobacteria minimum time bounds (total-group Nostocales and Pleurocapsales) removed; *noBiomarker*: the two bacterial biomarker minimum time bounds (total-group Chromatiaceae and Chlorobi) removed; *noGOE*: the minimum time bound based on GOE removed. *Joint_prob*: rejection sampling based on calculating the joint probability (see Fig. S6B) in Eq. (2). *Alt_lifestyle*: ASR inferred with alternative classification of lifestyles of modern symbionts (Note S3.1). *OTU97*: ASR performed with OTU at a cut-off of 97% sequence identity. *ASR_ML*: maximum likelihood algorithm instead of MCMC used in ASR. *IR*: the independent rate (IR) model is used instead of the auto-correlated rate (AR) model. (D) Changes in the posterior dates shown as 95% HPD interval resulting from different root positions of the phylogenomics tree (*root1*, *root2*, and *rootCPR*), and the use of different genes. *rate_top* and *deltaLL_top*: the 5, 10, and 15 (best) top-ranking genes according to relative rate difference between mitochondrial and bacterial lineages or △LL which measures the the degree of species- and gene-tree incongruence (Note S1.2.4). *all_genes*: molecular dating performed on all 32 mitochondria-originated genes conserved across bacteria. The centre is the posterior mean age. FA: Firmicutes and Actinomycetes. MBDD (δ-Proteobacteria): Myxococcota, Bdellovibrionota, Desulfomonadota, and Desulfobacterota.

### Accommodating the uncertainty in divergence time estimation

We evaluated the influence of multiple sources of uncertainties in molecular clock analysis on the posterior dates. First, fossil-based calibrations are known to play a major role in molecular dating (36). The absolute time calibrations consisted of 11 eukaryotic ones, five from within bacteria, and one at the root (LBCA). Replacing the 1.047 Ga-old fossil of *Bangiomorpha*, the most widely recognized oldest crown-group eukaryote (32, 33), by the recently discovered 1.6 Ga-old *Rafatazmia* (37), pushed the LCA of eukaryotes from ∼1600 Ma to ∼2100 Ma (Fig. S3), but had little impact on the time estimates of bacteria (*Euk1* in Fig. 3C). A similar pattern was observed when the recently discovered 0.89 Ga-old sponge fossil instead of the 0.55 Ga-old one (Note S3.2) were alternatively used as the minimum age of the total-group animals (*Euk2* in Fig. 3C). As to the five bacterial calibrations, the two cyanobacteria fossils and the two biomarkers displayed considerable impacts on the posterior dates as their removal led to a decrease of ∼7% in time estimates (*noCyanoFossil* and *noBiomarker* in Fig. 3C). Replacing the 1.6 Ga-old Nostocales fossil calibration (38) by a more disputable estimate based on the 2.0 Ga-old fossil (39) led to an increase in posterior dates by more than ∼10% (Fig. S11B). Except the above, the use of all alternative fossil calibrations returned similar time estimates of both eukaryotes and bacteria (Figs. S11B, S12).

Second, another source of time constraint is the use of ASR-based RTC. In the above analysis, the ancestral states (hosts) between nodes within the same tree were considered independent with each other such that their joint probability can be approximated by the multiplied marginal probability (Fig. S6B). To check if this approximation affected the final time estimates, we directly estimated the joint probability of their ancestral states using a stochastic character mapping (SCM) method (see Methods). This did not change the posterior dates (*joint_prob* in Fig. 3C). Further, while we have conducted comprehensive work in classifying the traits into different hosts or free-living, there exist different interpretations of the phenotypic traits particularly free-living, which might affect ASR. Additionally, we tried different cut-offs of 16S OTU and a different algorithm (maximum-likelihood) in ASR, all of which resulted in highly consistent time estimates (Fig. 3C). Including 13 more symbiont lineages returned similar time estimates (*more_RTCs* in Fig. S12), suggesting that adding host association-based RTCs would not lead to further change in the posterior ages of bacteria.

Third, depencence of the results on the phylogenomic tree topology was investigated. When molecular clock analysis was performed with three alternative root positions, two near the root (*root0*) used in the focal analysis (*root1* and *root2* in Fig. 3D; Fig. S1), and one (*rootCPR*) at the CPR clade as suggested by earlier studies (26, 27), the posterior dates remained similar for most clades, except those that were alternatively placed as an early-split phylogenetic position, particularly the CPR. Alternative phylogenetic positions of mitochondria or between eukaryotic groups (Fig. S8D-S8E) displayed minor impacts on the posterior dates of bacteria (Fig. S11C).

Fourth, the use of different genes in dating may lead to different divergence time estimates. Mitochondrial genes tend to evolve faster than their orthologs in bacterial genomes (11, 20). In theory, the issue can be accommodated by relaxed molecular clock models, but this is not guaranteed (6, 7). Also, MCMCTree assumes a single tree topology for all genes, which might not always hold true due to HGT, unresolved paralogy, or limited phylogenetic signal of individual genes. To minimize the bias caused by gene selection, we ranked the 19 mitochondria-originated genes by i) the relative difference in their estimated evolutionary rates between eukaryotic and bacterial lineages, and ii) △LL, an index to measure the species- and gene-tree incongruence (10). We selected the 5, 10, and 15 top-ranking (best) genes using each of the above two measures and repeated molecular clock analyses (Fig. 3D). Additionally, we analysed the full set of 32 mitochondria-originated genes (*all_genes* in Fig. 3D). Most analyses showed consistent results compared with using the 19-gene dataset; the schemes using only the 5 top-ranking genes (schemes *top5*) displayed larger uncertainty in the date estimates of certain nodes (e.g., MBDD), but this is likely due to their small number of genes.

Fifth, other factors potentially affecting the dating analysis were examined. The largest change in posterior dates were observed when the autocorrelated rates (AR) model, which was consistently supported as the preferred model (Fig. S13), was replaced by the poorly supported independent rates (IR) model: the divergence times of most lineages shifted toward the present by ∼20% (*IR* in Fig. 3C). The priors on the birth-death model which provides time priors for nodes without time calibration and on the lineage-specific rate showed limited impact (Fig. S11B). In addition, analysis with an independently sampled and expanded dataset with 285 bacterial genomes (24) showed similar time estimates (*Secondary* in Fig. S12), suggesting that the time estimates presented in the main analysis were not dependent on a specific taxon sampling. Note that the overall acceptance rate of the rejection sampling was less than 5%, meaning that more than 95% posterior were removed in Step 3. Because running time of MCMCTree is roughly proportional to the number of partitions, to save time, all analyses were performed on a single partitioned alignment, but similar patterns were found when two partitions were used (Fig. S14).

### A taxonomically high-resolution timetree of bacteria

Based on the time estimates from the above analysis where most lineages were sampled at a phylum level, we proceeded with a sequential approach to date a taxonomically higher-resolution timetree of bacteria. Due to the large computational burden associated with Bayesian-based molecular clock methods, we applied the fast dating program RelTime. Speficially, we used lineages included in the 148-genome dataset as the backbone, and used their posterior dates estimated in the above analysis as calibrations to further infer a full-scale bacterial timetree comprising 8898 tips each representing a genus based on GDTB (r95) classification (Figs. 4A, S15; Data S4). The three major clades of Proteobacteria, α-, β-, and γ-Proteobacteria, were estimated to originate ∼2.1, ∼1.6, and ∼1.9 Ga, respectively, greatly postdating the origin of those previously known as δ-Proteobacteria whose origin time was estimated no later than 3.0 Ga (Fig. 4A). This supports the recent proposal to exclude δ-Proteobacteria from the phylum Proteobacteria (40), which was further renamed as the MBDD group (Myxococcota, Bdellovibrionota, Desulfomonadota, and Desulfobacterota) (24). Firmicutes as a whole was the oldest phylum concerning the crown group, originating at ∼3.3 Ga. Family-, order-, class, and phylum-level taxonomy groups were estimated to diverge at 0.90±0.31, 1.4±0.36, 2.1±0.46, 2.5±0.56 Ga (mean ± standard deviation; Fig. 4B). Further, we found a significant correlation between substitution rate of 16S rRNA gene and divergence time at the family level (*p* = 1.98E-44, F-test; Fig. 4C). A simple linear regression analysis suggests an average substitution rate of ∼0.015 nucleotides/site/100 Ma for 16S rRNA gene, but there are great differences between lineages (Fig. 4C). In other words, the divergence time between bacteria may be implied by comparing the sequence of 16S rRNA gene where 1% nucleotide difference roughly corresponds to divergence time of 1/0.015=67 Ma. This indicates a longer time span for 1% nucleotide difference than previously estimated, likely because the bacteria analysed in prior studies are mainly fast-evolving endosymbionts (41–43). The correlation was robust to linear mixed models where the rate differnece of 16S rRNA gene evolution in different lineages was controlled for (Note S1.4; Data S5).

**Figure 4.**
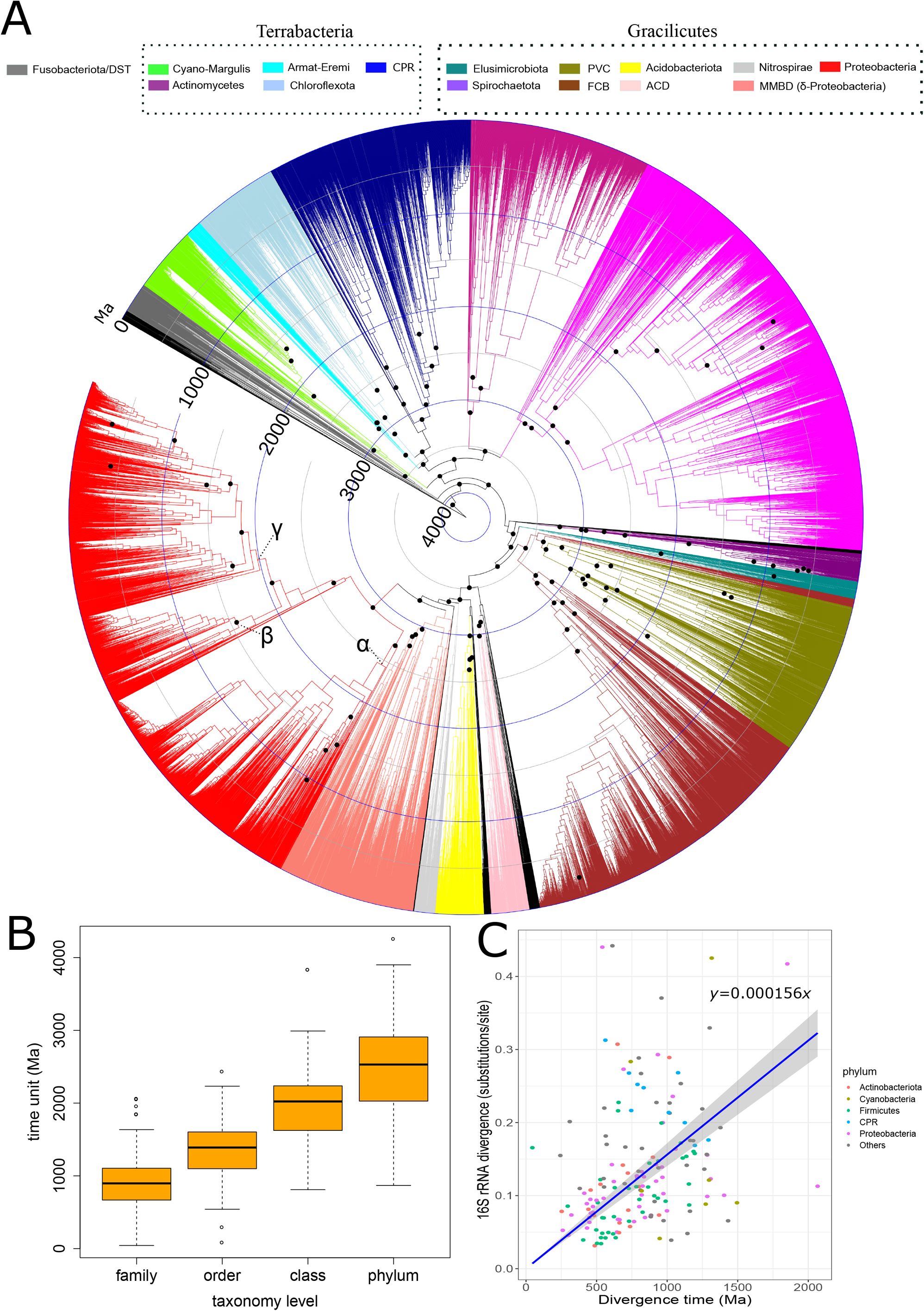
A high-taxonomy resolution timetree of the bacterial tree of life. (A) Divergence times of the 8898-tip bacterial tree, consisting of a randomly selected representative from all genera according to GTDB r95, estimated using RelTime and 118 calibrations (black dots), i.e., posterior dates calculated from the 148-genome molecular clock analysis. Ma: million years ago. (B) The divergence times at a family-, order-, class-, and phylum level inferred from the current study. Only families, orders, classes, and phyla with at least five members are considered. (C) A simple linear regression analysis of the relationship between 16S rRNA divergence (number of substitutions per site; *y*-axis) and absolute divergence time (*x*-axis) at a family level. The intercept is fixed to zero. Shaded areas indicate the 95% confidence interval of the estimated slope. For each family with at least five members, the largest difference in 16S sequence between any of its members is considered the 16S divergence of the family. The analysis is not performed at a higher taxonomic level as 16S sequence likely reaches saturation due to the large divergence time. Detailed results of this simple linear regression and alternative linear models are provided in Data S5 and Note S1.4.

## Discussion

A major challenge in dating bacterial evolution is the paucity of credible calibrations (10, 11), particularly the maximum time constraints. Previously, this was usually done by assigning a somewhat arbitrary maximum age to the root (LBCA), which greatly varies between studies due to the uncertainty in palaeobiological evidence and phylogenetic relationship around the origin of life (10, 44). We addressed this problem by leveraging the timing information of eukaryotes based on ancient symbiosis. This provides two additional layers of timing information besides bacterial calibrations. First, we linked the evolution of bacteria and eukaryotes by mitochondrial endosymbiosis. Because several eukaryotic calibrations have maximum time bounds achieved based on the time estimates of the most recent geological formation that ought to haven fossils of the clade of interest but that does not, this introduces many reliable (maximum) time bounds to indirectly constrain bacterial evolution by informing the substitution rates. Second, based on RTCs informed by host-symbiont association, the timing of eukaryote evolution directly constrains the origin time of symbionts to be no earlier than their eukaryotic host, which is further propagated across the bacterial phylogeny.

The idea of RTC in molecular dating has recently caught much attention (17, 18, 45). However, to our knowledge, none of the available RTC applications has considered the uncertainty in the inference of RTC. In other words, previously the relative time order established by horizontal gene transfer (i.e., gene recipient lineage cannot be older than gene donor lineage) or any other forms of RTC was assumed to be known with certainty, which might be oversimplified due to the uncertainties in identifying these evolutionary events. Our new approach removes this assumption by weighting the different ancestral hosts by their probability established in ASR through a rejection sampling framework. This greatly extends the RTCs based on host-symbiont co-evolution to dating the bacterial tree, which involves many ancient symbioses and large uncertainty in assigning the ancestral host. In addition, prior studies that have applied the idea of host-symbiont association in molecular dating typically used time estimates of the host from other studies (i.e., secondary calibrations) to calibrate symbiont evolution, which may cause wrong time estimates by propagating errors associated with the original time estimates (46). For example, the estimated origin times of animals range from 1355 to 650 Ma according to 11 published recorded in the TimeTree database (47). Apparently, calibrating with different times of animals can lead to dramatically different time estimates of the symbionts. Our strategy co-estimates the origin times of hosts and symbionts based on mitochondrial endosymbiosis (11), avoiding methodological and data differences in comparing times estimates from different studies.

Great care should be taken when interpreting the estimated timeline of the bacterial tree of life and reconstructed evolutionary events taking place anciently. Nonetheless, it is remarkable to note the congruence between time estimates obtained using different time constraints, different root positions, independent sets of genomes, and different settings in molecular clock analysis. Our time estimate of LBCA is ∼0.5 Ga younger than estimated in some studies (4, 48). As evident in Fig. 3A, this might be due to the lack of enough maximum time constraints and the use of poorly fit substitution models for inferring deep phylogenies in these early efforts. Our study also improves molecular clock inferences by using a most-updated phylogeny of the bacterial tree of life. For example, some groups once thought basal in the bacterial tree, e.g., Aquificae (4) and CPR (26), are suggested as later-split lineages in recent (10, 24) and the present study based on improved phylogenetics methods, leading to different time estimates of corresponding lineages.

The estimated timescale of bacterial evolution allows testing hypotheses involving early life evolution. For example, it is generally believed that the late heavy bombardment likely occurred on early Earth ∼4.1-3.8 Ga, during which large asteroids intensely pummelled Earth, melted most parts of its surface, and killed off any forms of life that might have begun to emerge (49, 50). Because primordial form of life likely had existed before LBCA, our estimates for the age of LBCA at 4.0-3.5 Ga hint that life had survived the cataclysm, consistent with modelling studies showing that much of Earth’s crust and the microbes living therein were unlikely affected by the intense bombardment (49). The robustness of the posterior age of the LBCA to the maximum time prior assigned to it (Fig. 3B) provides support to reject the hypothesis that life originated extra-terrestrially (35), where the LBCA should be much older than the age of Earth. Further, under the most parsimonious hypothesis that oxygenic cyanobacteria arose in the cyanobacterial stem lineage (45, 51), our analysis placed the appearance of the first oxygenic cyanobacteria between 3 Ga (total-group Cyanobacteria) and 2.5 Ga (crown-group Cyanobacteria) (Fig. 2). This suggests that one cannot link stromatolites found in ancient sediments dated ∼3.5 Ga to oxygenic cyanobacteria despite their dominance in stromatolites nowadays (52, 53). This is supported by recent evidence showing that stromatolites preserved in Archaean rocks neither required oxygen-producing cyanobacteria for formation (54) nor were necessarily biogenic (55).

The new bacterial timetree also makes it possible to infer the mechanisms that drive the origin of enzymes that involve oxygen. Accumulating evidence from isotope analyses suggests the presence of oxygen on Earth back to 3.1 Ga (56–58), which could be produced by cyanobacteria as mentioned above, and/or abiotically from H_2_O_2_ (59), potentially leading to the formation of “oxygen oasis” thus extracellular access to O_2_. A recent study (60) tracked the origin of most oxygen-utilizing and -producing enzymes families to the Gracilicutes-Terrabacteria split, which according to our estimate is ∼3.6 Ga, predating the above events for ∼0.5 Ga (Fig. 2). This hints that life might have access to oxygen earlier than the oldest geochemical record of the presence of oxygen (60), either extra- or intracellularly [possibly under anaerobic conditions using alternative electron acceptors as observed in Thaumarchaea (61) and the NC10 group (62)] but more likely the latter.

Our work shows that there is much more information beyond bacterial fossils for calibrating bacterial evolution. The estimated timeline of bacterial evolution may well serve as a foundation for studying the evolution of bacteria, their diversification rate, and its correlation with Earth’s geochemistry. It is hopeful that some of these questions can be answered, and competing hypothesis can be reconciled by future molecular clock studies in combination with modern phylogenetics and comparative genomics methods.

## Methods

### Phylogenomic reconstruction

A total of 62 marker genes considered conserved across bacteria were retrieved from a recent phylogenomics study of the bacterial tree (24). Their sequences were aligned using MAFFT v7.22261 (63) and trimmed using TrimAl v1.4 (settings: “-resoverlap 0.55 -seqoverlap 60”) (64). Unrooted phylogenomic trees were constructed based on the concatenated alignment with time-reversible Markov model and each of the three empirical site-heterogeneous profile mixture models C20, C40, and C60 using a PMSF approximation (guide tree built by LG4M+G+I) with 1000 ultrafast bootstraps (65). The model LG+G+C60 was selected as the best-fit model according to AIC and BIC, and the tree constructed by this model was used in the main analysis. Rooted bacterial trees were constructed using the recently developed time-non-reversible model implemented in IQ-Tree (25) that took the unrooted tree as either a fixed tree or a start tree (see Note S2 for details).

### Identification of mitochondria-originated genes conserved across the bacterial tree

We retrieved the sequences of the 108 genes identified to have a mitochondrial origin from the study (21). We identified their homologs in eukaryotes encoded by either the mitochondrial or nuclear genome based on BLAST with an e-value cut-off of 1e-30. Next, we followed (66) to keep 32 genes present in at least one of the following three gene sets namely bac120 (120 genes) (67), Battistuzzi2009 (25 genes) (4), and Coleman2021 (62 genes) (24), which include orthologs well conserved and likely vertically transmitted across bacteria. Note that two of the 62 orthologs originally included in Coleman2021 correspond to the same id (K04485), and were both discarded. For each of the remaining 32 genes, we built a gene tree using IQ-Tree, based on which we further removed i) two genes that did not show monophyly between mitochondria and α-proteobacteria, indicating unresolved paralogy, and ii) 11 genes where mitochondria/α-proteobacteria were placed at a basal position to all other bacteria with different rooting approaches, indicating fast-evolving genes. This returned 19 mitochondria-originated genes conserved across the bacterial tree, which were used in the main molecular clock analysis (Table S3; Fig. S7).

### Ancestral lifestyle reconstruction

The 16S rRNA genes of each interested symbiont group were collected from the NCBI Entrez Nucleotide database with the keywords “taxon_name[ORGN] AND 16S[TITL] AND rrna AND 1000:2000[SLEN] NOT spacer NOT trna” (April 2022). We also retrieved 16S rRNA genes predicted by RNAmmer v1.2 (68) from genomes deposited in Genbank (August 2021). Those without any information of host or isolation source in the metadata were removed (see SI). All remaining sequences were clustered using CD-HIT v4.7 (69) with 98.7% as the identity cut-off (70) into operational taxonomic units (OTUs), and one representative was chosen from each OTU (see SI for alternative strategies in sequence selection). We applied the stochastic character mapping (SCM) (71) to directly sample from the joint posterior distribution, using *make.simmap* from the R package “phytools” v0.2-40 to estimate ancestral hosts (or lifestyles) of modern symbionts. To account for phylogenetic uncertainty, we applied the so-called most-recent-common-ancestor approach by identifying the LCA of the group of species of interest, reconstructing the state of that node, and then summarizing the information across 100 ultrafast bootstrap (UFB) trees reconstructed by IQ-Tree (settings: -m GTR+G). For each of the 100 UFB trees, 100 simulated stochastic character maps were generated using the MCMC algorithm implemented in phytools under the “all-rates different” (ARD) model which allows transition rates at all directions to be different, totalling 100 × 100 = 10000 simulations. All ASR analyses were performed on phylogram—a phylogeny where branch lengths indicate amount of evolutionary change, built by the 16S rRNA gene, as done in previous studies (11, 74).

Specifically, let *S*_*i*_ be the inferred lifestyle (or state) of the *i*^th^ internal node of symbionts, which can take any possible hosts observed in the group of modern symbionts (including free-living), denoted as *s*_*i*_. By SCM, the marginal probability that the *i*^th^ internal node of symbionts takes the state *s*_*i*_ may be approximated by

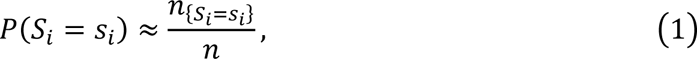

where *n*_{*S*1=*s*1}_ is the number that node i takes the state *s*_*i*_ during *n* rounds of SCMs. Likewise, the same approach can be extended to estimate the joint probability that internal nodes 1 to *k* take states *s*_1_, …, *s*_*k*_ respectively by

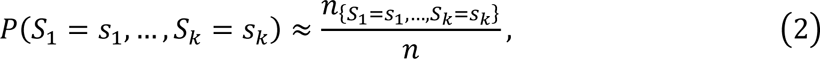

where *n*_{*S*1=*s*1,…,*Sk*=*sk*}_ is the number that the event {*S*_1_ = *s*_1_, …, *S*_*k*_ = *s*_*k*_} happens during *n* rounds of SCMs.

### pRTC informed by host-symbiont association

Denote the data (i.e., sequence alignments) for molecular dating as *D*, and the divergence times as *t* = (*t*_1_, …, *t*_*k*_) where *t*_1_, …, *t*_*k*_ indicate the divergence time of the *k* ancestral nodes of modern symbionts. Let *S*_*i*_ be the inferred lifestyle (or state) of the *i*^th^ ancestral node of symbionts, which can take any possible hosts observed in the group of modern symbionts or free-living, denoted as *s*_*i*_. Let *A* indicate the event that the origin time of all symbionts is no earlier than that of their host, i.e., *A* = {*t*_1_ ≤ *t*_*S*1_, …, *t*_*k*_ ≤ *t*_*Sk*_}. It follows that

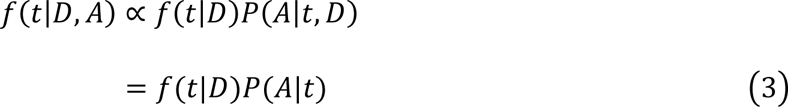

where *f*(*t*|*D*, *A*) is the posterior of divergence times *t* given sequence data *D* and the assumption *A*, *f*(*t*|*D*) is the posterior of divergence times *t* given only sequence data *D*, and *P*(*A*|*t*) indicates the probability *A* is satisfied given time *t*. We further assume that the inference of ancestral lifestyles of modern symbionts depends on only the phylogenetic comparative data estimated by SCM and is independent of time. The second term on the right-hand side of Eq. (3) can be rewritten as

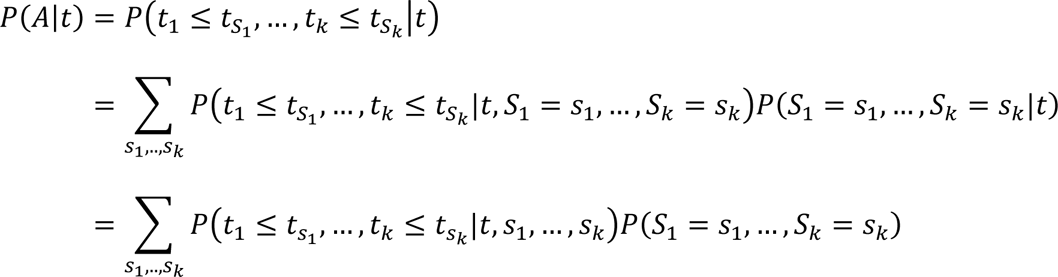

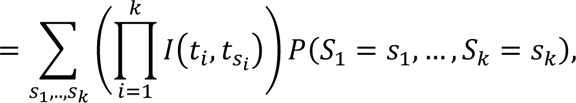

where *I*(*t*_*i*_, *t*_*si*_) is defined as

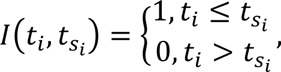

Particularly, we define *I*(*t*_*i*_, *t*_*fl*_) = 1 (free-living ancestral state) since the RTC that symbiont’s age is no older than the host’s is automatically satisfied for free-living bacteria.

Further, assuming that the states of the ancestral nodes of symbiotic bacteria are independent of each other so that 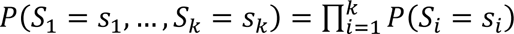, which generally holds if the ancestral nodes are not phylogenetically too closely related (see Fig. S6B for details), the above can be rewritten as

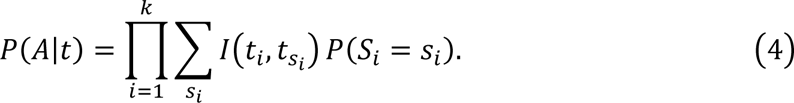

Collectively, by introducing Eq. (4) into Eq. (3), we have

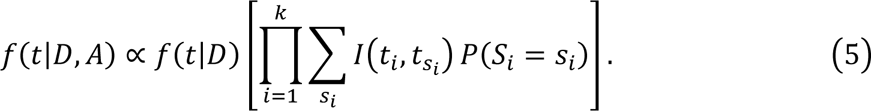

The above indicates that estimating the posterior dates under the assumption that the origin time of all symbionts is no earlier than their hosts’ while considering the uncertainty in the inference of ancestral hosts can be obtained by:

i. Perform ASR to obtain the probability of each ancestral state for selected nodes in the group of modern symbionts of interest.
ii. Perform MCMC molecular clock analysis with fossil calibrations but without any RTC.
iii. For each MCMC posterior sample collected from Step ii), perform a rejection sampling by accepting it with a probability according to Eq. (5) (based on the ASR results obtained from Step i).

Obviously, Step 2 is independent of Step 1, and Step 3 is based on Step 1 and Step 2.

### Sequential Bayesian approach to dating the bacterial tree of life

We applied a sequential Bayesian approach (28, 29) to establish a timeline of the bacterial tree, which allows using the genome-scale data and abundant fossil-based calibrations of eukaryotes to better constrain symbionts’ ages through the pRTC framework (see Supplementary Note 1.3.2 for details). Sequential Bayesian molecular dating contains two steps: i) run a dating analysis of the first subset of data and collect the posterior samples of times and rates, ii) approximate the posteriors of each node in the first-step analysis by a parametric probability distribution and use it as the prior in a subsequent analysis on a second subset of data (see SI for details). We first used MCMCTree from PAML v4.10.0 (75) (see Note S1.2 for details) to date an eukaryote tree with 29 selected species based on our previous study (11) and 320 orthologs (30) with 11 fossil-based calibrations (Note S3.2), then approximated the posterior samples of the dates obtained in the first-step analysis using a best-fit probability distribution available in MCMCTree. The approximated dates were used as the priors to calibrate the bacterial tree using the abovementioned 19 mitochondria-originated genes conserved across bacteria (no overlap in genes between the first- and the second-step analysis).

The posterior dates obtained in the above first-step sequential analysis were used together with the following bacterial calibrations to date the bacterial tree (Note S3.2). The calibrations from within bacteria were the 1.6 Ga-old akinete fossils for the filamentous cyanobacteria Nostocales (38), the 1.7 Ga-old microfossils for the coccoid cyanobacteria order Pleurocapsales (76), the GOE ∼2.32 Ga for photosynthetic cyanobacteria (77), and the hydrocarbon biomarkers for green and purple sulphur bacteria both dated at 1.64 Ga (78). The root minimum was given as 2.32 Ga, the oldest among the above five. A maximum was set as the age of Earth (4.5 Ga) for all the above calibrations within bacteria. If not otherwise specified, in all molecular clock analyses we followed Betts et al. 2018 to apply a uniform distribution as the prior: a soft tail was applied as the upper calibration bound, which means a small probability (by default 2.5%) where the age is beyond the bound, and a hard bound was set as the minimum age meaning that the age cannot be lower than the bound. This makes sense as minimum bounds are based on solid fossil or biomarker records while the maxima are often based on indirect evidence (8, 9).

### A bootstrap-based approach to improving MCMCTree branch length estimate by incorporating the mixture substitution model

We applied MCMCTree’s approximate likelihood method (34, 79) which estimated the branch lengths (expected number of amino acid substitutions per site) in the first step, and in the second step, the branch length estimates, the gradient and the Hessian matrix are used to calculate the likelihood by the second-order Taylor approximation to the likelihood function. Different from previous studies which often use the LG+G substitution model, we applied the bootstrap-based method developed in the present study to integrate the site-heterogeneous profile mixture model LG+G+C60 (80) in MCMCTree analysis. We based our method on the assumption that the maximum likelihood estimates of branch length follow a multivariate normal (MVN) distribution, according to the maximum likelihood (ML) estimation theory stating that ML estimators are asymptotically multivariate normally distributed (81). The idea was originally proposed by Thorne et al. 1998 and later implemented in MCMCTree as the approximate likelihood method (34). The approximate likelihood method (34) first obtains the maximum-likelihood estimates (MLEs) of the branch length 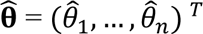, plus the gradient **g** and the Hessian matrix **H**, which are the first and second derivatives of the log-likelihood evaluated at 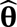 respectively. Next, according to Eq. (1) in the study (34), Taylor expansion at the second order derivative was applied to approximate the log-likelihood

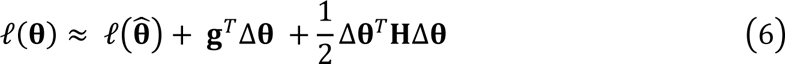

where ***θ*** = (*θ*_1_, *θ*_2_, …, *θ*_*n*_) ^*T*^ denotes the parameters to estimate (branch length) and ℓ(***θ***) is the log-likelihood as a function of branch lengths.

While the gradient is zero at the maximum likelihood estimate (***g*** = ***0***), directly calculating **H** under IQ-Tree’s profile mixture and other complex substitution model is difficult and has not been available. However, note that when the sample size is large enough, the Hessian matrix evaluated at 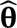 is approximately the negative inverse matrix of the covariance matrix of the MLEs 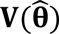) (81):

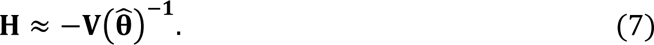

Hence, we instead applied a bootstrap method to estimate 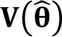 by the bootstrap covariance matrix 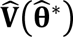 (82) by taking *B* bootstrap alignments and estimating the MLEs of branch lengths 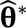:

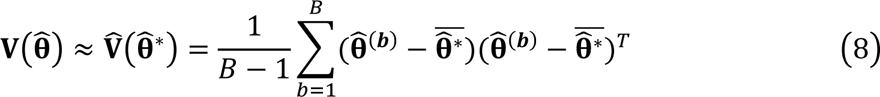

where 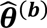 indicate the MLEs of the branch lengths in the *b*^th^ bootstrap sample. Specifically, 1000 bootstraps (*B* = 1000) were performed using the site-heterogeneous profile mixture model LG+G+Cxx with a fixed tree topology by IQ-Tree, therefore generating 1000 bootstrap MLEs for each branch length and allowing calculating the bootstrap covariance matrix 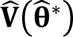. Accordingly, the Hessian matrix can be approximated by solving the inverse matrix of 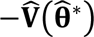. We imported the information of branch length estimates, i.e. the MLEs of branch length, the gradient as a zero matrix, and the Hessian matrix approximated by the above procedure, into a single file *in.BV*, which was used for time estimation using MCMCTree’s approximate likelihood method (34). A series of simulations were performed to show the feasibility of this approach to integrating profile mixture substitution model into MCMCTree analysis (see Note S4).

### A taxonomically high-resolution timeline of the bacterial tree

A representative genome was downloaded from each genus deposited in the Genome Taxonomy Database (GTDB; version r95) (83). We selected 14 out of the 25 orthologs conserved across prokaryotes identified by a prior study (4), the other 11 being removed to avoid squaring of likelihood using the sequential approach [otherwise the same genes are used twice; see Note S1.2.2 and the studies (28, 29)]. These genes were then used to construct a phylogenomic tree based on the first two codon positions of coding sequences using GTR+G model and the “-fast” mode with IQ-Tree to reduce computational burden. We applied RelTime implemented in MEGA X (84), which is based on estimating the relative rates (85), to establish an evolutionary timescale for bacteria at the genus-level resolution. The posterior dates obtained in the 148-genome dataset were approximated by a normal (shown in Fig. 4) or uniform (in alternative analysis) distribution. Because RelTime calculates divergence time by measuring the relative rate between neighbouring branches (85) which is insensitive to overlaps in time priors between parent and child nodes (also known as “truncation effect”; see Note S1.2.2), the age estimates using RelTime may not change much from the specified calibrations. However, to minimize the impact of the overlap in the calibrations between parent and child nodes, we alternatively removed those calibrations where more than 10% overlap in the specified distributions between the parent and child nodes to keep a maximum number of calibrations without significant overlap with each other (Fig. S15D), and re-performed RelTime analysis (Fig. S15A-S15C). Taxonomic classification was according to GTDB version r95 (86). Genetic divergence (substitutions/site) between 16S rRNA sequences was calculated using IQ-Tree under GTR+G+I model.

## Author Contributions

SW and HL conceived the study and wrote the manuscript. SW performed the experiments and analysed the data.

## Data availability

The scripts to do pRTC and bootstrap-based approach hessian for MCMCTree are available at https://github.com/evolbeginner/rrtc and https://github.com/evolbeginner/bs_inBV/ respectively. Other scripts and data are available at https://figshare.com/s/1b39783f1ddda9797914.

## Acknowledgements

We are particularly grateful to Jianhao Lv for the critical comments on the mathematical derivation of the formulae, and the professional insights in statistics. We also thank Daniel Mills and Philip Donoghue for insights in the interpretation of eukaryotic fossils, Mario dos Reis, José Barba-Montoya and Suha Naser for technical support in phylogenetics analysis, and Andrew Meade, Stilianos Louca and Liam Revell for guidance in ancestral state reconstruction. We thank László Nagy, Torda Varga and Tomáš Flouris for kindly sharing with us software. We also thank Luo Lab members Tianhua Liao and Hao Zhang for discussion, and PhyloPic (https://www.phylopic.org/) for the free images of organisms. The work is supported by the Hong Kong Research Grants Council General Research Fund (14116922), the Natural Science Foundation of China (42293294), and the Hong Kong Research Grants Council Area of Excellence Scheme (AoE/M-403/16).

