## Supplementary material for "Dating the bacterial tree of life based on ancient symbiosis": Figs. S1-S15, Tables S1-S4, Notes S1-S5

**This file includes the following items**

- Figures S1 to S15
- Tables S1 to S4
- Notes S1 to S5
- Supplementary references

Note: you can use bookmarks to move between sections.

Figure S1. Determining the root placement of the bacterial tree by IQ-Tree's non-reversible model. All trees are constructed with the 265 genomes and 60 orthologs used in (Coleman et al. 2021). (A) Time-reversible model with a single partition plus the site-heterogeneous models LG+G+C20 (REV_C20), LG+G+C40 (REV_C40), LG+G+C60 (REV_C60), or with partitioning identified by ModelFinder and the best-fit profile site-homogeneous model from LG, WAG, and JTT for each partition (REV_par). The three most likely root positions suggested by tree reconstruction using non-reversible models (see panels B-E and Table S2) are indicated by arrows Root0 (used in the main analysis), Root1 and Root2, all of which are placed near Fusobacteria/DST (deep grey branches; also circled in B-E). The rooting at the CPR, as suggested in earlier studies, is also labelled by a purple arrow. (B) Non-reversible model with a fixed topology according to those constructed by REV_Cxx in panel A and with partitioning. (C) Non-reversible model with a fixed topology according to those constructed by REV_Cxx in panel A and without partitioning. (D) Non-reversible model with a start tree according to those constructed by REV_Cxx in panel A and with partitioning. (E) Non-reversible model with a start tree according to those constructed by REV_Cxx in panel A and without partitioning. Branches with rootstrap support (an index providing information on the support for any given root position using NONREV; see Note S2.2) higher than 10% are labelled with the corresponding rootstrap value (%). Rootstrap support is displayed for only panels (D) and (E) as they cannot be calculated if the tree topology is fixed.

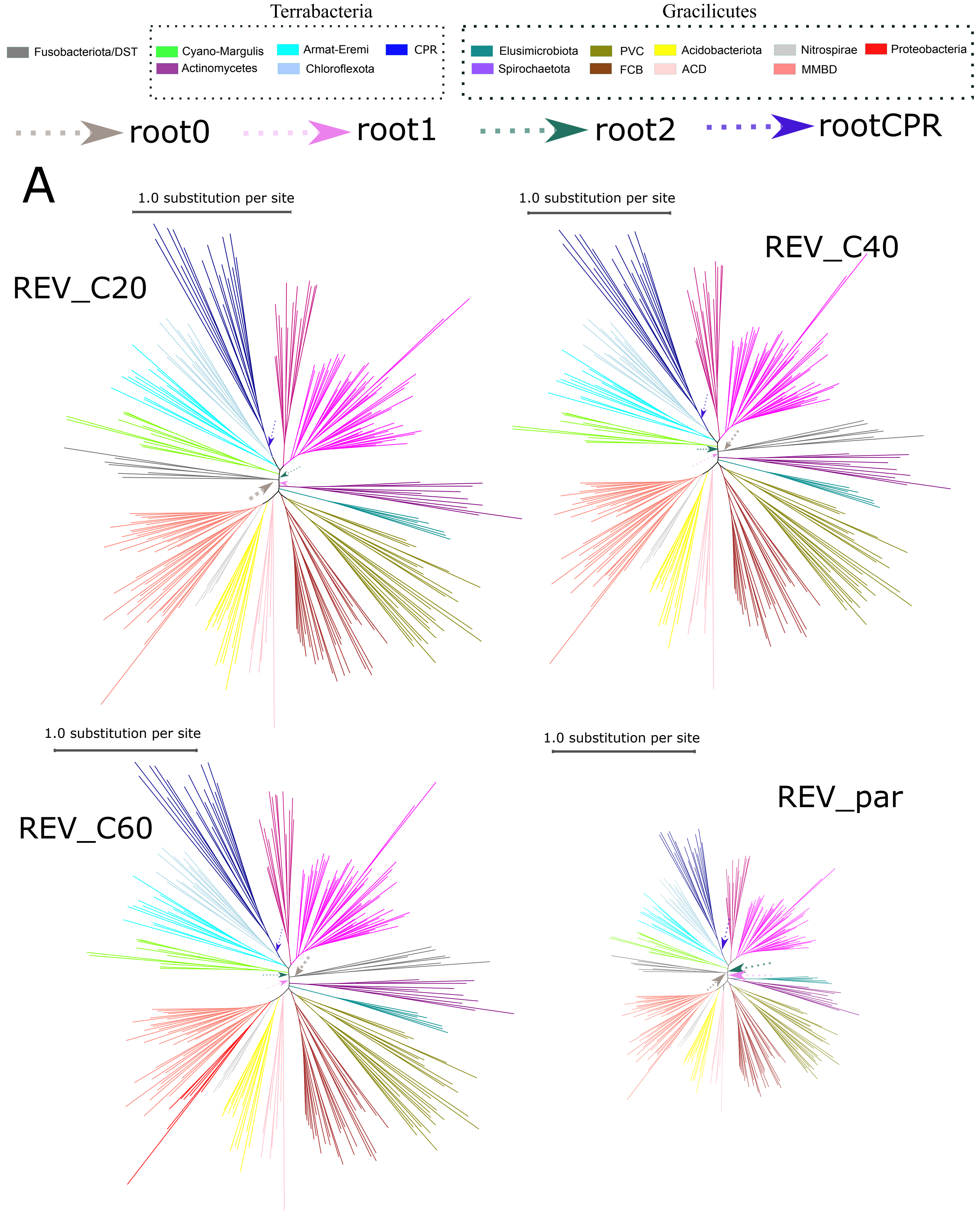

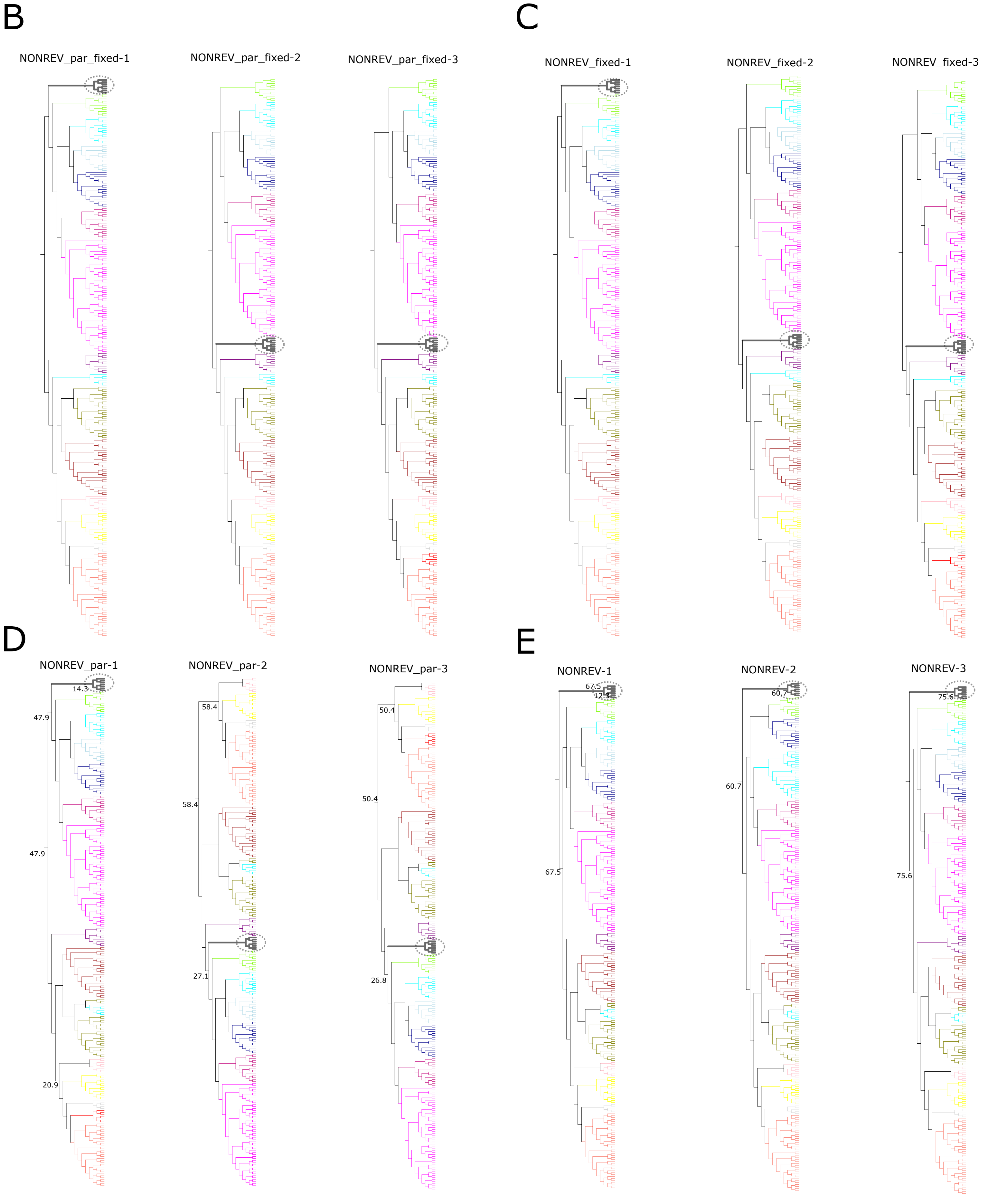
B
NONREV_par_fixed-1
NONREV_par_fixed-2
NONREV_par_fixed-3
C
NONREV_fixed-1
NONREV_fixed-2
NONREV_fixed-3
D
NONREV_par-1
14.3
47.9
47.9
20.9
NONREV_par-2
58.4
58.4
27.1
NONREV_par-3
50.4
50.4
26.8
E
NONREV-1
67.5
67.5
NONREV-2
60.7
60.7
NONREV-3
75.6
75.6

Figure S2. The eukaryote timetree estimated in the first-step sequential molecular clock analysis. (A) Estimated divergence times of 29 eukaryotes based on the 320 orthologs. Circled nodes are those with fossil-based calibrations. The node numbers and the distributions of the posterior dates are indicated. (B) The user-specified fossil-based probability densities (calibration densities) for all calibration points (Note S3.2.1). This is according to the Euk_focal dating scheme (Data S2).

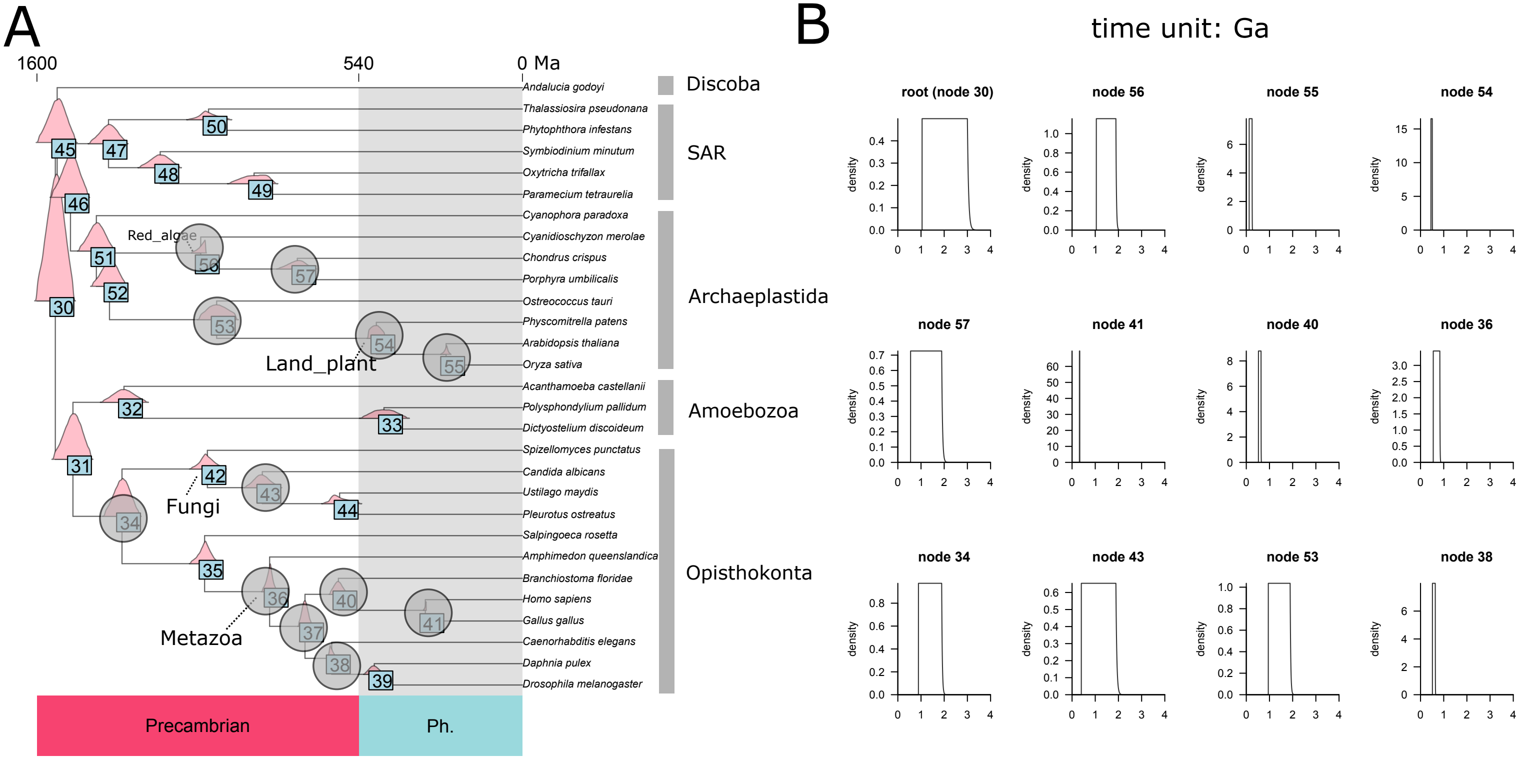

Figure S3. Divergence times of eukaryotes estimated under alternative dating schemes in the first -step sequential molecular clock inference. Detailed information of each scheme is given in Data S2. EUK_soft: a soft minimum is set, meaning a probability of 2.5% that the time is beyond the minimum bound. EUK_AR: the AR model is used. EUK_fossil1: the red algae crown group minimum age set according to the 1.6 Ga-old Rafatazmia fossil. EUK_fossil2: the minimum age of crown -group animals set according to the 0.89 Ga-old sponge fossils. EUK_fossil3: the crown-group land plants maximum age set as 1.042 Ga instead of 0.509 Ga. EUK_fossil4: the minimum age of total -group Nematoda set as 0.528 Ga. Euk_redAlgaeTotalGrp: the minimum age established by the 1.047 Ga -old Bangiomorpha pubescens fossil set on the total group, instead of crown group, of red algae. Euk_Cauchy: for those whose maximum time is constrained by the 1.891 -Ga-old fossil (see Note S3.2) as the maximum time bound, this maximum time bound is removed and instead a truncated Cauchy distribution is applied. Euk_Betts2018: the same calibrations used in the study Betts et al. 2018. Euk_rootMax3500: eukaryote tree’s root maximum of 3500 Ma. Euk_rootMax4000: eukaryote tree’s root maximum of 4000 Ma. Euk_rootMax4500: eukaryote tree’s root maximum of 4500 Ma. Euk_2_partitions: genes are divided into two partitions. Euk_5_partitions: genes are divided into 5 partitions. Euk_20_partitions: genes are divided into 20 partitions.

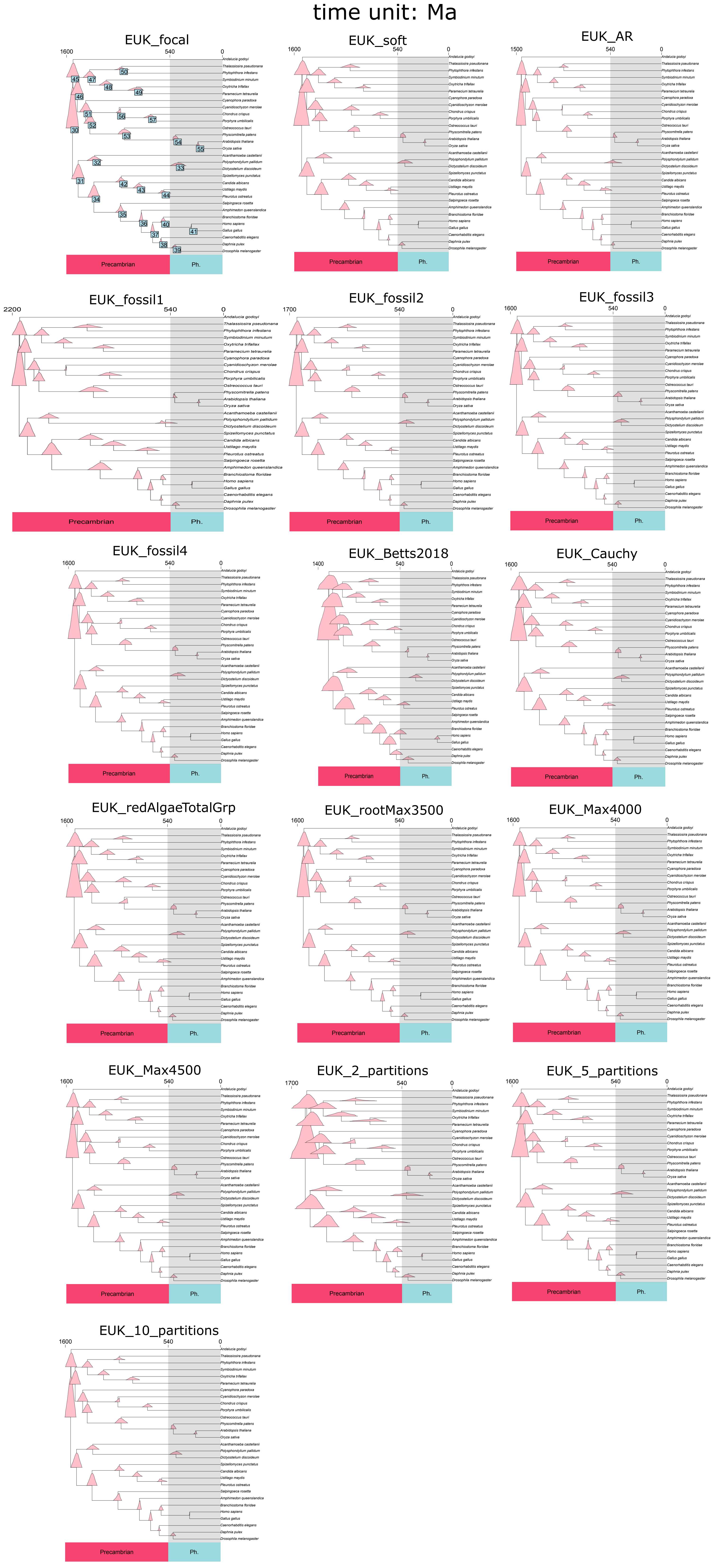

Figure S4. Assessment of the distributions fitted to the internal nodes of the eukaryotic timetree used in the first-step sequential molecular dating. In each plot, the node number is indicated for each plot as “t_nX” where X is the node number shown Fig. S2A, and the parameters of the fitted distribution are shown. The best-fit distributions among skew-t, skew-normal, and gamma distribution based on AIC are plotted in purple. Those estimated when sampling from the posterior ages with MCMCTree using the original calibrations are plotted in blue (first-step sequential dating). The effective time priors (see also Note S1.2.2) for the second-step sequential dating are displayed in red. If all three curves overlap perfectly, the sequential approach would be appropriate to apply to the specific node by using fitted posteriors from the first-step sequential analysis in the second-step sequential analysis. As displayed in Fig. S2A, the posterior ages of nodes 31, 45, and 46 greatly overlap with their parent or child node, resulting in poor approximation. Hence, these three nodes are deemed inappropriate and as such their posteriors are not used in the second-step sequential analysis (see also Note S1.2).

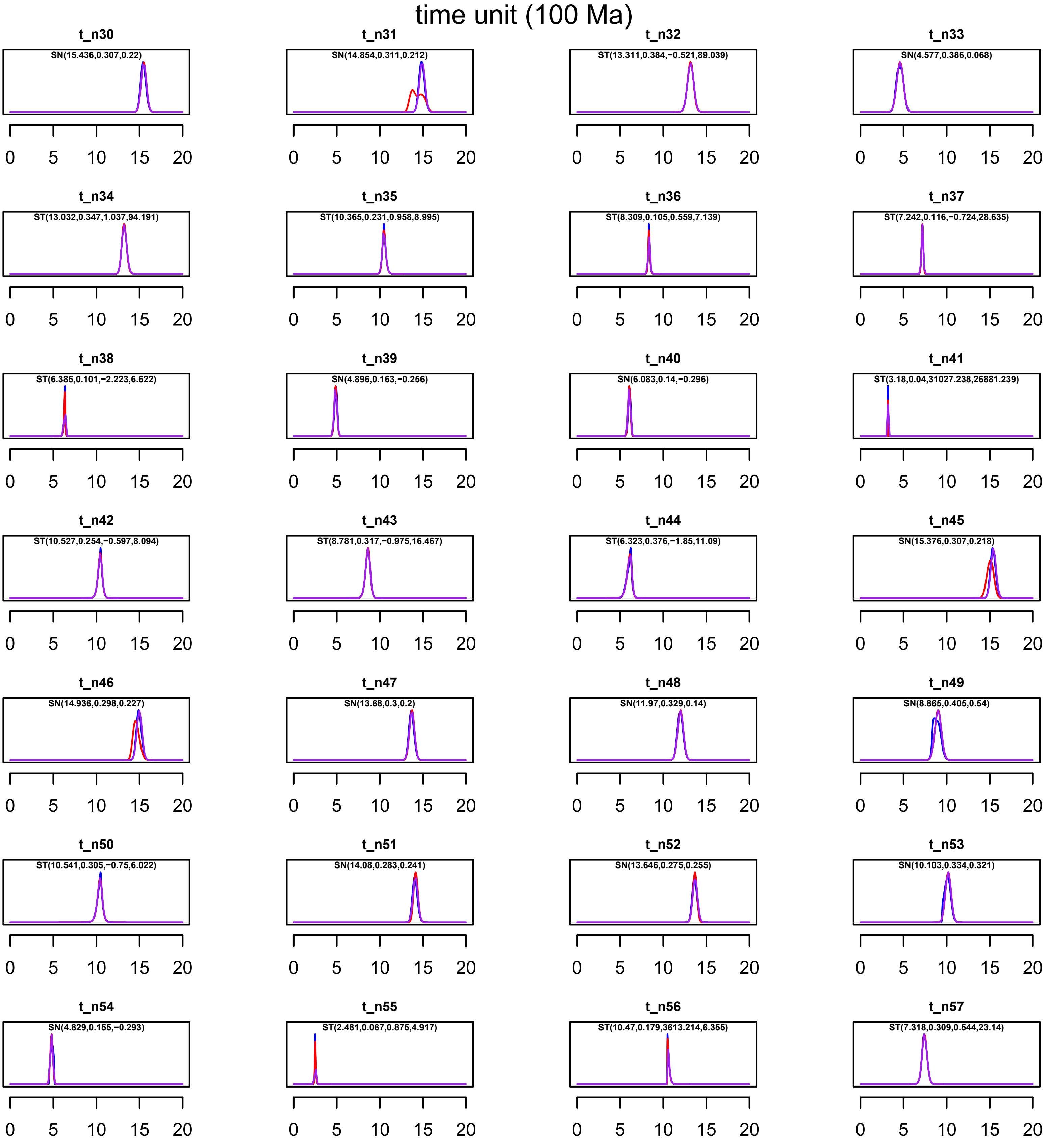

Figure S5. Ancestral reconstruction of the lifestyle in selected bacterial groups by stochastic character mapping (SCM) using the 16S rRNA gene. Pie charts on the nodes indicate posterior probability of each ancestral state at the nodes. The layer (green dots) adjacent to the taxon name represents those with genome available. The black strip in the next layer indicates those used as RTCs in the main molecular clock analysis. The nodes used in RTC-based dating (Note S3.1) are labelled by an arrow. The views of both phylogram, where branch lengths indicate the expected number of substitutions per site, and cladogram, a branching diagram showing only the relationships among clades, are displayed. The ASR analysis is performed using the phylogram. Only seven out of the nine bacterial groups mainly consisting of symbionts involved in RTC-based molecular dating, namely Spirochaetota, Tenericutes, Chlamydiae, Elusimicrobiota, Rickettsiales, Rickettsiales, Holosporales, and Legionellales, are shown. The other two groups are not presented. The reason is because ASR requires at least two different states of the trait (lifestyle), but all of their extant members are isolated from highly specific hosts, Buchnera from aphids and Blattabacterium from cockroaches and termites (see Note S3.1).

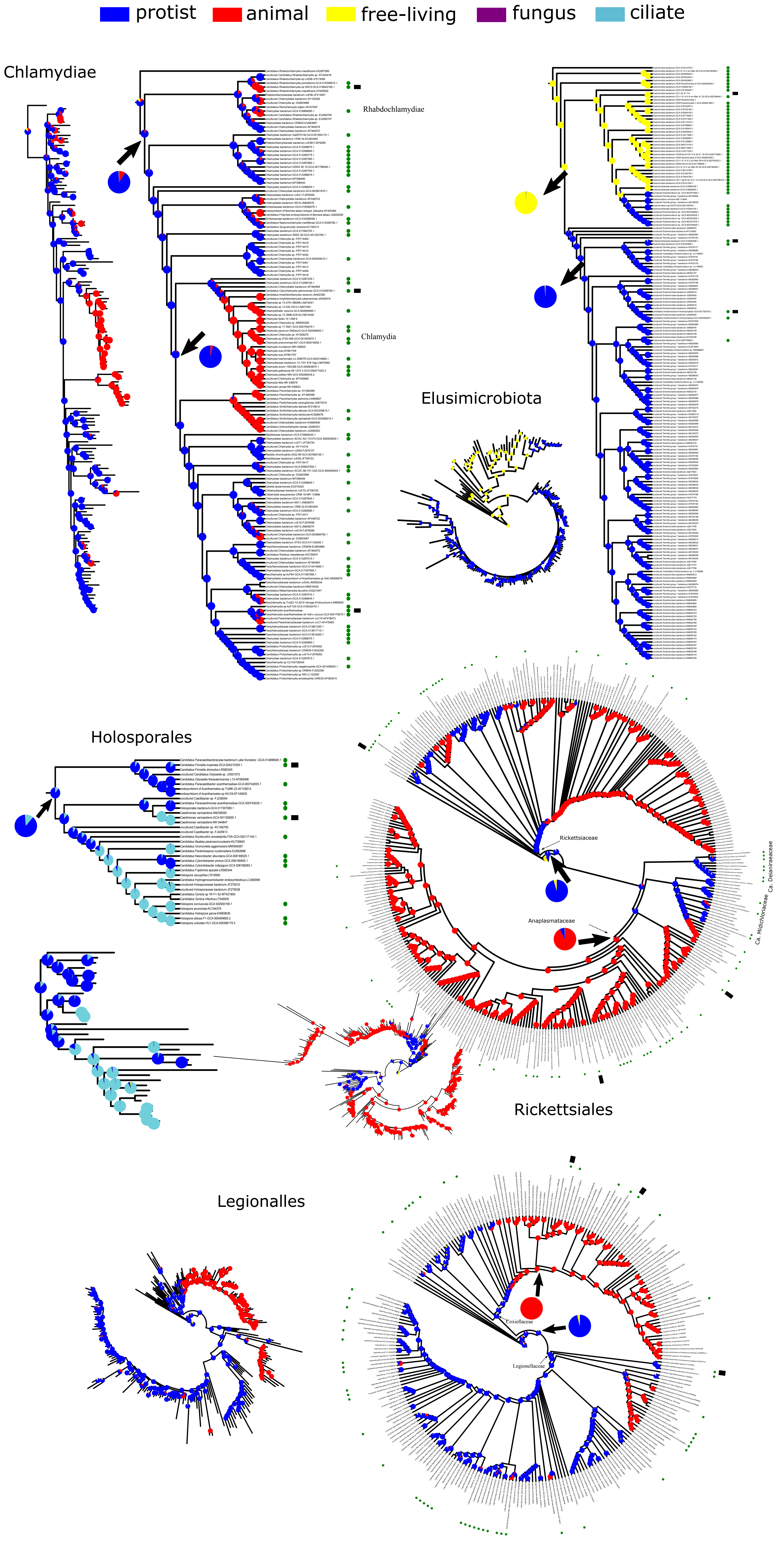

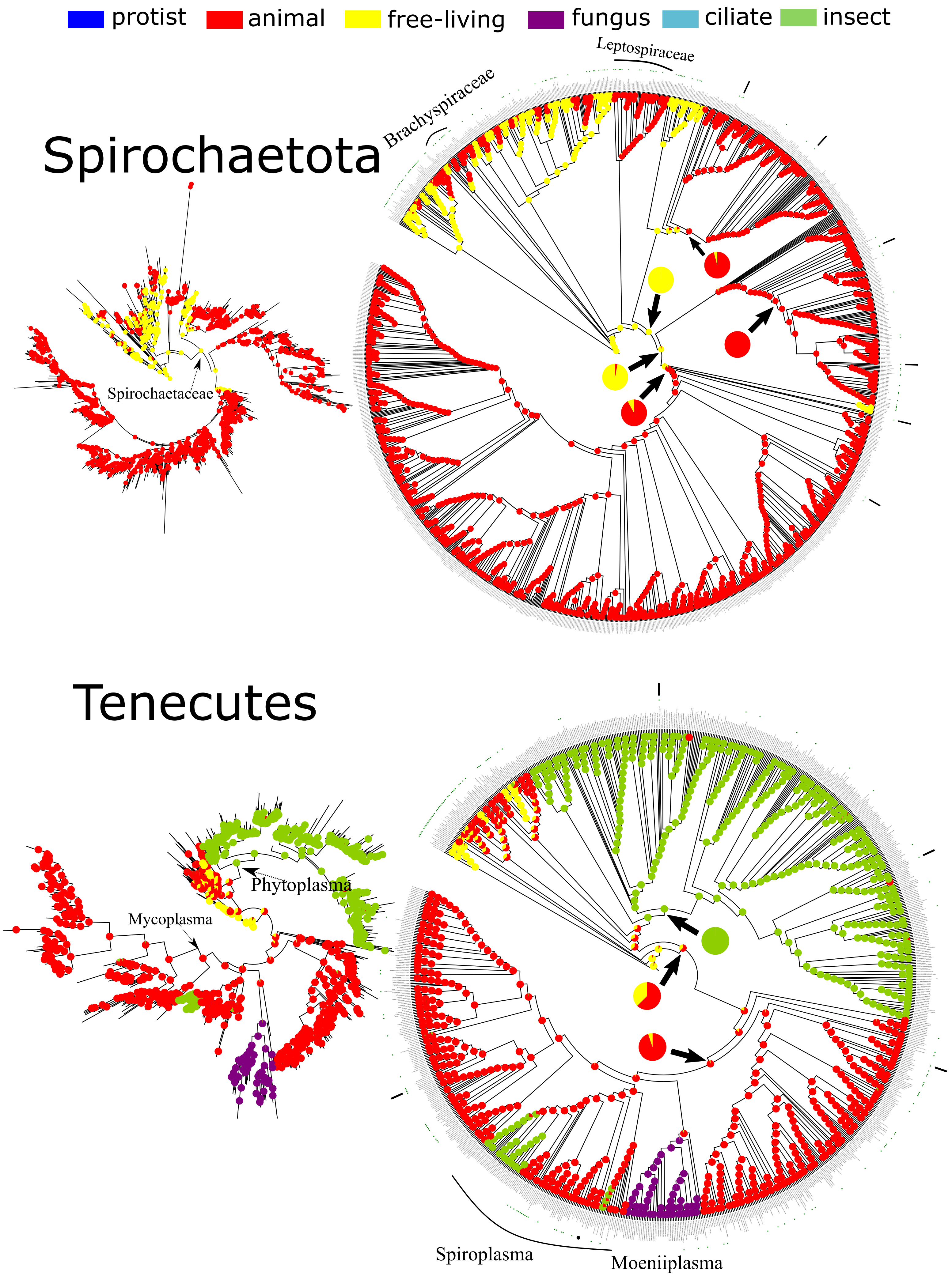

protist
animal
free-living
fungus
ciliate
insect
Spirochaetota
Leptospiraceae
Brachyspiraceae
Spirochaetaceae
Tenecutes
Phytoplasma
Mycoplasma
Spiroplasma
Moeniiplasma

Figure S6. The procedures of selecting symbiotic bacteria in ASR and approximating the joint probability of ancestral states by assuming independence of ancestral states. (A) The general principle of selecting the symbiotic bacteria for pRTC dating based on lifestyle ASR with the 16S rRNA gene. Details are given in Note S3.1.1. (B) An example of how the joint probability of ancestral states is approximated by assuming that the independence of ancestral states of internal nodes in Rickettsiales. Two internal nodes are included, and their ancestral states are denoted by $S_1$ and $S_2$, respectively. In the table at the top, the joint probability, $P(S_1, S_2)$, and marginal probability $P(S_1)$ and $P(S_2)$, are directly estimated by performing the stochastic character mapping (SCM) procedure 10000 times. In the table at the bottom, it is assumed that $S_1$ and $S_2$ are independent, thus the joint probability $P(S_1, S_2)$ is approximated by $P(S_1) \times P(S_2)$. For example, as calculated from the top table, $P(S1 = animals) = 0.9382$ and $P(S2 = protists) = 0.9176$. Assuming the independence of $S_1$ and $S_2$, the joint probability $P(S_1 = animsl, S_2 = protists) = 0.9382 \times 0.9176 = 0.86089232$ in the bottom table, compared to 0.0860 as estimated directly from SCM in the top table. In fact, ancestral states are not independent on each other in the same phylogeny. But when the evolutionary distance between ancestral nodes is large enough, which is the case in our analysis as closely related symbionts are not considered in ASR, the assumption of independence makes sense and simplifies calculation.

A

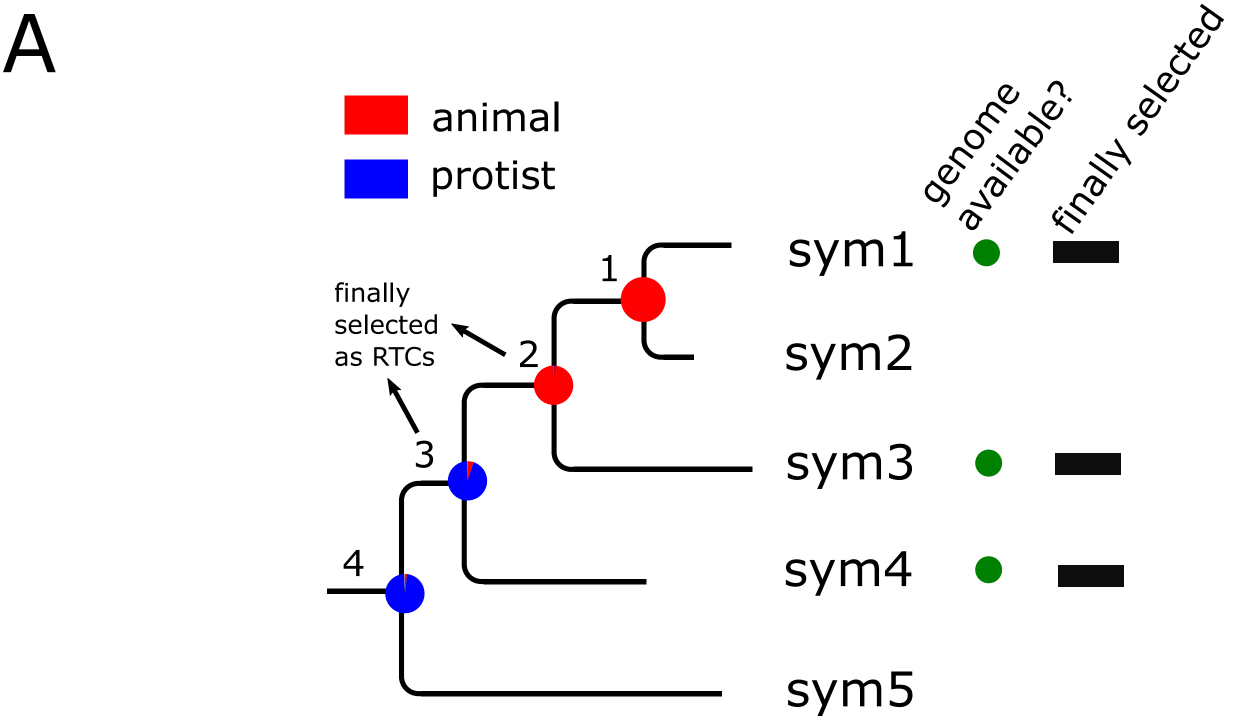


B

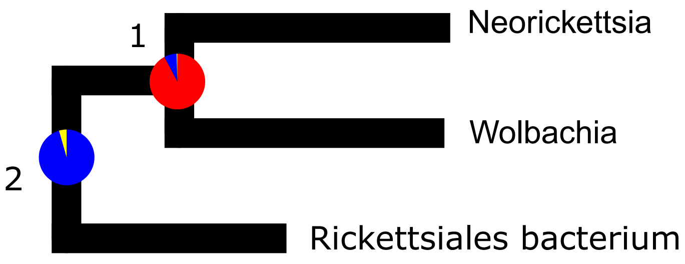


P(S1,S2) estimated directly by SCM

| | S2=animals | S2=pro�sts | S2=fl | marginal of S1 |
|---|---|---|---|---|
| S1=animals | 0.003 | 0.86 | 0.0752 | 0.9382 |
| S1=pro�sts | 0 | 0.0565 | 0.003 | 0.0595 |
| S1=fl | 0.0012 | 0.0011 | 0 | 0.0023 |
| marginal of S2 | 0.0042 | 0.9176 | 0.0782 | |

P(S1,S2) approximated by P(S1)*P(S2)

| | S2=animals | S2=pro�sts | S2=fl | marginal of S1 |
|---|---|---|---|---|
| S1=animals | 0.00394044 | 0.86089232 | 0.073367 | 0.9382 |
| S1=pro�sts | 0.0002499 | 0.0545972 | 0.004653 | 0.0595 |
| S1=fl | 0.00000966 | 0.00211048 | 0.00018 | 0.0023 |
| marginal of S2 | 0.0042 | 0.9176 | 0.0782 | |

Figure S7. The individual gene phylogeny of the 32 mitochondria-originated genes conserved across the bacterial tree. Blue and red branches respectively denote mitochondria and α-Proteobacteria. The tree is rooted using the midpoint approach where the root is placed halfway between the longest tips. (A) The 19 genes selected for the main analysis. (B) The other 13 genes that are excluded from the main analysis due to the fast-evolving mitochondrial sequences (mitochondrial sequences at the basal of the tree), non-monophyly of mitochondria and α-Proteobacteria, or unresolved paralogy (see Table S3).

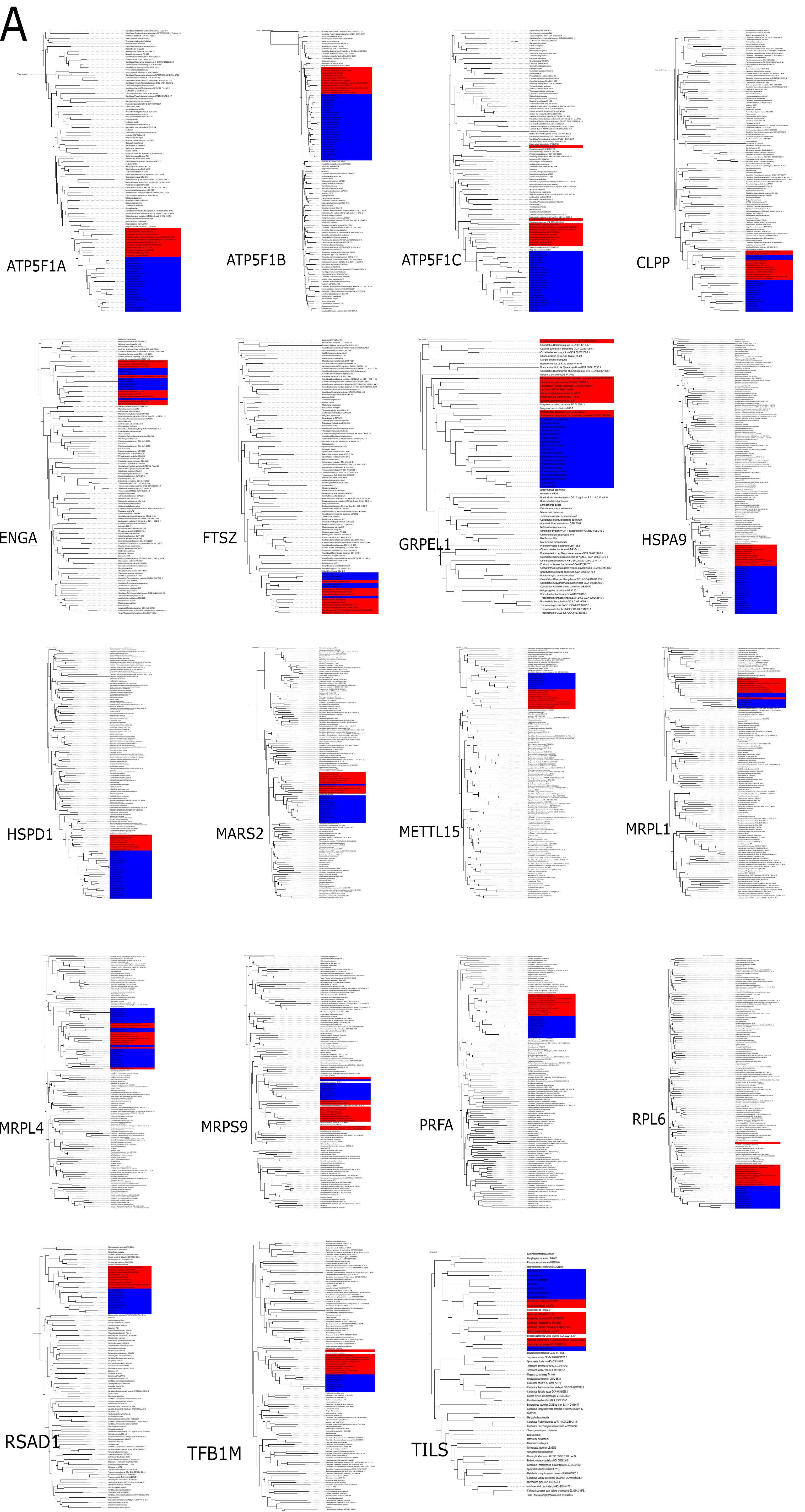

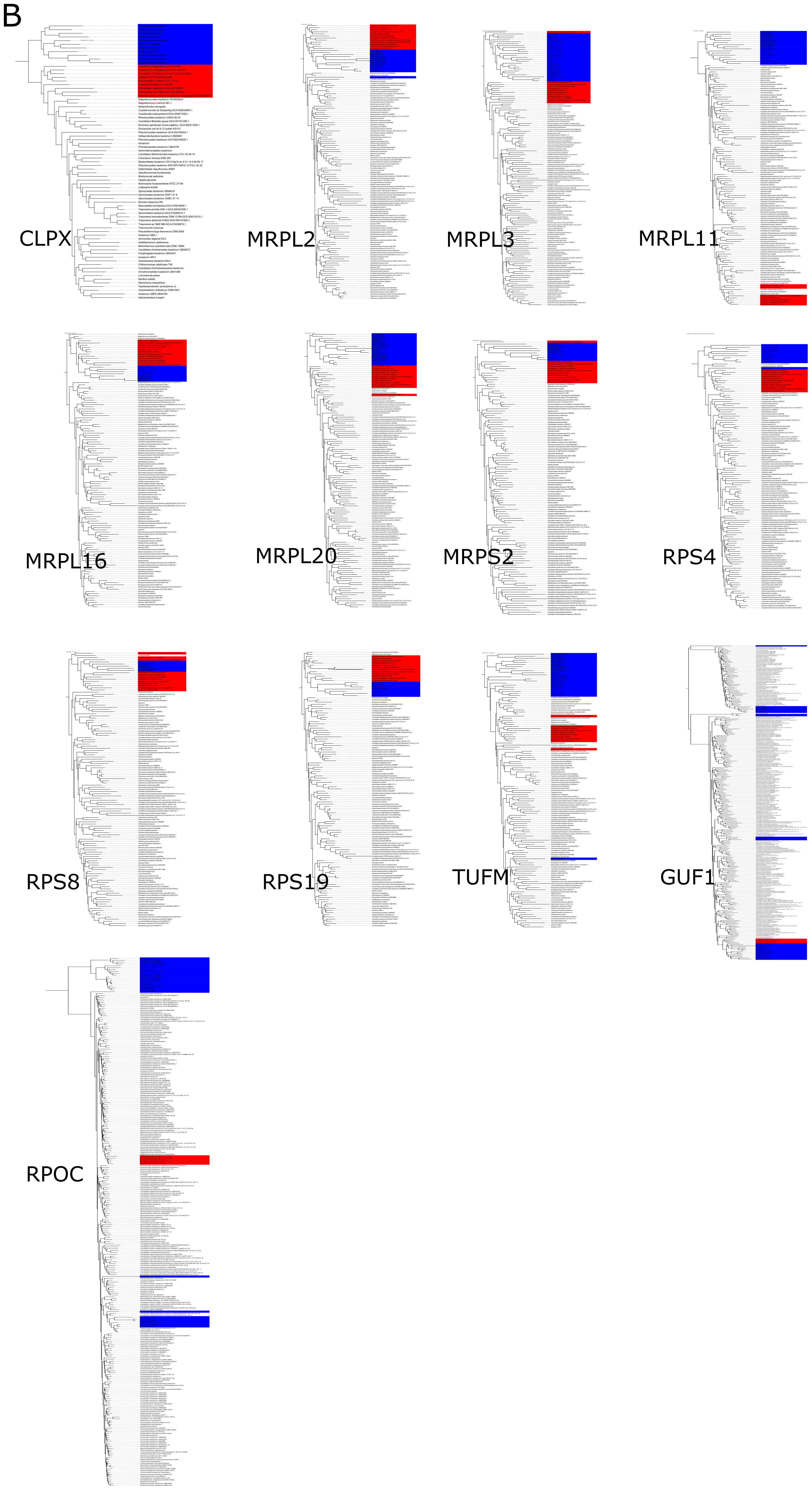
B
CLPX
MRPL2
MRPL3
MRPL11
MRPL16
MRPL20
MRPS2
RPS4
RPS8
RPS19
TUFM
GUF1
RPOC

Figure S8. Calibrations and RTCs used in the focal dating analysis. (A) Phylogenetic placement of all calibration points, inc luding one at the root, five bacterial ones (orange), and 25 from within eukaryotes (yellow) obtained from the first-step sequential analysis, as well as 19 RTCs from "symbiont clades" within bacteria. The presence/absence of the mitochondria -originated genes are also shown, with the 19 genes used in the focal analysis shown in green. The two genes (RPOC and GUF1) with unresolved paralogy are not shown (Table S3; Fig. S7). (B) The user-specified probability densities (calibration densities) for all bacterial calibration points. This is according to t he dating scheme Euk_focal (Data S2). Note that the eukaryotic calibrations used in the 148-genome molecular dating is shown in Fig. S4. (C) The 19 selected RTCs (r1 - r19) and the probability of each of the inferred ancestral hosts (lifestyles) obtained fro m ASR (see also Fig. S5). (D) Four alternative topologies and the reference topology of the eukaryote tree, as used in the study (Wang and Luo 2021). (E) The reference and alternative (Mito-Rick in Fig. S11) phylogenetic position of mitochondria. The refer ence tree topology is used in the main analysis.

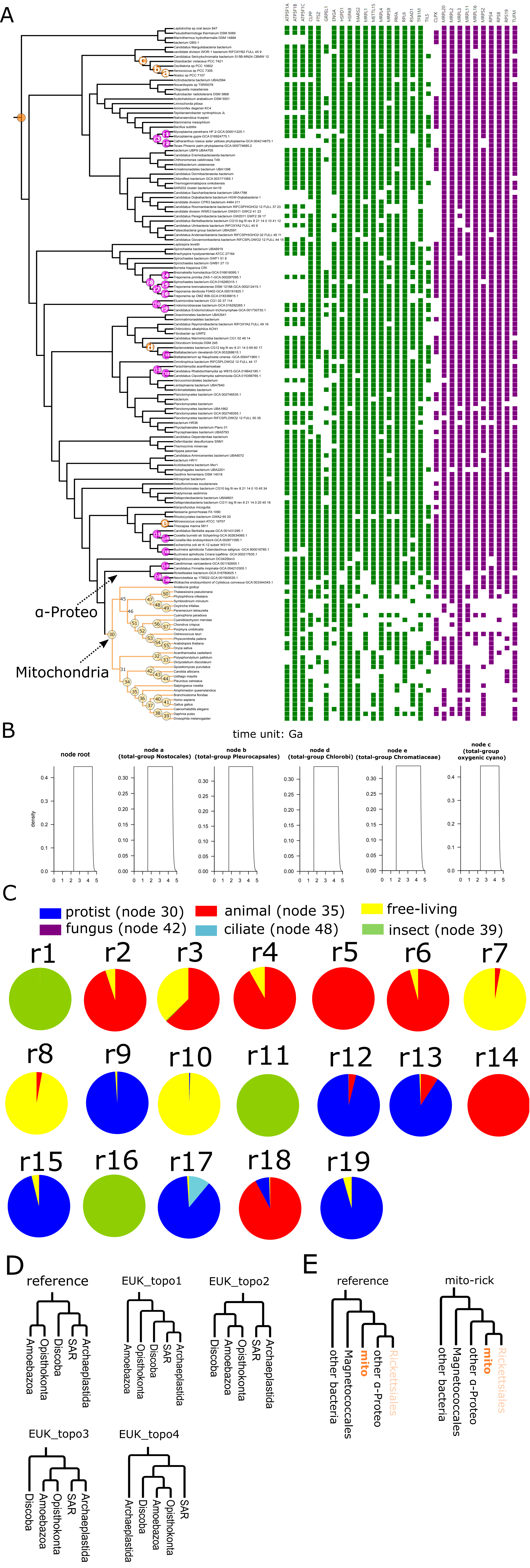

Figure S9. Comparison of the time estimation by different substitution models with simulated sequences. For each substitution model, 30 timetrees each with 30 tips are simulated with a birth-death process under the four root ages (1.0-4.0 Ga) using TreeSim. The sequences are simulated using IQ-Tree's AliSim with compositionally homogeneous model LG+G{1.0} (A), and compositionally heterogeneous model LG+G+C40{1.0} (B) and LG+G+C40{0.5} (C), respectively. Details of simulation are given in Note S4. Red and green boxplots indicate those estimated with a single root calibration and with three calibrations (one at the root, and two internal calibrations set at the 1/3 and 2/3 age quantiles of the true timetree used in simulation). The time priors of all calibrated nodes are set to be uniform within the interval [true_age - (true_age/5), true_age + (true_age/5)]. Two indices are used to measure the accuracy of time estimates of all nodes as compared with the true ages. BSD: branch score distance. Reldiff: relative difference of the dates (see Note S4.4 for definition). Each boxplot contains 30 values representing the above two indices based on MCMCTree analysis with the corresponding model on 30 simulated datasets. LG+G (MCMCTree): branch lengths and hessian both directly calculated by MCMCTree under LG+G. LG+G (bs): branch lengths calculated by IQ-Tree's LG+G and hessian approximated by bootstrapping. LG+G+C20 (bs): branch lengths calculated by IQ-Tree's LG+G+C20 and hessian approximated by bootstrapping. LG+G+C40 (bs): branch lengths calculated by IQ-Tree's LG+G+C40 and hessian approximated by bootstrapping (see also Note S4). P-values are obtained with a Wilcoxon signed-rank test. *: P-value < 0.05, **: P-value < 0.01, ***: P-value < 0.001, ns: non-significant.

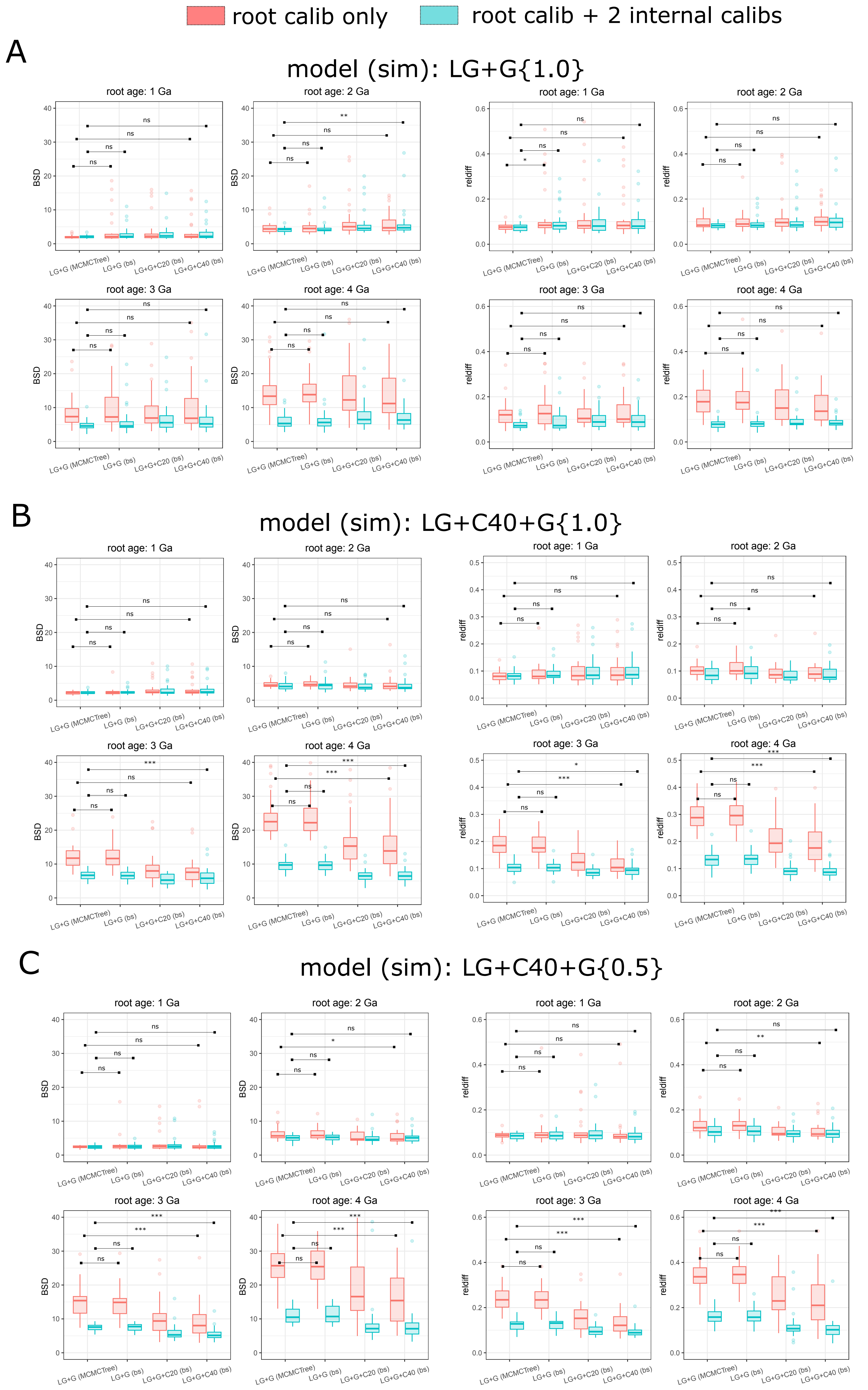

Figure S10. Comparison of the estimated average absolute rate (“mu” in MCMCTree’s output) by different substitution models wi th simulated sequences. The unit for the absolute rate is number of amino acid subs titutions per site per Ga. The true value of the absolute rate used in all simulations is 0.25 substitutions/site/Ga, which is indicated by the grey dashed line. For each MCMCTree analysis, the mean o f the estimated absolute rate across all branches is plo tted in boxplot. Details of simulation are given in Note S4. Abbreviations are the same as Fig. S9.

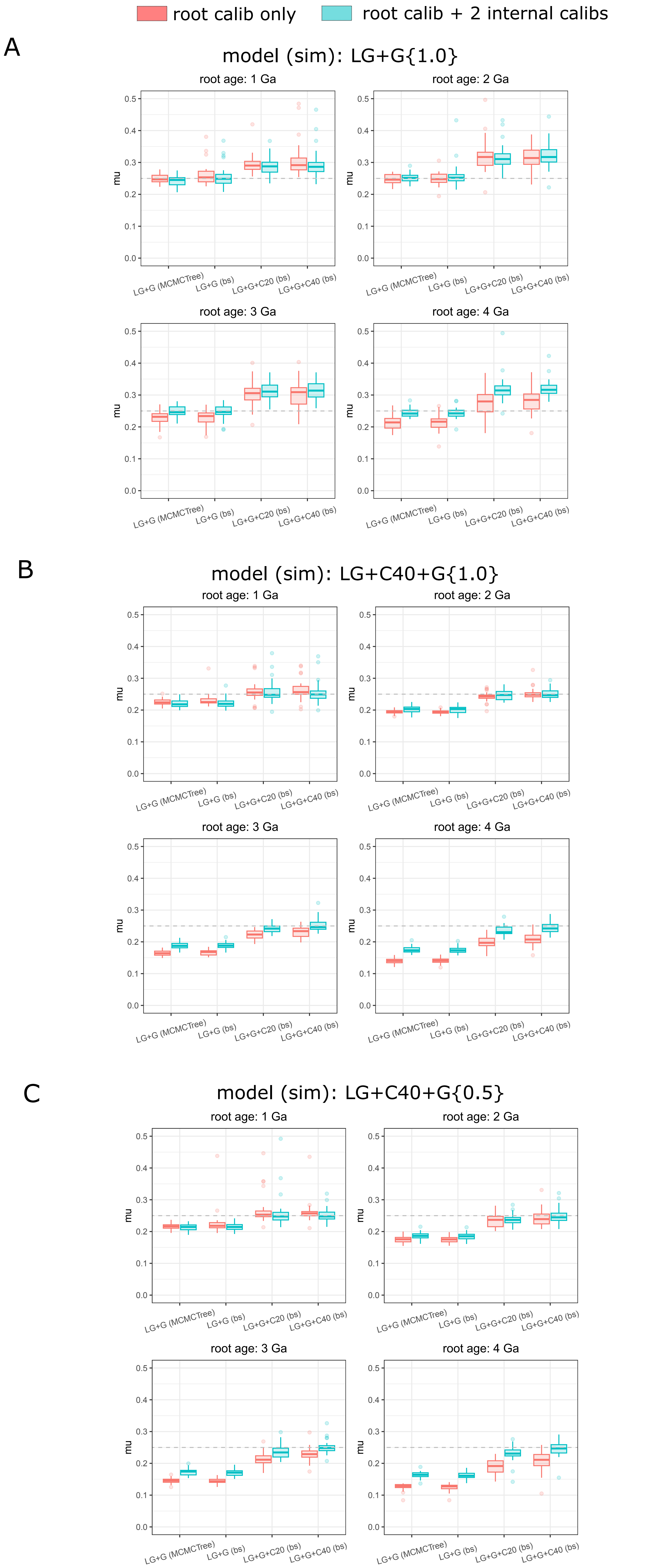

Figure S11. Assessing the uncertainty in the posterior dates with additional alternative settings in molecular clock analysis (Data S3). (A-C) Divergence times of bacteria estimated by alternative schemes (y-axis) versus using the focal scheme used in the main molecular clock analysis (x-axis). (A) Different root maximum ages as the prior, instead of a soft maximum of 4500 Ma used in the focal scheme. RootMax5000: soft maximum of 5000 Ma; RootMax5500: soft maximum of 5500 Ma; RootMax6000: soft maximum of 6000 Ma; rootMaxHard: hard maximum of 4500 Ma. (B) Different internal calibrations and settings of MCMCTree. EUK3: times of eukaryotes estimated using the AR model; EUK4: times of eukaryotes estimated with soft lower bounds; EUK5: the calibrations of nodes 31, 45, and 46 that overlap with their parent nodes and that may cause "truncation effect" are kept in the second step of the sequential mol ecular clock analysis (see Note S1.2.2); CyanoFossilCauchy: the two cyanobacteria nodes with fossils (total -group Nostocales and total-group Pleurocapsales) calibrated by a truncated Cauchy distribution; CyanoFossilSoft: the two cyanobacteria fossils with soft lower bounds; CyanoNos1200: lower bound of total-group Nostocales set as 1200 Ma based on a conservative assessment of the fossil record; CyanoNos2000: lower bound of total-group Nostocales set as 2000 Ma based on its earliest possible fossil records; BiomarkerSoft: the two nodes (total-group Chromatiaceae and total-group Chlorobi) calibrated by the biomarker with a soft lower bound; BiomarkerCauchy: the two nodes calibrated by the biomarker with a truncated Cauchy distribution; GOE_soft: total -group oxygenic cyanobacteria with soft lower bound based on GOE (2320 Ma); noBacCalib: no bacterial calibrations used; bd: the parameters for the birth -death (BD) process set as birth rate = 0.4 and death rate = 0.2 lineages/100 Ma according to (Scholl and Wiens 2016) instead of a flat prior. rate_mu: the prior of the mean rate based on empirical estimate; rate_sigma2: the prior on the variance of branch -wise log-transformed rate set as 1.0 to inform very large among-branch rate variation. 2-partition: two partitions, instead of a single partition of sequence alignment, is used. (C) Changes in the posterior dates of the crown group of selected bacterial groups shown as 95% HPD interval estimated with different tree topologies. euk_topo 1-4: different tree topologies within the eukaryotes (Fig. S8D); mito-rick: mitochondria are placed as the sister to Rickettsiales based on earlier studies (Andersson et al. 1998; Wang and Wu 2015) (Fig. S8E).

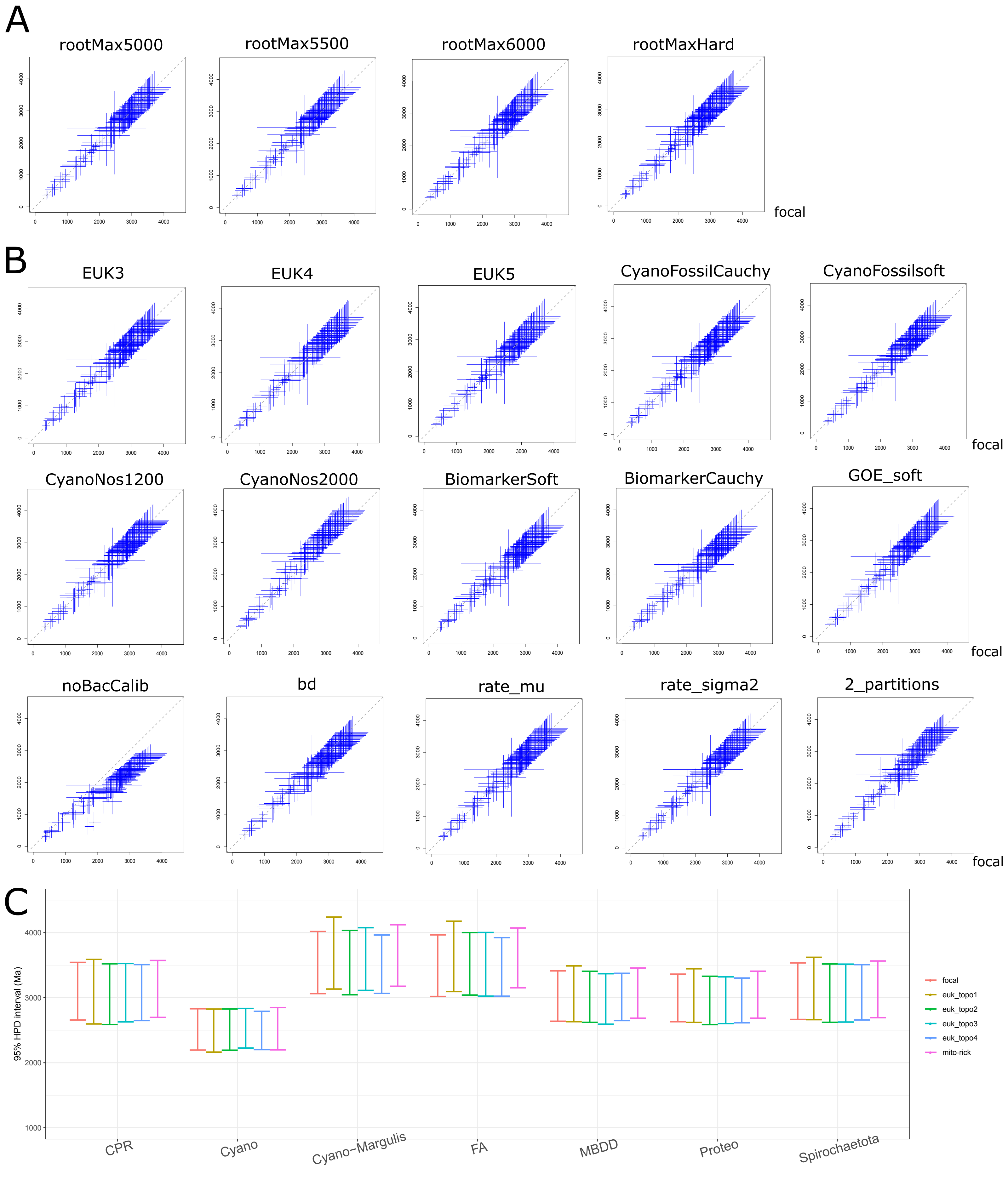

Figure S12. Comparison of the timetrees obtained with selected dating schemes. Green, red, and orange dashed lines indicate the estimated ages of the crown groups of oxygenic cyanobacteria, Proteobacteria, and mitochondria, respectively. Boxes in red adjacent to tips of the tree indicate those selected from "symbiont" clades as RTCs, as used in most of the analyses. Boxes in blue used in the scheme more_RTCs indicate those additionally included RTCs to examine if more RTCs will lead to a decrease of time. 2 -partition: two partitions, instead of a single partition of sequence alignment, is used. Euk1: the minimum time bound of crown-group red algae alternatively set as 1.6 Ga based on Rafatazmia. noCyanoFossil: the two cyanobacteria minimum time bounds (total-group Nostocales and Pleurocapsales) removed. Rick-Mito: mitochondria placed as the sister to Rickettsiales. all_genes: molecular dating performed on all 32 mitochondria-originated genes conserved across bacteria. more_RTCs: additional 13 symbionts included. Secondary: different from the 148 -genome set, a secondary independent and expanded sampling of bacterial genomes where 40% of organisms are from the CPR clade based on the "secondary dataset" from the study (Coleman et al. 2021); 314 genomes (285 bacteria and 29 eukaryotes) are included (Data S4).

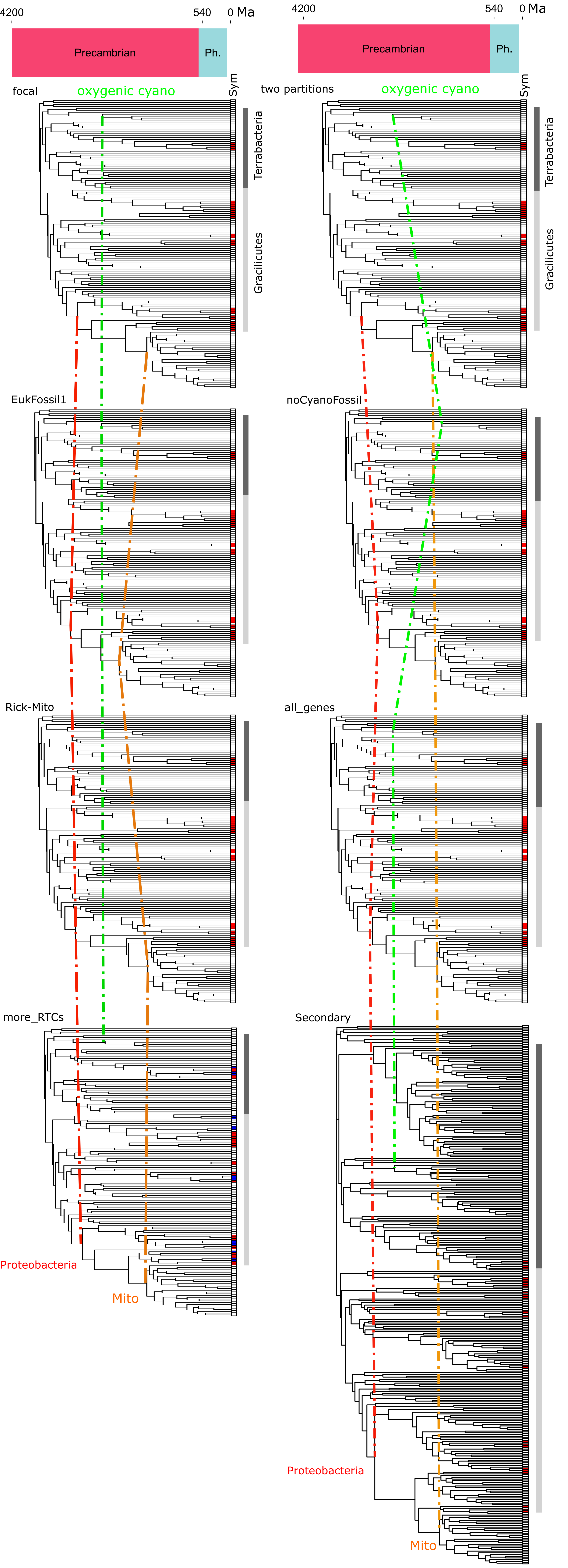

Figure S13. The posterior probability of the preferred model, i.e. the auto-correlated rate model. Two competing models, the autocorrelated rates (AR) and the independent rates (IR) model, are compared using MCMCTree's exact likelihood and mcmc3r's stepping-stones integrator. The histogram of the probability of the AR model given the alignment D, Pr(AR|D), is displayed. For molecular clock of the bacterial tree (148-genome focal dataset), due to the large computational burden, 20 and 40 randomly selected organisms are used. The full set of the 32 mitochondria-originated genes conserved across bacteria are analysed. For the eukaryote timetree, 320 orthologs and 29 species are analysed.

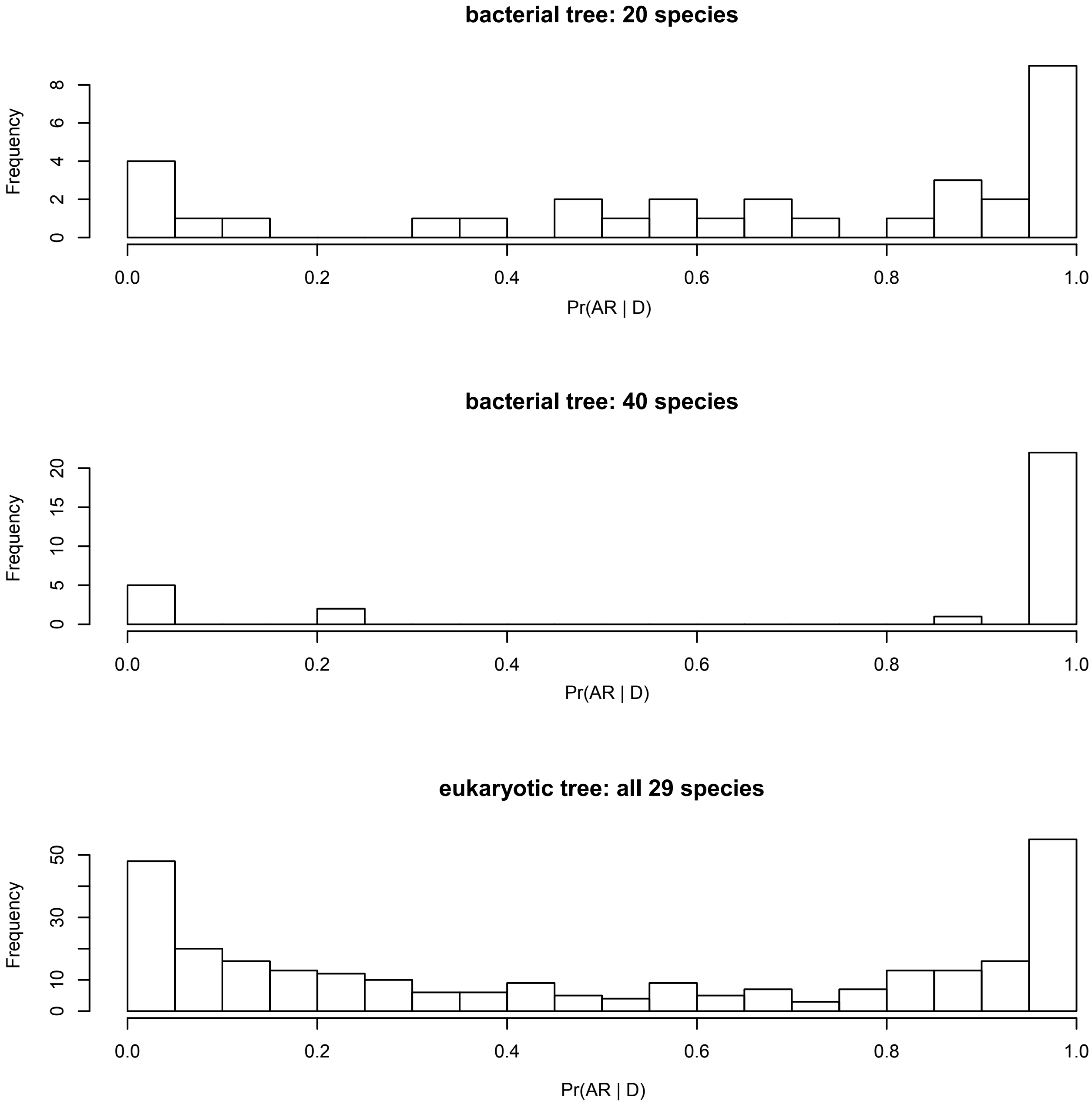

Figure S14. Assessing the uncertainty in Bayesian molecular clock analysis with alignment of two partitions. This is similar to what is shown in Fig. 3 except that in Fig. 3 results are obtained based on molecular analysis with alignment of a single partition. (A) Comparison of the posterior mean ages (lower triangle; unit: Ma) and rates (upper triangle; unit: number of substitutions per site per Ga) of bacteria estimated with different strategies. Strategy 1 (traditional strategy used in prior studies): five bacterial calibrations with hard minimum bounds and a soft maximum bound <4.5 Ga at LBCA, no eukaryote timing information, substitution model LG+G, no RTCs. Strategy 2: bacterial calibrations, eukaryote times by sequential molecular dating, substitution model LG+G, no RTCs. Strategy 3: bacterial calibrations, eukaryote times by sequential molecular dating, substitution model LG+G+C60, no RTCs. Focal: bacterial calibrations, eukaryote times by sequential molecular dating, substitution model LG+G+C60, with RTCs. (B) Posterior mean ages of the selected clades calculated under root maximum prior ages from 4500 to 6000 Ma with the focal strategy (dashed line) and Strategy 1 (solid line). (C) Divergence times of bacteria estimated by alternative schemes (y-axis; see Data S3) versus the one in the focal strategy (x-axis). The blue bars denote the 95% HPD. Euk1: the minimum time bound of crown-group red algae alternatively set as 1.6 Ga based on Rafatazmia; Euk2: the minimum time bound of animal crown group alternatively set as 0.89 Ga (Note S3.2); noCyanoFossil: the two cyanobacteria minimum time bounds (total-group Nostocales and Pleurocapsales) removed; noBiomarker: the two bacterial biomarker minimum time bounds (total-group Chromatiaceae and Chlorobi) removed; noGOE: the minimum time bound based on GOE removed. Joint_prob: rejection sampling based on calculating the joint probability in Eq. (2). Alt_lifestyle: ASR inferred with alternative classification of lifestyles of modern symbionts (Note S3.1). OTU97: ASR performed with OTU at a cut-off of 97% sequence identity. ASR_ML: maximum likelihood algorithm instead of MCMC used in ASR. IR: the independent rate (IR) model is used instead of the auto-correlated rate (AR) model.

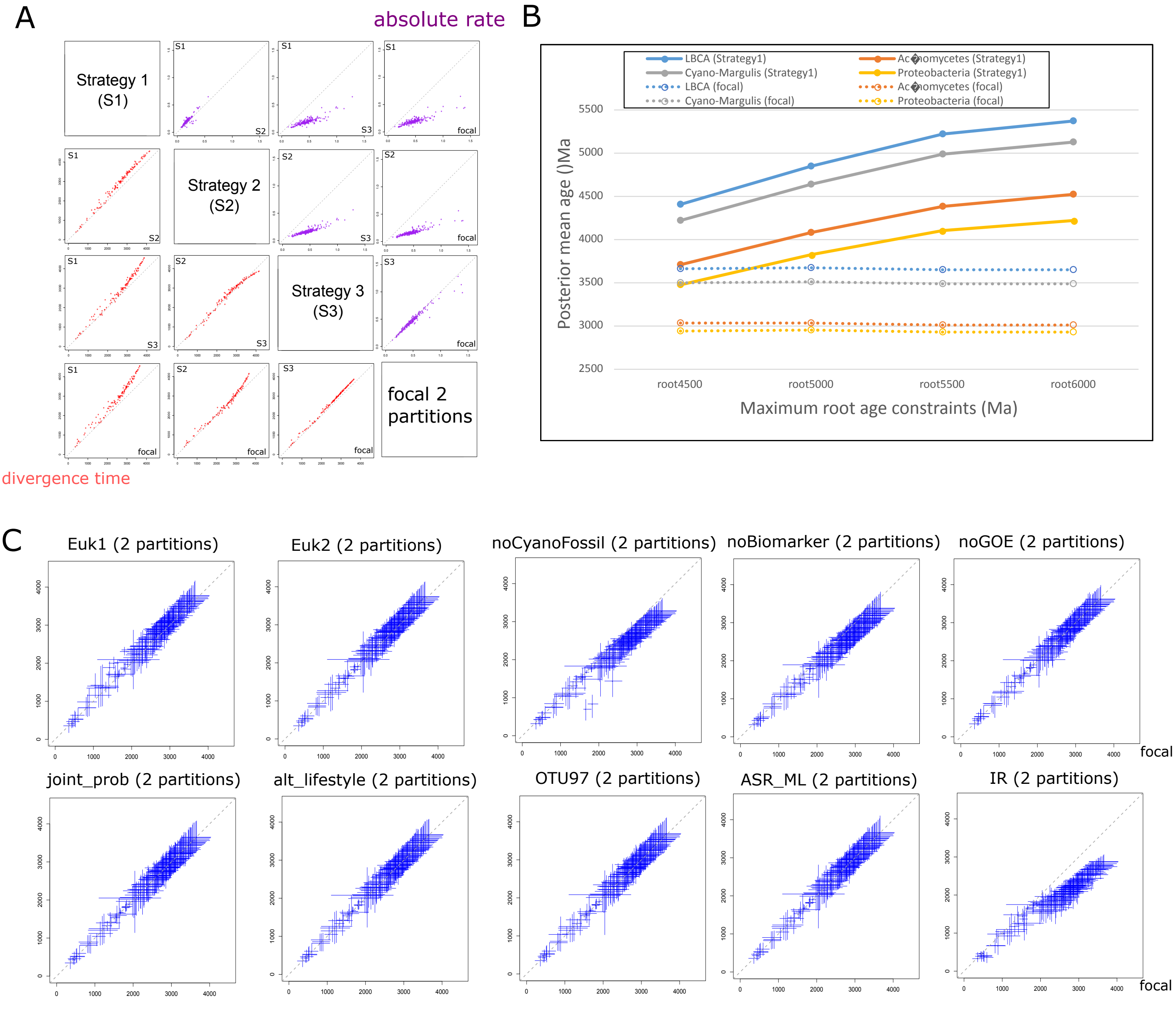

Figure S15. Divergence times of the taxonomically high-resolution bacterial tree with 8898 tips estimated using RelTime. The calibrations are indicated by black dots on the internal nodes. The time unit is 100 million years. (A) Posterior dates calculated from the 148-genome molecular clock analysis are fitted by uniform distribution. (B) Posterior dates calculated from the 148-genome molecular clock analysis are fitted by normal distribution but with calibrations where more than 10% overlap in the specified distributions between the parent and child nodes removed. (C) Posterior dates calculated from the 148-genome molecular clock analysis are fitted by uniform distribution but with calibrations where more than 10% overlap in the specified distributions between the parent and child nodes removed. (D) Schematic graph showing how overlapped calibrations are iteratively removed to keep a maximum number of calibrations without significant overlap. In the displayed tree, the calibration of node 7 is removed as it "significantly" overlaps with that of its child nodes while all the other six calibrations are kept.

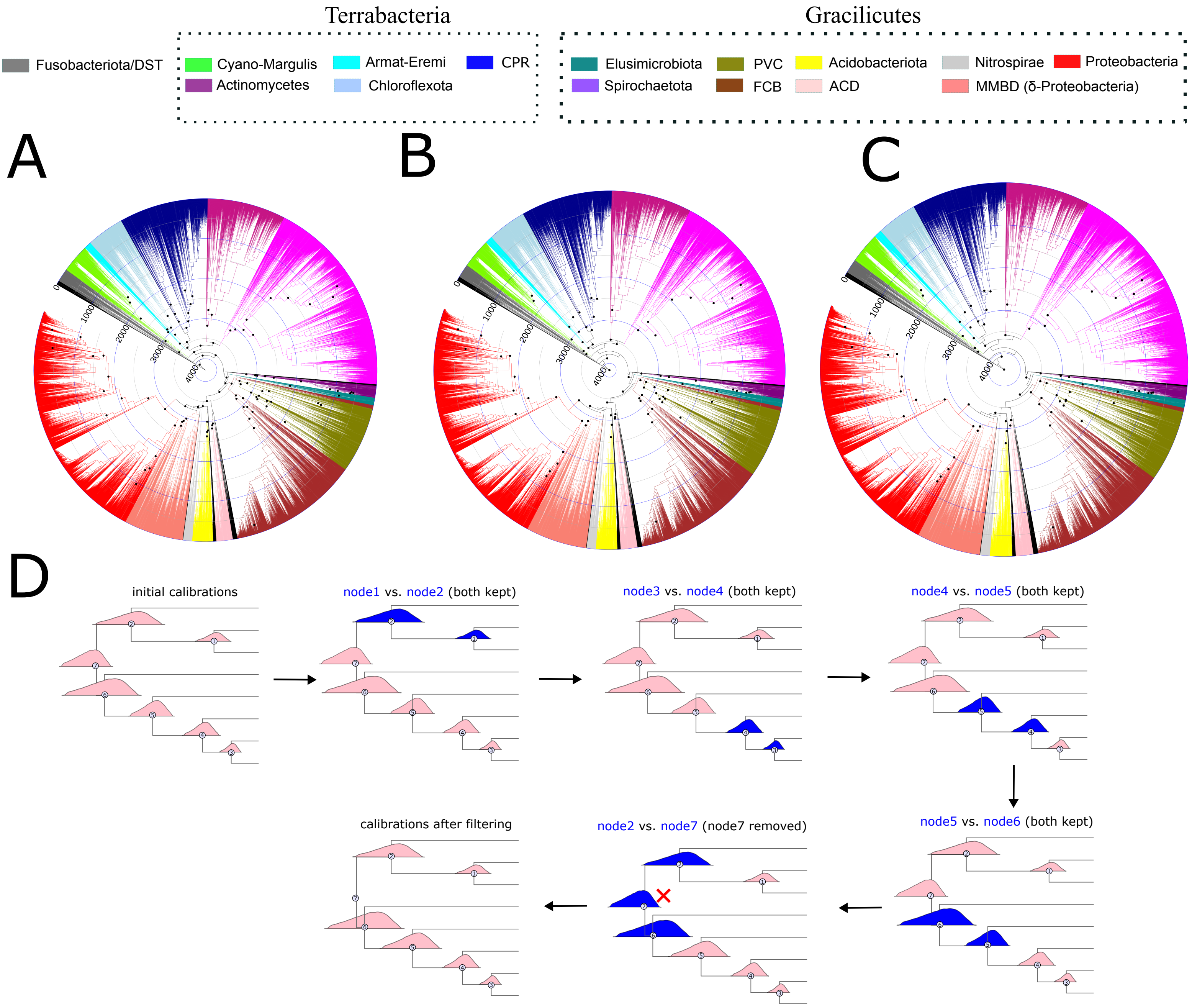

**Table S1A.** Genome sources of bacteria used in the focal dataset of molecular dating (148-genome dataset). Shaded lines indicate genomes used in the study (Coleman et al. 2021).

| **Taxonomy** | **Organism** | **Assembly** | **RTC?** |
|---|---|---|---|
| Cyanobacteria | Gloeobacter violaceus PCC 7421 | GCF_000011385 | |
| Cyanobacteria | Nostoc sp. PCC 7107 | GCF_000316625 | |
| Cyanobacteria | Oscillatoria sp. PCC 10802 | GCF_000332335 | |
| Cyanobacteria | Xenococcus sp. PCC 7305 | GCF_000332055 | |
| Firmicutes | Bacillus subtilis | GCA_000009045 | |
| Proteobacteria | Escherichia coli str K-12 substr W3110 | GCF_000010245 | |
| Proteobacteria | Neisseria gonorrhoeae FA 1090 | GCF_000006845 | |
| Proteobacteria | Rhodocyclales bacterium GWA2 65 20 | GCA_001828925 | |
| Proteobacteria | Nitrosococcus oceani ATCC 19707 | GCF_000012805 | |
| Proteobacteria | Thiocapsa marina 5811 | GCF_000223985 | |
| Chlamydiota | Candidatus Clavichlamydia salmonicola | GCA_015356765 | Y |
| Chlamydiota | Candidatus Rhabdochlamydia sp. W815 | GCA_018642185 | Y |
| Elusimicrobiota | Candidatus Endomicrobium trichonymphae | GCA_001730735 | Y |
| Elusimicrobiota | Endomicrobiaceae bacterium | GCA_016292265 | Y |
| Firmicutes | Catharanthus roseus aster yellows phytoplasma | GCA_004214875 | Y |
| Firmicutes | Mycoplasma gypis | GCA_016924775 | Y |
| Firmicutes | Mycoplasma penetrans HF-2 | GCA_000011225 | Y |
| Firmicutes | Texas Phoenix palm phytoplasma | GCA_005774685 | Y |
| Proteobacteria | Blattabacterium clevelandi | GCA_003268615 | Y |
| Proteobacteria | Blattabacterium sp. Nauphoeta cinerea | GCA_000471965 | Y |
| Proteobacteria | Buchnera aphidicola Cinara tujafilina | GCA_000217635 | Y |
| Proteobacteria | Buchnera aphidicola Tuberolachnus salignus | GCA_900016785 | Y |
| Proteobacteria | Caedimonas varicaedens | GCA_001192655 | Y |
| Proteobacteria | Candidatus Berkiella aquae | GCA_001431295 | Y |
| Proteobacteria | Candidatus Finniella inopinata | GCA_004210305 | Y |
| Proteobacteria | Coxiella burnetii str Schperling | GCA_002634065 | Y |
| Proteobacteria | Coxiella-like endosymbiont | GCA_002871095 | Y |
| Proteobacteria | Neorickettsia sp. 179522 | GCA_001593535 | Y |
| Proteobacteria | Rickettsiales bacterium | GCA_016780625 | Y |
| Proteobacteria | Wolbachia endosymbiont of Cylisticus convexus | GCA_003344345 | Y |
| Spirochaetota | Breznakiella homolactica | GCA_016616095 | Y |
| Spirochaetota | Spirochaetes bacterium | GCA_016285315 | Y |
| Spirochaetota | Treponema brennaborense DSM 12168 | GCA_000212415 | Y |
| Spirochaetota | Treponema denticola F0402 | GCA_000191825 | Y |
| Spirochaetota | Treponema primitia ZAS-1 | GCA_000297095 | Y |
| Spirochaetota | Treponema sp. OMZ 806 | GCA_018336815 | Y |
| Acidobacteriota | Acidobacteria bacterium Mor1 | GCA_001664505 | |
| Acidobacteriota | bacterium HR11 | GCA_002898535 | |
| Acidobacteriota | Candidatus Aminicenantes bacterium UBA6072 | GCA_002436105 | |
| Acidobacteriota | Geothrix fermentans DSM 14018 | GCF_000428885 | |
| Acidobacteriota | Holophagales bacterium UBA2201 | GCA_002327305 | |
| Actinobacteriota | Actinobacteria bacterium UBA2594 | GCA_002339355 | |

| | | | |
|---|---|---|---|
| Actinobacteriota | Nocardiopsis sp. TSRI0078 | GCF_001905145 | |
| Actinobacteriota | Olegusella massiliensis | GCF_900078545 | |
| Actinobacteriota | Rubrobacter radiotolerans DSM 5868 | GCF_900175965 | |
| Aquificota | Thermocrinis minervae | GCF_900142435 | |
| Armatimonadota | Abditibacterium utsteinense | GCF_002973605 | |
| Armatimonadota | Armatimonadetes bacterium UBA1398 | GCA_002305165 | |
| Armatimonadota | Chthonomonas calidirosea T49 | GCF_000427095 | |
| Bacteroidota | Bacteroidetes bacterium CG12 big fil rev 8 21 14 0 65 60 17 | GCA_002787815 | |
| Bacteroidota | Chlorobium limicola DSM 245 | GCF_000020465 | |
| Bdellovibrionota | Bdellovibrionales bacterium CG10 big fil rev 8 21 14 0 10 45 34 | GCA_002778785 | |
| Bdellovibrionota | Deltaproteobacteria bacterium CG11 big fil rev 8 21 14 0 20 45 16 | GCA_002787535 | |
| Campylobacterota | Hippea jasoniae | GCF_000744435 | |
| Chlamydiota | Parachlamydia acanthamoebae | GCF_000875975 | Y |
| Chloroflexota | Chloroflexi bacterium GCA 003171065.1 | GCA_003171065 | |
| Chloroflexota | SAR202 cluster bacterium bin16 | GCA_002238425 | |
| Chloroflexota | Thermogemmatispora onikobensis | GCF_001748285 | |
| Cloacimonadota | Cloacimonetes bacterium UBA3541 | GCA_002376725 | |
| Cyanobacteria | Candidatus Sericytochromatia bacterium S15B-MN24 CBMW 12 | GCA_002083825 | |
| Deferribacterota | Deferribacter desulfuricans SSM1 | GCF_000010985 | |
| Deinococcota | Marinithermus hydrothermalis DSM 14884 | GCF_000195335 | |
| Dependentiae | Candidatus Dependentiae bacterium | GCA_002401785 | |
| Desulfobacterota | Desulfuromonas soudanensis | GCF_001278055 | |
| Dormibacterota | Candidatus Dormibacteraeota bacterium | GCA_003139695 | |
| Elusimicrobiota | Elusimicrobia bacterium CG1 02 37 114 | GCA_001871125 | Y |
| Eremiobacterota | bacterium UBP9 UBA4705 | GCA_002407045 | |
| Eremiobacterota | Candidatus Eremiobacteraeota bacterium | GCA_003158175 | |
| Fibrobacterota | Candidatus Raymondbacteria bacterium RIFOXYA2 FULL 49 16 | GCA_001789205 | |
| Fibrobacterota | Chitinivibrio alkaliphilus ACht1 | GCF_000474745 | |
| Fibrobacterota | Fibrobacter sp. UWP2 | GCF_900141705 | |
| Firmicutes | Marininema mesophilum | GCF_900106775 | |
| Firmicutes | Tepidanaerobacter syntrophicus JL | GCF_001485475 | |
| Firmicutes | Ammonifex degensii KC4 | GCF_000024605 | |
| Firmicutes | Natranaerobius trueperi | GCF_002216005 | |
| Firmicutes | Acetohalobium arabaticum DSM 5501 | GCF_000144695 | |
| Firmicutes | Limnochorda pilosa | GCF_001544015 | |
| Fusobacteriota | Leptotrichia sp. oral taxon 847 | GCF_001553645 | |
| Gemmatimonadota | Gemmatimonadetes bacterium | GCA_003223395 | |
| Margulisbacteria | candidate division WOR-1 bacterium RIFOXYB2 FULL 45 9 | GCA_001771575 | |
| Margulisbacteria | Candidatus Margulisbacteria bacterium | GCA_003242895 | |
| Marinisomatota | Candidatus Marinimicrobia bacterium CG1 02 48 14 | GCA_001872685 | |
| Myxococcota | Bradymonas sediminis | GCF_003258315 | |
| Myxococcota | Deltaproteobacteria bacterium UBA6601 | GCA_002433485 | |
| Nitrospirota | Nitrospirae bacterium | GCA_002737345 | |
| Omnitrophota | Omnitrophica bacterium RIFCSPLOWO2 12 FULL 44 17 | GCA_001804285 | |

| | | |
|---|---|---|
| Patescibacteria | candidate division CPR3 bacterium 4484 211 | GCA_002084955 |
| Patescibacteria | candidate division WWE3 bacterium GW2011 GWC2 41 23 | GCA_000996895 |
| Patescibacteria | Candidatus Andersenbacteria bacterium RIFCSPHIGHO2 02 FULL 45 11 | GCA_001817055 |
| Patescibacteria | Candidatus Berkelbacteria bacterium CG10 big fil rev 8 21 14 0 10 41 12 | GCA_002778735 |
| Patescibacteria | Candidatus Dojkabacteria bacterium HGW-Dojkabacteria-1 | GCA_002840365 |
| Patescibacteria | Candidatus Giovannonibacteria bacterium RIFCSPLOWO2 12 FULL 44 15 | GCA_001778905 |
| Patescibacteria | Candidatus Peregrinibacteria bacterium GW2011 GWF2 39 17 | GCA_000993615 |
| Patescibacteria | Candidatus Roizmanbacteria bacterium RIFCSPHIGHO2 12 FULL 37 23 | GCA_001788095 |
| Patescibacteria | Candidatus Saccharibacteria bacterium UBA1788 | GCA_002315165 |
| Patescibacteria | Candidatus Uhrbacteria bacterium RIFOXYA2 FULL 40 9 | GCA_001791615 |
| Patescibacteria | Patescibacteria group bacterium UBA2591 | GCA_002339755 |
| Planctomycetota | bacterium- GCA_003245715 | GCA_003245715 |
| Planctomycetota | bacterium HR36 | GCA_002898995 |
| Planctomycetota | Phycisphaerales bacterium Planc 01 | GCA_001603075 |
| Planctomycetota | Phycisphaerales bacterium UBA5793 | GCA_002418285 |
| Planctomycetota | Planctomycetes bacterium GCA 002746535.1 | GCA_002746535 |
| Planctomycetota | Planctomycetes bacterium GCA 002748355.1 | GCA_002748355 |
| Planctomycetota | Planctomycetes bacterium | GCA_003136555 |
| Planctomycetota | Planctomycetes bacterium RIFCSPLOWO2 12 FULL 50 35 | GCA_001828605 |
| Planctomycetota | Planctomycetes bacterium UBA1662 | GCA_002320775 |
| Proteobacteria | Magnetococcales bacterium DC0425bin3 | GCA_002753665 |
| Proteobacteria | Mariprofundus micogutta | GCF_001895085 |
| Spirochaetota | Borrelia hispanica CRI | GCF_000500065 |
| Spirochaetota | Brachyspira hyodysenteriae ATCC 27164 | GCF_001676785 |
| Spirochaetota | Leptospira levettii | GCF_002812165 |
| Spirochaetota | Spirochaetes bacterium GWB1 27 13 | GCA_001829125 |
| Spirochaetota | Spirochaetes bacterium GWF1 51 8 | GCA_001829415 |
| Spirochaetota | Spirochaetia bacterium UBA6919 | GCA_002450905 |
| Synergistota | bacterium GBS-1 | GCA_001443005 |
| Thermotogota | Pseudothermotoga thermarum DSM 5069 | GCF_000217815 |
| Verrucomicrobiota | Kiritimatiellales bacterium | GCA_003230915 |
| Verrucomicrobiota | Lentisphaeria bacterium UBA7640 | GCA_002483765 |
| Verrucomicrobiota | Verrucomicrobiales bacterium | GCA_002715965 |

**Table S1B.** Genome sources of eukaryotes.

| Taxonomy | Organism | Nuclear genome | Mitogenome |
|---|---|---|---|
| Metazoa | Homo sapiens | Ensembl release 96 (Howe et al. 2020) | MitoCOGs |
| Metazoa | Gallus gallus | Ensembl release 96 (Howe et al. 2020) | MitoCOGs |
| Metazoa | Branchiostoma floridae | UniProt (Bateman 2019) | MitoCOGs |
| Metazoa | Amphimedon queenslandica | Ensembl Metazoa release 46 (Howe et al. 2020) | MitoCOGs |
| Metazoa | Caenorhabditis elegans | Ensembl release 96 (Howe et al. 2020) | MitoCOGs |
| Metazoa | Drosophila melanogaster | Ensembl release 96 (Howe et al. 2020) | MitoCOGs |
| Metazoa | Daphnia pulex | ACJG00000000 (Colbourne et al. 2011) | NC_000844 |
| Choanoflagellata | Salpingoeca rosettas | Ensembl Protists release 52 (Howe et al. 2020) | |
| Fungi | Candida albicans | UniProt (Bateman 2019) | MitoCOGs |
| Fungi | Ustilago maydis | Ensembl Fungi release 46 (Howe et al. 2020) | MitoCOGs |
| Fungi | Pleurotus ostreatus | UniProt (Bateman 2019) | MitoCOGs |
| Fungi | Spizellomyces punctatus | UniProt (Bateman 2019) | MitoCOGs |
| Amoebozoa | Acanthamoeba castellanii | Ensembl Protists release 43 (Howe et al. 2020) | MitoCOGs |
| Amoebozoa | Dictyostelium discoideum | Ensembl Protists release 43 (Howe et al. 2020) | MitoCOGs |
| Amoebozoa | Polysphondylium pallidum | dictyBase (Basu et al. 2013) | MitoCOGs |
| Archaeplastida | Arabidopsis thaliana | PLAZA 4.0 (Van Bel et al. 2018) | MitoCOGs |
| Archaeplastida | Oryza sativa | PLAZA 4.0 (Van Bel et al. 2018) | MitoCOGs |
| Archaeplastida | Physcomitrella patens | PLAZA 4.0 (Van Bel et al. 2018) | MitoCOGs |
| Archaeplastida | Ostreococcus tauri | PLAZA 2.0 (Proost et al. 2009) | MitoCOGs |
| Archaeplastida | Chondrus crispus | Ensembl Plants release 46 (Howe et al. 2020) | MitoCOGs |
| Archaeplastida | Porphyra umbilicalis | UniProt (Bateman 2019) | NC_018544 |
| Archaeplastida | Cyanidioschyzon merolae | Ensembl Plants release 43 (Howe et al. 2020) | MitoCOGs |
| Archaeplastida | Cyanophora paradoxa | http://cyanophora.rutgers.edu/cyanophora/ | MitoCOGs |
| Discoba | Andalucia godoyi | https://megasun.bch.umontreal.ca/Andalucia_godoyi/ (Gray et al. 2020) | MitoCOGs |
| SAR | Symbiodinium minutum | https://marinegenomics.oist.jp/ v1.2 (Shoguchi et al. 2013) | |
| SAR | Paramecium tetraurelia | Ensembl Protists release 45 (Howe et al. 2020) | NC_001324 |
| SAR | Oxytricha trifallax | OxyDB (http://oxy.ciliate.org/) | JN383843 |
| SAR | Phytophthora infestans | Ensembl Protists release 46 (Howe et al. 2020) | MitoCOGs |
| SAR | Thalassiosira pseudonana | Ensembl Protists release 46 (Howe et al. 2020) | MitoCOGs |

**Table S2.** Determining the root placement of the bacterial tree based on the 265 genomes and 60 orthologs used in (Coleman et al. 2021) by IQ-Tree's non-reversible model.

(A) Tree reconstruction using time reversible model.

| Scheme | Partition | Model | Fixed topology? | start tree | AIC[a] | BIC[b] | AICc[c] |
|---|---|---|---|---|---|---|---|
| REV_C20 | - | REV (mixture): LG+G+C20 | - | default | 10305504 | 10305536 | 10309810 |
| REV_C40 | - | REV (mixture): LG+G+C40 | - | default | 10218128 | 10218160 | 10222434 |
| REV_C60 | - | REV (mixture): LG+G+C60 | - | default | 10180630 | 10180661 | 10184936 |
| REV_par | ModelFinder [d] | REV (non-mixture) [e] | - | default | 10849748 | 10849805 | 10855503 |

(B) Tree reconstruction using time non-reversible (NONREV) model with those constructed by reversible models as either the fixed topology or start tree.

| Scheme | Partition | Model | Fixed topology? | start tree | AIC[a] | BIC[b] | AICc[c] |
|---|---|---|---|---|---|---|---|
| NONREV_par_fixed-1 | ModelFinder | NONREV (non-mixture) [f] | REV_C20 | REV_C20 | 10801857 | 10801965 | 10809713 |
| NONREV_par_fixed-2 | ModelFinder | NONREV (non-mixture) | REV_C40 | REV_C40 | 10801854 | 10801963 | 10809711 |
| NONREV_par_fixed-3 | ModelFinder | NONREV (non-mixture) | REV_C60 | REV_C60 | 10801518 | 10801626 | 10809375 |
| NONREV_fixed-1 | - | NONREV (non-mixture) | REV_C20 | REV_C20 | 10807603 | 10807692 | 10814758 |
| NONREV_fixed-2 | - | NONREV (non-mixture) | REV_C40 | REV_C40 | 10807811 | 10807900 | 10814966 |
| NONREV_fixed-3 | - | NONREV (non-mixture) | REV_C60 | REV_C60 | 10807706 | 10807795 | 10814861 |
| NONREV_par-1 | ModelFinder | NONREV (non-mixture) | - | REV_C20 | 10800672 | 10800781 | 10808529 |
| NONREV_par-2 | ModelFinder | NONREV (non-mixture) | - | REV_C40 | 10800677 | 10800785 | 10808533 |
| NONREV_par-3 | ModelFinder | NONREV (non-mixture) | - | REV_C60 | 10800724 | 10800833 | 10808581 |
| NONREV-1 | - | NONREV (non-mixture) | - | REV_C20 | 11643798 | 11643887 | 11650938 |
| NONREV-2 | - | NONREV (non-mixture) | - | REV_C40 | 11643746 | 11643835 | 11650886 |
| NONREV-3 | - | NONREV (non-mixture) | - | REV_C60 | 11643823 | 11643912 | 11650963 |

[a] AIC: Akaike information criterion. A lower value indicates a better model fit (same below for BIC and AICc).

[b] BIC: Bayesian information criterion

[c] AICc: a version of AIC that has a correction for small sample sizes

[d] Partitioning determined by ModelFinder implemented in IQ-Tree.

[e] For model selection of each partition, the settings "-m MFP+MERGE -mset LG,WAG,JTT -mrate G,I,G+I,E" in IQ-Tree are used, hence only non-mixture model is considered. In other words, the mixture model Cxx is not considered.

[f] Note that NONREV is a non-mixture model (cannot be used together with Cxx model).

**Table S3.** List of the 32 mitochondria-originated genes conserved across the bacterial tree. The first 19 genes listed in the table are used in the main analysis.

| Mito-originated genes in Gomez et al. 2022 | Gene names in commonly used sets of genes conserved across bacteria | | | Final 19-gene set? | Reason for exclusion [a] |
|---|---|---|---|---|---|
| | **Battistuzzi2009** | **bac120** | **Coleman2021** | | |
| ATP5F1A | | atpD | K02112 | Y | |
| ATP5F1B | | atpD | K02112 | Y | |
| ATP5F1C | | ATPsyn_F1gamma | | Y | |
| CLPP | | | K01358 | Y | |
| ENGA | | GTPase_EngA | | Y | |
| FTSZ | | ftsZ | | Y | |
| GRPEL1 | | GrpE | | Y | |
| HSPA9 | | prok_dnaK | | Y | |
| HSPD1 | | | K04077 | Y | |
| MARS2 | | metG | | Y | |
| METTL15 | | TIGR00006 | K03438 | Y | |
| MRPL1 | RplA | rplA_bact | K02863 | Y | |
| MRPL4 | | rplD_bact | | Y | |
| MRPS9 | RpsI | Ribosomal_S9 | K02996 | Y | |
| PRFA | | | K02835 | Y | |
| RPL6 | RplF | L6_bact | K02933 | Y | |
| RSAD1 | | hemN_rel | | Y | |
| TFB1M | | ksgA | | Y | |
| TILS | | lysidine_TilS_N | | Y | |
| CLPX | | clpX | | N | MB |
| GUF1 | | | K02355 | N | UP |
| MRPL11 | RplK | L11_bact | K02867 | N | NM |
| MRPL16 | RplP | rplP_bact | K02878 | N | MB |
| MRPL2 | RplB | rplB_bact | K02886 | N | MB |
| MRPL20 | | rplT_bact | K02887 | N | MB |
| MRPL3 | RplC | L3_bact | K02906 | N | MB |
| MRPS2 | RpsB | rpsB_bact | K02967 | Y | MB |
| RPOC | RpoC | rpoC_TIGR | K03046 | N | UP |
| RPS19 | | | K02965 | N | MB |
| RPS4 | RpsD | rpsD_bact | | N | MB |
| RPS8 | | Ribosomal_S8 | K02994 | N | MB |
| TUFM | TufB | | K02358 | N | NM |

[a]: Abbreviations of the reason to exclude the gene, MB: mitochondria basal at the tree when midpoint rooted (see Data Availability for individual gene trees with alternative rooting methods); NM: mitochondria and α-Proteobacteria not monophyly (they together do not form a monophyletic group); UP: unresolved paralogy.

**Table S4.** Comparison of the posterior and prior dates (mean and 95% HPD intervals) using the focal scheme (either one or two partition) on the 148-genome dataset. Prior dates are determined by the probability distribution specified by user for those with a calibration and a birth-death process for uncalibrated nodes, while posteriors are informed by both the priors and the data. Dates are given in the unit of million years (Ma).

(A) focal (single partition)

| | Effective priors | | | Posterior | | |
|---|---|---|---|---|---|---|
| Clade | mean | lower bound | upper bound | mean | lower bound | upper bound |
| ACD | 2908.08 | 1031.86 | 4351.91 | 3026.43 | 2624.71 | 3436.44 |
| Actinomycetes | 2512.69 | 829.083 | 4312.58 | 3065.92 | 2630.02 | 3551.96 |
| Armati-Eremi | 2824.64 | 1026.45 | 4352.79 | 3382.81 | 2947.26 | 3883.26 |
| Chloroflexota | 2327.72 | 823.681 | 4282.1 | 3008.35 | 2586.19 | 3489.19 |
| CPR | 3102.59 | 1358.25 | 4331.22 | 3070.11 | 2656.77 | 3543.42 |
| Cyano-Margulis | 3889.67 | 3078.21 | 4540.13 | 3548.88 | 3062.91 | 4018.45 |
| Cyanobacteria | 3436.95 | 2449.21 | 4365.11 | 2478.99 | 2194.96 | 2830.09 |
| Elusimicrobiota | 2473.21 | 599.206 | 4367.69 | 1763.58 | 1037.03 | 2447.22 |
| Mitochondria | 1549.06 | 1489.39 | 1607.89 | 1599.5 | 1557.63 | 1644.81 |
| FCB | 3728.91 | 2778.64 | 4523.51 | 3291.43 | 2834.57 | 3706.37 |
| Firmicutes | 3331 | 1468.07 | 4421.13 | 3197.55 | 2768.71 | 3673.54 |
| Fuso/DST | 2972.57 | 1014.1 | 4483.26 | 3628.57 | 3130.54 | 4138.29 |
| MBDD | 2705.21 | 1015.24 | 4295.37 | 3002.4 | 2638.55 | 3412.8 |
| Proteobacteria | 3613.06 | 2651.54 | 4511.42 | 2972.43 | 2630.64 | 3362.49 |
| PVC | 3688.56 | 2634.47 | 4524.82 | 3357.79 | 2900.39 | 3793.31 |
| root | 4139.27 | 3383.03 | 4646.32 | 3714.45 | 3200.49 | 4211.64 |
| Spirochaetota | 3774.66 | 2652.32 | 4554.94 | 3077.16 | 2666.51 | 3535.64 |

(B) focal (two partitions)

| | Effective priors | | | Posterior | | |
|---|---|---|---|---|---|---|
| Clade | mean | lower bound | upper bound | mean | lower bound | upper bound |
| ACD | 2920.89 | 1187.38 | 4506.79 | 3014.48 | 2689.4 | 3372.07 |
| Actinomycetes | 2517.41 | 829.946 | 4317.52 | 3100.24 | 2722.54 | 3482.09 |
| Armati-Eremi | 2824.99 | 1018.73 | 4328.18 | 3322.82 | 2960.89 | 3720.81 |
| Chloroflexota | 2314.15 | 827.916 | 4277.56 | 3054.24 | 2683.17 | 3430.15 |
| CPR | 3093.29 | 1354.33 | 4309.69 | 3016.12 | 2651.81 | 3396.51 |
| Cyano-Margulis | 3899.74 | 3071.03 | 4516.58 | 3581.94 | 3206.38 | 3983.85 |
| Cyanobacteria | 3442.14 | 2451.3 | 4357.05 | 2388.11 | 2149.04 | 2621.73 |
| Dependentiae | 2762.06 | 1006.19 | 4433.2 | 3336.37 | 2969.18 | 3706.29 |
| Elusimicrobiota | 2473.18 | 603.381 | 4365.99 | 2294.64 | 1627.38 | 2829.25 |
| Mitochondria | 1549.08 | 1490.14 | 1608.27 | 1632.36 | 1586.83 | 1676.78 |
| FCB | 3732.3 | 2767.01 | 4514.46 | 3303.08 | 2950.54 | 3678.66 |
| Firmicutes | 3323.4 | 1469.45 | 4423.59 | 3254.57 | 2902.23 | 3652.58 |
| Fuso/DST | 2953.87 | 1015.66 | 4482.91 | 3655.98 | 3258.92 | 4080.43 |
| MBDD | 2708.4 | 1024.21 | 4286.75 | 3038.35 | 2745.71 | 3369.29 |

| | | | | | | |
|---|---|---|---|---|---|---|
| Proteobacteria | 3615.3 | 2624.5 | 4506.05 | 2989.59 | 2700.99 | 3278.44 |
| PVC | 3690.05 | 2596.76 | 4518.97 | 3382.95 | 3024.64 | 3760.39 |
| root | 4153.19 | 3388.45 | 4653.51 | 3743.82 | 3355.61 | 4169.91 |
| Spirochaetota | 3779.63 | 2651.94 | 4564.78 | 3162.01 | 2811.99 | 3554.76 |

### Supplementary Note 1: Supplementary methods

#### 1.1 Taxon sampling

##### 1.1.1 Eukaryotic lineages

Twenty-nine eukaryotes were used to establish an evolutionary timeline of eukaryotes, which was further used in the Bayesian sequential molecular clock analysis. These taxa were mainly based on our previous study (Wang and Luo 2021) which cover major eukaryotic lineages. In addition, we added four to include lineages containing the host of important symbiotic bacteria lineages, thereby allowing applying relative time constraint (RTC)-based dating by comparing the posterior age between the host and symbiont.

Specifically, we included 25 out of the 27 taxa from the nuclear-encoded dataset used in Wang and Luo 2021. Note that two rhizarians originally included in Wang and Luo 2021, namely *Reticulomyxa filosa* and *Elphidium margaritaceum*, were discarded as they had only few of the final set of the mitochondria-originated genes used in molecular dating, likely because their mitochondrial genomes have not been sequenced (*Elphidium margaritaceum* even has only transcriptomic data). The remaining 25 eukaryotes consisted of eight from Archaeplastida (referred to as plants hereafter for simplicity), four animals, four fungi, three amoebae, four from the Stramenopiles-Alveolata-Rhizaria (SAR) supergroup, and one from Discoba. Specifically, the eight plants included two flowering plants (*Arabidopsis thaliana* and *Oryza sativa*), a bryophyte (*Physcomitrella patens*), a green alga (*Ostreococcus tauri*), three red algae (*Porphyra umbilicalis*, *Chondrus crispus* and *Cyanidioschyzon merolae*), and a glaucophyte (*Cyanophora paradoxa*). The four animals included two amniotes (*Homo sapiens* and *Gallus gallus*), a primitive chordate amphioxus (*Branchiostoma floridae*), and a sponge (the presumably earlies-split metazoan lineage; *Amphimedon queenslandica*). The four fungi comprised three from Dikarya, a subkingdom of fungi that in general produce dikaryons (*Ustilago maydis*, *Candida albicans*, and *Pleurotus ostreatus*), as well as an early-branching fungal lineage chytrid (*Spizellomyces punctatus*). The three amoebae were two social amoebae (*Dictyostelium discoideum* and *Polysphondylium pallidum*), and *Acanthamoeba castellanii* from Discosea (flattened amoebae moving as an entity). The four SAR lineages consisted of a dinoflagellate (protists where most members are characterized by two dissimilar flagella; *Symbiodinium minutum*), two ciliates [characterized by the presence of cilia; *Paramecium tetraurelia* and *Oxytricha trifallax*], and a diatom (*Thalassiosira pseudonana*). The only one from Discoba was the jakobid *Andalucia godoyi*, famous for having one of the most bacteria-like and most gene-rich mitochondrial genomes known to date.

The four eukaryotes added in the presented study included one from Choanoflagellates, and three from Ecdysozoa. Choanoflagellates are a group of unicellular and colonial flagellate eukaryotes that adapt to a free-living lifestyle. They have been long considered as the closest living relatives of animals (Metazoa) (Carr et al. 2008; Paps and Holland 2018). The choanoflagellate included in the present study is *Salpingoeca rosetta,* a rare marine eukaryote that consists of cells embedded in a jelly-like matrix and that shows a very primitive level of cell differentiation (Dayel et al. 2011). It was therefore used in the RTC-based dating to indicate the phylogenetic position of total-group animals, where association between modern bacterial symbionts and multicellular animals probably had not been developed. Ecdysozoa is a group of protostome animals including nematodes, arthropods, and several other phyla (Telford et al. 2008). The two arthropods were *Drosophila melanogaster* (fruit fly) and *Daphnia pulex*, belonging to Hexapoda (insects and related groups) and Crustacea respectively. *Daphnia pule*, the common species of water flea, is a model species, and was the first crustacean whose genome has been sequenced. Crustacean is the closest relative of insects known to date (Thomas et al. 2020). Hence, the LCA of *Drosophila melanogaster* and *Daphnia pulex* well defines the total group of insects, which was further used in RTC-based dating. The addition of the model organism and free-living transparent nematode *Caenorhabditis elegans*, allowed using an additional fossil (see Note S3.2.1) to better calibrate the evolution of arthropods and other animals.

##### 1.1.2 Bacteria in the main dating analysis (148-genome dataset)

To select bacterial genomes used in molecular clock analysis, we retrieved all 265 genomes used as the primary dataset in a recent phylogenomics study that investigated the root of the bacterial tree of life (Coleman et al. 2021). To reduce the computational burden of molecular clock analysis, we applied TreeCluster v1.0.3 (Mai et al. 2017) with the phylogenetic depth cut-off of 1.8 on a phylogenomic tree of all of these 265 genomes built by the rapid tree construction software FastTree (Price et al. 2010). This generated 81 representative genomes covering all phylum-level lineages used in the original 265-genome dataset. We further added the following two classical model organisms *Escherichia coli* from γ-proteobacteria and *Bacillus subtilis* from Firmicutes, and two from β-proteobacteria, *Rhodocyclales bacterium* GWA2 65 20 and *Neisseria gonorrhoeae,* as no β-proteobacteria was sampled in Coleman et al. 2021.

The following six bacterial organisms were added to allow using relevant calibrations within bacteria. Two of them were *Thiocapsa marina* 5811 and *Nitrosococcus oceani* ATCC

19707 in Chromatiaceae (γ-proteobacteria). They are associated with the biomarker calibration based on okenone (Brocks et al. 2005). The other three were all oxygenic cyanobacteria: *Nostoc* sp. PCC 7107, *Xenococcus* sp. PCC 7305, *Oscillatoria* sp. PCC 10802, and *Gloeobacter violaceus* PCC 7421, according to the section of "Cyanobacteria fossil based dating strategy" in our previous study (Wang and Luo 2021). These three organisms are associated with three calibrations of cyanobacteria. See the Note S3.2.2 for more information.

As to the bacterial symbionts used in RTC-based dating, we selected 28 genomes to represent 19 RTCs from the nine ancient clades of bacterial symbionts (see Note S3.1.1 for more details): four in Tenderises, six in Spirochaetota, three in Elusimicrobiota, three in Chlamydiae, three in Rickettsiales, two from Holosporales, three in Legionellales, two in the aphid endosymbiont *Buchnera*, and two in the cockroach endosymbiont *Blattabacterium*. Note that two of them (*Parachlamydia acanthamoebae* and *Elusimicrobia bacterium* CG1 02 37 114) are included in the primary dataset of Coleman et al. 2021.

Collectively, 148 organisms were used in the main molecular clock analysis: 91 non-symbiotic bacteria, 28 symbiotic bacteria, and 29 eukaryotes attached as the mitochondrial subtree.

##### 1.1.3 Genomes used in molecular dating of the high-resolution timetree of bacteria

The genomes were obtained from the Genome Taxonomy Database (GTDB) v95 (Parks et al. 2022). According to the taxonomic annotation in GTDB, a genome was randomly selected from each genus, and their genome sequences were downloaded from the NCBI Genbank or RefSeq databases, depending on the assembly accession recorded in GTDB (GCA for Genbank and GCF for RefSeq). Five genomes (*Nitrosococcus oceani* ATCC 19707, *Thiocapsa marina* 5811, *Nostoc* sp. PCC 7107, *Oscillatoria* sp. PCC 10802, and *Xenococcus* sp. PCC 7305) were removed because their assembly accessions recorded in GTDB were no longer available in NCBI. This totalled XXX genomes (see Data S4 for the full list).

#### 1.2 Molecular dating

##### 1.2.1 Settings of the MCMCTree analysis

The time unit in MCMCTree analysis was set to 100 Ma. The prior on divergence times for nodes without a calibration was constructed using a birth-death process (Nee et al. 1994). The parameters of the birth-death process were set as: birth rate = 1 (birth of one lineage per time unit), death rate = 1 (death of one lineage per time unit), and the taxon sampling proportion =

0, which specifies a uniform kernel and thus represents a diffuse time prior as used in the studies (dos Reis et al. 2012; Dos Reis et al. 2015). The prior on the variance of branch-wise log-transformed rate was specified as “1 10 1” which follows a gamma distribution Gamma(1,10). This setting should account for much among-lineage rate variation (Brown and Yang 2011). The Dirichlet-gamma prior on the substitution rate for each gene partition was set as “rgene_gamma = 1 50 1”, which specifies a gamma distribution Gamma(1,50) and indicates a mean substitution rate of 0.02 amino acid substitution per 100 Ma, as calculated from previous studies (Wang and Luo 2021; Álvarez-Carretero et al. 2022). The impacts of alternative settings of the above parameters on posterior dates were also investigated (Figs. S11-S12; Data S3).

For MCMCTree analysis of the eukaryote tree, each MCMC chain was run sampling every 100 iterations to provide $10^5$ posterior samples (after burn-in) with ten partitions using the independent rates (IR) model, the best-fit clock model determined by mcmc3r (Dos Reis et al. 2018) (Note S1.2.3). For MCMCTree analysis of the bacterial tree, each MCMC chain was run sampling every 100 iterations to generate $3 \times 10^5$ posterior samples (after burn-in), except for the analysis under the focal scheme where $6 \times 10^5$ posterior samples were collected. The autocorrelated rate (AR) model, as determined by mcmc3r as the best fit, was used (Note S1.2.3). Due to the large proportion (usually more than 90%) of posterior samples rejected in the rejection sampling step (Step 3) with the RTC approach, a single partition was used in the main analysis to reduce the computational time. Alternative analyses were performed with different clock model or different number of partitions (Figs 3, S11; Data S3). For analysis with more than one partition, genes were clustered into a user-specified number of partitions by fitting the substitution rates, which were estimated by Codeml from the PAML package, into a Gaussian mixture model using ClusterR (Mouselimis 2022). Posterior dates were compared with effective time priors (“usedata = 0”) to ensure that their distributions were different and therefore sequences were informative in MCMCTree analysis (Table S4).

1.2.2 Bayesian sequential molecular dating analysis using MCMCTree

Suppose the phylogenomic dataset $D$ (i.e., sequence alignment) is composed of two non-overlapping subsets $D_1$ and $D_2$ conditionally independent of each other, then we can write the posterior of parameters $\theta$ (including the ages and substitution rates among others) as

$$f(\theta|D) \propto f(\theta)f(D_1, D_2|\theta)$$

$$= f(\theta)f(D_1|\theta)f(D_2|\theta)$$

$$\propto f(\theta|D_1)f(D_2|\theta).$$

It is obvious from the above formula that the posteriors (a posterior distribution) of parameters calculated with the first subset of data $f(\theta|D_1)$ can be used as the prior for following analysis on $D_2$ (dos Reis et al. 2012).

Five distributions are included in MCMCTree for setting calibrations, namely uniform distribution, truncated Cauchy distribution, gamma distribution, skew-normal distribution, and skew-t distribution. For each node, the posterior date was fitted by skew-normal distribution and skew-t distribution using the R package "sn" (Azzalini 2019), and gamma distribution was fitted using the R package "fitplusdistr" (Delignette-Muller and Dutang 2015). The other two were not used because they are either too simplistic (uniform distribution) for approximating posterior dates or difficult to fit as a truncated distribution (truncated Cauchy distribution). The log-likelihood value of the fit was calculated using the "sn::st.mple" R function for skew-normal and skew-*t* distribution, and using the R function "fitdist" for gamma distribution. The degrees of freedom of skew-normal, skew-*t*, and gamma distribution are three, four, and two, respectively. AIC was calculated as $AIC = 2k - \log(\hat{L})$ where $k$ is the degree of freedom and $\hat{L}$ is the maximum of the likelihood for the model (Akaike 1974). The parameters of each fitted distribution are given in Fig. S4.

One has to keep in mind that the sequential Bayesian molecular clock procedure used here is an approximate approach. When fitting the posteriors obtained from the first-step sequential analysis to the marginal posteriors and when using them as priors in the second-step analysis, we considered the posterior dates of different nodes are independent. However, this might not hold. One such source comes from the requirement that any child node must be younger than its parent node. In brief, user-specified calibrations are assigned to nodes on the tree and may not satisfy the above requirement. Therefore, the software must "truncate" the calibration densities to satisfy the constraint. As such, the resulting priors actually applied by the dating software are known as the effective prior (Barba-Montoya et al. 2017). As detailed in Álvarez-Carretero et al. 2022, in case of a significant overlap in the dates between a child node and its parent node in the first-step sequential analysis, there would be a poor approximation by fitting the marginal distribution with this method due to the truncation effect.

For the first-step sequential dating analysis, we estimated an eukaryote timetree using 320 orthologs conserved in eukaryotes identified in Strassert et al. 2021. As shown in Fig. S2, three nodes (nodes 31, 45 and 46) in the eukaryote tree displayed large overlap in the posterior dates with its parent or child node. Hence, in the second-step sequential analysis (i.e., dating the bacterial tree of life with the 148-genome set), we calibrated all but the above three internal nodes with the best-fit probability distribution described above. To check the quality of approximation, we conducted the following comparisons (Fig. S4). i) the posteriors of the dates of eukaryote nodes obtained in the first-step sequential analysis vs. the probability distribution fitted to it. If the two were nearly indistinguishable, it suggests that the probability distribution well fit the posterior dates obtained in the first-step sequential analysis. ii) the best-fit parametric probability distribution vs. the effective priors in the second analysis by setting *usedata* = 0 in MCMCTree. If the two were highly similar, then the truncation effect mentioned above had little impact, and accordingly the sequential method would work for such nodes.

Note that sequential Bayesian analysis is distinct from the use of secondary calibrations (using time estimates from other studies as calibrations), which was often used in previous studies to date bacterial evolution, for the following reasons (Dos Reis et al. 2018; Álvarez-Carretero et al. 2022). First, secondary calibrations were treated as point calibrations (with the uncertainties on node age estimates ignored). Second, in many cases the data analysed under the secondary calibration was the same as the data analysed to obtain the calibration in the first place, leading to duplicate use of the same data, thus a squaring of likelihood. On the other side, the Bayesian sequential approach is justified if the datasets are independent and do not overlap.

1.2.3 Selection of the best-fit clock model

We employed the mcmc3r package (Dos Reis 2017) to determine the best-fit clock models. It allows using a stepping-stone method (Xie et al. 2011) to calculate the marginal likelihood of three competing clock models available in MCMCTree: the independent rate (IR) model, the autocorrelated model (AR; more specifically, the geometric Brownian motion rate model), and the strict clock model (STR). The best-fit clock model for each gene was determined according to the marginal likelihood. Note that mcmc3r works with only the exact likelihood but not the approximate method in MCMCTree, and that the exact likelihood method works for only nucleotides. Hence, we recoded the amino acids into 4-character states based on the Dayhoff-4 scheme.

In the first-step sequential analysis, i.e., dating the eukaryote tree based on the 320 orthologs identified in (Strassert et al. 2021), the IR and AR models were favoured by similar proportions of genes (53% vs. 47%; *eukaryotic_tree* in Fig. S13). Because date estimates obtained by IR and AR were similar, we arbitrarily chose the IR model. For the second-step analysis, i.e., dating the bacterial tree (148 species), we followed prior studies (McGowen et al. 2020; Wang and Luo 2021) to overcome the huge computational burden: i) We compiled two reduced datasets by randomly selecting 20 and 40 species, ii) We fixed the root prior at 4.0 Ga and applied only a single internal calibration that constrained the age of the LCA of eukaryotes to 1.7-1.5 Ga (here we did not want to estimate divergence times but simply select a better-fit model). The AR model was shown to have the higher marginal likelihood for 69% and 73% of the genes for the 20- and 40-species sets (Fig. S13), respectively, and was therefore selected as the preferred model, as done in a previous study (Álvarez-Carretero et al. 2022).

The burn-in, sampling frequency, and number of samplings were set as 5000, 2, and 1000, respectively, for all mcmc3r analyses.

1.2.4 Selection of genes with high verticality or low among-branch rate heterogeneity

To test if genes used in analysis impacted the posterior dates in molecular dating of the bacterial tree, the 19 mitochondria-originated genes were evaluated with ΔLL as a metric of marker gene verticality (i.e., if genes were vertically inherited in evolution) (Moody et al. 2022), and relative rate difference which measures the extent of the violation of molecular clock for a gene (Wang and Luo 2021). Specifically, ΔLL calculates the difference in likelihood in log scale between a constrained maximum likelihood tree based on the topology of the species tree and the maximum-likelihood gene tree. Genes with a higher ΔLL indicate large topology difference between the gene tree and the species tree, thus lower verticality (Moody et al. 2022). To this end, ΔLL well serves as a proxy for the extent to which the gene tree differs from the species tree. The relative difference in rate between mitochondria and others was calculated as $\frac{|r_{mito}-r_{bac}|}{\max(r_{mito}, r_{bac})}$ (Wang and Luo 2021), where the substitution rates of mitochondria or bacteria lineages were estimated by Codeml from PAML (Yang 2007). The higher the relative rate difference is, the more rate heterogeneity between lineages would be expected. Selecting only those with the smallest relative rate difference in molecular dating could ensure less violation of molecular clock caused by the fast-evolving mitochondrial lineages, as used in Wang and Luo 2021. We selected the 5, 10, and 15 top-ranking genes using each of the above two measures and repeated the molecular clock analyses.

Additionally, we performed the analysis with the full set of 32 mitochondria-originated genes.

<u>1.2.5 MCMC convergence diagnostics</u>

The convergence of MCMC analysis was checked by running the MCMCTree analysis for twice and comparing the date estimates. Effective sample size (ESS) was calculated using the R package coda (Plummer et al. 2006) to ensure all posteriors have ESS of at least 200.

### 1.3 Linear regression analysis of the relationship between the divergence time and 16S rRNA gene divergence

We performed regression analysis to study the relationship between 16S rRNA gene divergence and the divergence time at a family level. For each family with at least five members (remember that one genome was randomly selected from each genus), the 16S rRNA gene divergence was defined as the maximum pairwise substitution rate (number of nucleotide substitutions per site) within the family. The analysis was based on the taxonomically high-resolution timetree of bacteria.

The simple linear regression takes the form

$$Y = \beta_0 + \beta_1 X + \epsilon,$$

where $Y$ is the dependent variable (16S rRNA gene divergence), $X$ is the independent variable (divergence time), $\beta_0$ is the intercept, $\beta_1$ is the slope, and $\epsilon$ is the error item.

Note however that families from the same phylum might have a more similar 16S rRNA gene substitution rate. To model this effect, we applied the following three linear mixed models to consider the "phylum" as a random effect using R packages "lmer" (Bates et al. 2021) and "lmerTest" (Kuznetsova et al. 2017): the random-intercept model, the random-slope model, and the random-intercept random-slope model.

For the random-intercept mixed modelling framework, $\beta_0$ and $\beta_1$ are fixed effects (or the population-averaged values), and $b_i$ is thought as a random effect for family $i$, depending on which phylum the family belongs to. It takes the form

$$Y = (\beta_0 + a_i) + \beta_1 X + \epsilon.$$

The random effect can be regarded as each subject's deviation from the fixed intercept. In this model, the predictions vary depending on the random intercept term of each subject, but the slopes are always the same. In other words, different phyla could have different intercepts, denoted by the term $\beta_0 + a_i$.

The second linear mixed model specifies a random effect term on the slope. Hence, predictions may vary according to each subject's slope while having the same intercept. In other words, different phyla could have different intercepts, denoted by $\beta_1 + b_i$. The random-slope model takes the form

$$Y = \beta_0 + (\beta_1 + b_i)X + \epsilon.$$

The third linear mixed model specifies a random effect term on both the slope and intercept. Therefore, predictions may vary according to each subject's slope and the intercept. In other words, different phyla could have different slopes and intercepts, denoted respectively by $\beta_1 + b_i$ and $\beta_0 + a_i$. The random-slope model takes the form

$$Y = (\beta_0 + a_i) + (\beta_1 + b_i)X + \epsilon.$$

Note that in the main linear regression analysis we followed prior studies (Moran et al. 1993; Ochman et al. 1999; Kuo and Ochman 2009) to set the intercept to zero, because in theory divergence time of zero corresponds to 16S gene divergence of 0%. Alternative analyses showed that including intercept as a parameter did not change the general pattern (Data S5).

#### 1.4 Data visualization

Multiple sequence alignments were visualized with BioEdit v7.0.5.3 (Hall 1999). Phylogenetic trees were visualized using FigTree v1.4.3 (Rambaut 2010), iTOL v4 (Letunic and Bork 2019), TreeGraph v2.5 (Stöver and Müller 2010), the Newick Utilities (Junier and Zdobnov 2010), and MCMCtreeR (Puttick 2019).

**Supplementary Note 2. Determining the root position of the bacterial tree by the time non-reversible model (NONREV)**

2.1 Phylogenetic reconstruction of the bacterial tree under non-reversible model

To root the bacterial tree, we applied the time non-reversible substitution model recently implemented in IQ-Tree (Minh et al. 2020). This method infers the root as part of maximum-likelihood phylogenetic reconstruction where different root positions may have different likelihoods, allowing outgroup-free inference of the root. Thus, it potentially avoids long branch attraction caused by the use of fast-evolving or very distantly related sequences, which is a common issue in rooting deep phylogenies (Coleman et al. 2021). We first applied each of the three empirical site-heterogeneous mixture models C20, C40, and C60 (settings: -m LG+G+Cxx) to build unrooted phylogenies using the 60 marker genes identified in Coleman et al. 2021. For each of the three trees built with time-reversible model LG+G+Cxx, we followed the study (Naser-Khdour et al. 2022) to build four rooted phylogenies using the non-reversible model (settings: --model-joint NONREV) based on: i) a fixed tree topology according to those built by the reversible model without partitions, ii) a fixed tree topology according to those built by the reversible model with partitions, iii) an initial tree built by the time-reversible model without partitions, iv) an initial tree built by the time-reversible model with partitions. Partitions were identified by ModelFinder using the time-reversible model (settings: -m MFP+MERGE -mset LG,JTT,WAG -mrate E,I,G,I+G).

Note the following points in tree reconstruction using the non-reversible model. First, an initial tree, also known as start tree, is the tree that serves as a starting point from which heuristics tree searching is performed with modern maximum-likelihood tree reconstruction algorithms. The topology of the initial tree is likely to change during maximum-likelihood tree reconstruction. A good initial tree is important for accurately inferring the phylogeny (Yue et al. 2009; Money and Whelan 2012). Second, a fixed topology means that, during tree reconstruction (using the non-reversible model), the tree topology built by a reversible model is not changed, but a root position is identified, and branch lengths are estimated to maximize the likelihood. The use of the above combinations thus allows capturing different possibilities to better search for a tree with the highest likelihood based on the non-reversible model.

In brief, for the four strategies mentioned above, strategies i) and ii) find a most likely root using the NONREV model on the tree topology inferred by reversible model, while strategies iii) and iv) employ the NONREV model to infer both the root and tree topology with the initial tree inferred by reversible model serving as a “starting point”.

2.2 Determining the root position of the bacterial tree

As shown in Fig. S1B-S1E, in most analyses, the clade consisting of Fusobacteria and DST (deep grey in Fig. S1) were the most basal clade or were placed at a position close enough to the root. On the other side, the root position varied between analyses (Fig. S1B-S1E). Further, as evident by the rootstrap statistic (Naser-Khdour et al. 2022), there was not a single root placement that received a significantly high support to statistically distinguish it from others () in all analyses (Fig. S1D-S1E). The rootstrap support provides information on the support for a given root placement. Basically, a bootstrap analysis is conducted to obtain rooted bootstrap trees using NONREV model. The rootstrap support for each branch is defined in the ML tree as the fraction of rooted bootstrap trees with the root position on that specific branch. Although there is no consensus as to how large the recently developed rootstrap is in order that the root is considered “high confidence”, obviously rootstrap values varied across nodes when different models were used (Fig. S1D-S1E).

When compared with the reversible model that did not involve Cxx (*REV_par*), NONREV displayed a lower AIC value in most analyses indicating that it better fit the data than its time reversible counterpart (Table S2A). Nevertheless, all NONREV displayed poorer model fit than LG+G+Cxx, indicated by their higher AIC values. In fact, the time reversible mixture model LG+G+C60 (*REV_C60* in Table S2A) was selected as the best-fit model. It is important to emphasize that NONREV cannot be used with the Cxx mixture model. Hence, the reason for its lower model fit than LG+G+C60 (*REV_C60*) is likely because the benefit of using the non-reversible model did not counteract the benefits of accounting for the compositional heterogeneity by the mixture model Cxx.

The above indicates that while it is difficult to figure out which root position is the “best”, it is very likely that Fusobacteria/DST represents one of, if not the, earliest-split clade among all analysed in the present study. This is also indicated by the high rootstrap support at positions close to Fusobacteria/DST (Fig. S1D-S1E). In the main analysis, we used the tree topology inferred by LG+G+C60 as it showed the highest model fit (*REV_C60* in Table S2A). We arbitrarily placed the root at the clade consisting of Fusobacteria and DST (root0 in Fig. S1) in the main analysis, and included another two potential root positions close to root0 but on different branches (root1 and root2 in Fig. S1A) as additional analysis (Fig. 3D). These two alternatives are also suggested as likely roots in the study (Coleman et al. 2021). Further, some earlier studies argue for a root of the bacterial tree on the CPR branch (Hug et al. 2016; Méheust et al. 2019; Zhu et al. 2019) using archaea as the outgroup. This hypothesis was not supported by our and other analyses that employed newly developed outgroup-free methods for rooting (Coleman et al. 2021) or that used carefully selected orthologs and substitution models with better fit to data (Moody et al. 2022). However, to test its impact on the time estimates, we also forced the tree to be rooted at the CPR clade (rootCPR) as an alternative dating analysis.

**Supplementary Note 3: Time constraints.**

### 3.1 Symbiosis-informed RTCs

3.1.1 General principle of selecting bacterial lineages to represent RTCs based on ASR

In the present study, we selected four in Tenericutes, six in Spirochaetota, three in Elusimicrobiota, three in Chlamydiae, three in Rickettsiales, two from Holosporales, three in Legionellales, two in the aphid endosymbiont *Buchnera*, and two in the cockroach endosymbiont *Blattabacterium*, as internal nodes where symbiosis-based RTCs are imposed. In general, these lineages were selected to maximize the effect of each RTC based on the following four criteria. A graphical illustration is provided in Fig. S6A.

First, if the ancestral state of a child node is inferred to be (almost) the same as its parent node, then using the parent node as the RTC makes more sense. This is because that the child (e.g., node 1) is younger than its most likely host (animal) is already implied by the same constraint imposed on its parent (node 2). Therefore, constraining the age of the parent instead of its child to be younger than that of the host apparently better constrains age of the whole symbiont clade.

Second, in case that the child and the parent are inferred to have different states, the time constraints on both nodes may be effective. As shown in Fig. S6A, that node 2 is younger than its parent node 3 does not necessarily mean that node 3 has to satisfy the same constraint because their preferred hosts are different. Hence, both internal nodes may be included.

Third, because the molecular dating strategy involves *post hoc* subsampling a large number of the posterior distribution of timetrees based on RTCs, to save time, shallow lineages (late-split) are generally not considered. Also, it is apparent that constrains on late-split nodes contribute less to deep time dating than those on early-split lineages.

Fourth, while the ASR was conducted with the 16S rRNA gene, the dating analysis was performed using mitochondria-originated genes. Thus, we had to ensure that all selected lineages in ASR analysis have genome sequenced (indicated by the green circle in the layer next to the taxon name in Fig. S4). As such, Sym2 and Sym5 in Fig. S4 must be excluded because their genome sequences are not available.

Collectively, for the example in Fig. S4, it is Sym1, Sym3, and Sym4 that would be selected to represent two RTC nodes, i.e., node 2 and node 3, in subsequent analysis.

3.1.2 Classification of the lifestyle and ASR for clades mainly comprised by symbionts

**Group:** Rickettsiales

**Lifestyle classification:** We classified samples collected from an animal host as animal-associated, and those collected from protists as protist-associated. Samples where no host is indicated in the metadata were given equal probability to free-living and protist-associated lifestyle. The only exception is those recently reported in the study (Schön et al. 2022), which represent the earliest-split Rickettsiales family Mitibacteraceae, and members from this lineage was classified as free-living (see reasons below).

**Justification:** Rickettsiales is an order of α-proteobacteria traditionally recognized as mainly animal symbionts. Many members from the genera *Rickettsia*, *Anaplasma*, *Ehrlichia*, and *Neorickettsia* are medically relevant bacteria, and *Wolbachia* infects almost all filarial nematodes and around half of insect species (reviewed in Bourtzis and Miller 2003; Werren et al. 2008). However, increasing evidence supports a much broader range of hosts of Rickettsiales expanding from marine invertebrates (*Ca.* Aquarickettsia) (Klinges et al. 2019) to diverse protists especially *Paramecium* and amoebae (Castelli et al. 2016). Our compiled datasets included 3159 16S rRNA sequences, with 2848 (90%) detected from animals and 234 (7.4%) detected from protists, respectively (see Data availability). Most protist-associated members were from the two families *Ca.* Midichloriaceae and *Ca.* Rickettsiaceae (Fig. S5), in agreement with prior studies (Castelli et al. 2019; Schön et al. 2022).

It is worth noting that 77 (2.4%) collected sequences do not have an identified host. While the possibility that they could represent free-living members cannot be ruled out, considering the large number of host-associated Rickettsiales and that 16S rRNA gene-based molecular method may not distinguish free-living bacteria from protist-associated ones without additional information, it seems at least equally if not more possible that they are associated with a protist host. Hence, samples where no host is indicated in the metadata were given equal probability to free-living and protist-associated lifestyle, which reflects a conservative interpretation of the lack of any known eukaryotic host. The only exception is those recently reported in the study (Schön et al. 2022), which represent the family Mitibacteraceae, the earliest-split clade within the Rickettsiales (another early-split family Athabascaceae reported in the same study was not included as no 16S rRNA sequence was available for it). They were assigned as free-living because comparative genomics analysis indicated the lack of several important genes for establishing symbiosis for classical Rickettsiales, such as ATP/ADP translocase (Schön et al. 2022).

**Alternatives:** We alternatively classified those without any host indicated in the metadata as free-living, instead of assigning an equal probability to both free-living and protist-associated lifestyle.

**Group:** Holosporales

**Lifestyle classification:** We classified Holosporales detected in ciliates as ciliate-associated, and those from any other eukaryote sources as protist-associated.

**Justification:** Holosporales was previously classified as a subclade of the α-proteobacterial order Rickettsiales. However, this is considered as a phylogenetic artefact due to the poor-fit model in phylogenetic reconstruction and recent studies have suggested Holosporales as independent order- or family-level group closely related to Rhodospirillales (Muñoz-Gómez et al. 2019; Fan et al. 2020). All known Holosporales members are identified from protist host, which includes ciliates, amoebae, cercozoans and euglenozoans among others. Ciliate-dwelling Holosporales are mostly distributed in the genera *Holospora* and *Gortzia* (Schrallhammer and Potekhin 2020). In contrast, those associated with other protists are distributed in other Holosporales lineages (Hess et al. 2016; Chan et al. 2018). In our collected 16S gene dataset of Holosporales comprised by 126 sequences, 73 and 31 were considered as ciliate- and protist-associated, and 22 did not indicate any host in the metadata, which however do not indicate that they are necessarily free-living (see Data availability). Like the case of Rickettsiales, it seems equally possible that they are associated with a protist host.

**Alternatives:** Like the case of Rickettsiales, we alternatively classified those derived from environmental samples as free-living, instead of assigning an equal probability to both free-living and protist-associated lifestyle.

**Group:** Legionellales

**Lifestyle classification:** We classified those collected from animal and protist hosts as animal- and protist-associated respectively. For those where no host is indicated in the metadata, an equal probability was given to the free-living and protist-associated lifestyle.

**Justification:** Legionellales is a γ-proteobacterial order all of whose known members adapt an intracellular lifestyle during life cycle and includes several accidental human pathogens such as *Legionella pneumophila* (Legionnaires' disease) and *Coxiella burnetiid* (Q fever) (Duron et al. 2018). There are two described families of this order, Coxiellaceae and Legionellaceae (Fig. S5). Most members of Coxiellaceae, e.g., *Coxiella*, *Rickettsiella*, and *Diplorickettsia* are found to infect various animal hosts ranging from various invertebrates to mammals. The exceptions are *Aquicella* and the newly identified *Berkienalla* which are found to be associated with protists and which form independent phylogenetic clades

(Hugoson et al. 2022). Notably, although Legionellaceae can grow in natural biofilms, they need protist host like amoebae or ciliates to replicate intracellularly. *Legionella* can co-exist with their amoeba host in a benign relationship upon infection where the amoeba may not use the bacterium as food and the bacterium does not cause any disease to the amoeba host. It is thought that protists serve as natural reservoirs of *Legionella* and promote disease in human (Conza et al. 2013; Boamah et al. 2017). For this reason, a conservative way may be to give those without any host indicated in the metadata [313 out of 817 (38%); Data availability] an equal probability of free-living and protist-associated.

**Alternatives:** Like the case of Rickettsiales, we alternatively classified sequences derived from environmental samples as free-living, instead of assigning an equal probability to both free-living and protist-associated lifestyle.

**Group:** Chlamydiae

**Lifestyle classification:** We classified those collected from animal and protist hosts as animal- and protist-associated respectively. For those where no host is indicated, an equal probability was given for the free-living and protist-associated lifestyle.

**Justification:** Chlamydiae is a bacterial phylum usually placed as a sister group to Verrucomicrobia, which together form the so-called PVC clade with Planctomycetes (Gupta et al. 2012). Traditionally, members of the phylum Chlamydiae include pathogens of animals and symbionts of ubiquitous protists. In general, animal pathogenic members are divided into two groups, one mainly consisting of the genus *Chlamydia* some of which are famous pathogens that can cause chlamydiosis in vertebrates (Vanrompay et al. 1995; Bachmann et al. 2014), and the other including *Actinochlamydia*, *Similichlamydia* and *Piscichlamydia* which may cause epitheliocystis in fish, a condition associated with epithelial hyperplasia, hypertrophy and inflammation of the infected tissue (Stride et al. 2013; Sood et al. 2018). Animal-associated Chlamydiae infect their host through a strictly intracellular developmental cycle. Other Chlamydiae can thrive within unicellular eukaryotes particularly amoebae (Lienard et al. 2017; Haselkorn et al. 2021) but there is evidence for alternative protist host like Heterolobosea (Casson et al. 2008). These protist hosts are highly diverse and abundant in aquatic and terrestrial ecosystems. Some Chlamydiae lineages, for instance the family Rhabdochlamydiae (Halter et al. 2022), have both protist- and arthropod-associated members. Recently, members of Chlamydiae were detected in ocean-floor environments using metagenomics, some of which are likely novel lineages within the phylum (Dharamshi et al. 2020). It is unknown if these marine sediment Chlamydiae have any eukaryotic host

(Dharamshi et al. 2020), but to date all cultured environmental Chlamydiae are strictly intracellular (Collingro et al. 2020). For this reason, a conservative way may be to give those without any host indicated in the metadata [175 out of 656 (27%); Data availability] an equal probability of free-living and protist-associated (Collingro et al. 2020).

**Alternatives:** We alternatively classified those without identified host as free-living instead of giving free-living and protist-associated an equal probability.

**Note:** We selected three genomes from this group to represent two RTCs (*Candidatus* Rhabdochlamydia sp. W815, *Candidatus* Clavichlamydia salmonicola, and *Parachlamydia acanthamoebae*). As displayed in Fig. S5, the ancestral lifestyles are very similar, suggesting the younger node could be redundant. However, as Chlamydiae is an important phylum-level taxonomy group, we still considered all of the three in the focal analysis.

**Group:** Tenericutes

**Lifestyle classification:** We classified those found in animals as animal-associated, and those found in fungi as fungus-associated. Those found in other habitats were classified as free-living. For mollicutes found in plants, because it is widely known that they are insect-transmitted plant endosymbionts and require both insects and seed plants to replicate, we required that their origin time to be no older than that of both insects and seed plants.

**Justification:** Formerly proposed as a phylum but more recently suggested to be affiliated with the Bacilli clade of Firmicutes (Davis et al. 2013; Wang et al. 2020), Tenericutes consists of bacteria that do not have a peptidoglycan cell wall. The most well-known clade of Tenericutes is Mollicutes, which contains many medically or agriculturally relevant genera. Almost all reported mollicutes adapt to an endosymbiotic lifestyle. Perhaps the most abundant ones in nature are those that are animal endosymbionts which have been found in most major animal lineages like human (Waites and Talkington 2004), mammal (Deeney et al. 2021), reptile (Brown et al. 1999), arthropod (Sapountzis et al. 2018), mollusc (Pimentel et al. 2021) and jellyfish (Cleary et al. 2016). Animal endosymbiotic mollicutes are most often found in the genera *Mycoplasma*, *Ureaplasma*, and *Acholeplasma*. While some of them are famous human pathogens (e.g., *Mycoplasma. pneumoniae*, *Mycoplasma hominis*, *Ureaplasma urealyticum*), others may behave as “silent parasites” that do not apparently harm their hosts (Borchsenius et al. 2020).

All known members from the genus *Phytoplasma* and some from *Spiroplasma* (Fig. S5) are causative agents of plant diseases and are transmitted by phloem-feeding insects from only the order Hemiptera which includes psyllids, leafhoppers and planthoppers. While they can be pathogenic to some insect hosts, in general plant-associated mollicutes do not negatively affect their insect vector's fitness (Hogenhout et al. 2008). These plant-symbiotic mollicutes cycle between plant and insect-vector hosts where they survive and replicate in both hosts although some can be passed on to next generations of insect vectors or plant seeds (Ammar and Hogenhout 2006), indicating the essentiality of both hosts to plant symbiotic mollicutes.

The genus *Moeniiplasma*, which was discovered in recent years (Naito et al. 2017), consists of a unique group of mollicutes in that all its members are endosymbionts of arbuscular mycorrhizal fungi (AMF, subphylum Glomeromycotina) and are found in most AMF species. The effect of having *Moeniiplasma* on the host fungus, however, is unknown.

Different from Mollicutes, other Tenericutes can be found in a more diverse range of environments. This is evident by the recent discoveries of free-living *Izemoplasma* (Skennerton et al. 2016; Zheng et al. 2021) and *Haloplasma* (Antunes et al. 2008) in deep-sea cold seep and brine pool, respectively. These marine environmental Tenericutes display adaptive flexibility and metabolic versatility, and likely represent early-split lineages of Tenericutes (Wang et al. 2020).

**Group:** Spirochaetota

**Lifestyle classification:** we classified those detected in animals as animal-associated, and those detected in environmental sources as free-living.

**Justification:** The phylum Spirochaetota contains bacteria can be distinguished from other flagellated bacteria by their long, thin and spiral (or wavy) cell bodies (Nakamura 2020). The phylum Spirochaetota consists of four major families Spirochaetaceae, Leptospiraceae, Brachyspiraceae, and Brevinemataceae (Gupta et al. 2013). A unique feature of Spirochaetota is their motility, mediated by periplasmic flagella with a rapid drifting rotation. The unique morphology and rotational motility as well as directional control of flagellar motors enable Spirochaetota to be successful in diverse ecological niches as commensals or parasites of animals, as metabolic symbionts of insects, and as free-living bacteria. These skills are considered important for some members of this phylum to invade and colonize host tissues, eventually leading to diseases like leptospirosis (*Leptospira interrogans*), Lyme disease (*Borrelia burgdorferi*), syphilis (*Treponema pallidum*), swine dysentery (*Brachyspira hyodysenteriae*) and many other animal diseases (Karami et al. 2014).

Members of the family Spirochaetaceae adapt to different lifestyles (Haake 2009). At one extreme is the obligate pathogen, *Treponema pallidum*, the most invasive of pathogenic treponemes (Norris et al. 2001). At the other extreme, many members of this family, particularly the genus *Spirochaeta*, are free-living bacteria (Leschine et al. 2006). In between these two extremes are commensal or parasitic organisms with life cycles involving insects, animals, or both, as in the case of *Borrelia* (Haake 2009). The family Brachyspiraceae contains bacteria that colonize digestive tracts of animals either as commensals or parasites (Rosenberg 2014). The family Leptospiraceae includes both environmental saprophytes and animal parasites that cycle between aquatic environments and animal host via renal tubules (Picardeau 2014).

**Group:** Elusimicrobiota
**Lifestyle classification:** We classified those found in the gut of termites or ants as protist-associated because these *Elusimicrobia* are actually symbionts of single-cell eukaryotes living in the gut. Those from environmental sources were classified as free-living.
**Justification:** Elusimicrobiota is an phylum-level taxonomic group placed as a basal lineage to most Gracilicutes (Coleman et al. 2021), It was previously known as Termite Group 1 (TG1), because its members were found to be endosymbionts (Ohkuma et al. 2007) or ectosymbionts (Mikaelyan et al. 2017) of various flagellated protists (e.g., Parabasalia and Preaxostyla) that live in the gut of termites and ants. Recent metagenomic studies have indicated that free-living members belonging to this phylum are prevalent in diverse ecosystems like sewage sludge, marine environment, contaminated sites and soils (Herlemann et al. 2007; Méheust et al. 2020). Compared with gut microbiome *Elusimicrobia*, those found in environmental samples display higher metabolic diversity and it was predicted that animal-associated members evolved from free-living species (Méheust et al. 2020).

**Group:** *Buchnera*
**Lifestyle justification:** We classified all *Buchnera* members as insect-associated.
**Justification:** *Buchnera*, a genus of γ-Proteobacteria, is found exclusively in the specialized cells of aphids (Hemiptera: Sternorrhyncha: Aphidoidea), and constitutes an essential component of their life cycle. *Buchnera* has been considered a prime example of how intimate symbiotic relationships between organisms can have important effects on their evolution and ecological success. All aphids carry *Buchnera* in their cell cytoplasm. Roughly 5.6 million cells of *Buchnera* were estimated to reside in a 10-day-old aphid (Douglas 1998). Aphids either die or cannot reproduce when treated with antibiotics (Houk and Griffiths 1980). In exchange for a stable, nutrient-rich environment, *Buchnera* supply their host with

the amino acid tryptophan, which is occasionally found in plant sap, and they also help with the production of vitamins and leucine (Klepzig et al. 2009). No free-living members belonging to *Buchnera* has been reported to our knowledge.

In general, *Buchnera* has been thought to co-diverge with their aphid host, as supported by both experimental evidence for their maternal transmission (Koga et al. 2012) and phylogenetic evidence (Munson et al. 1992; Moran 2001; Liu et al. 2013). Studies have based on this information to use secondary calibration to calibrate the divergence time of *Buchnera* by the estimated divergence time of aphids' crown group (Moran et al. 1993; Kuo and Ochman 2009). Nevertheless, to be conservative, we assume the LCA of *Buchnera* as insect-associated, instead of aphid-associated in our RTC-based molecular dating. A major difference from prior studies is that we assumed that the origin time of *Buchnera* no earlier than that of their host, rather than assuming their ancestors co-occurred in the same time.

**Group:** *Blattabacterium*

**Lifestyle justification:** We classified all *Blattabacterium* as insect-associated.

**Justification:** *Blattabacterium* is a genus of obligate mutualistic endosymbiont bacteria inhabiting almost all species of cockroach known to date (Latorre et al. 2022) as well as the termite *Mastotermes darwiniensis* (Lo et al. 2007). In the insect host, *Blattabacterium* lives inside the fat cells of tissues in the abdominal cavity that store fat, designated as fat bodies. It likely plays a crucial role in nitrogen recycling, which is important for insects feeding on diets mainly composed of plant material where nitrogen supply is poor (Sabree et al. 2009). To our knowledge, no free-living members of this genus have been reported. Additionally, members of this genus all have highly reduced genomes. The above evidence strongly suggests an LCA of *Blattabacterium* that was associated with insects, mostly likely Blattodea, the order including cockroaches and termites.

### 3.2 Fossil- or biomarker-based time calibrations

Distinct from RTC which provides only information of the order of divergence, the time constraints described in this section are "real" and traditional time calibrations in molecular clock analysis since they provide absolute time bound(s) based on fossils or biomarkers to calibrate the time the corresponding node occurs.

#### 3.2.1 Time calibrations within eukaryotes

Eukaryote fossil calibrations consist of two sources: those used in our previous study which developed the mitochondria-based molecular dating approach and applied it to date α-proteobacteria evolution (Wang and Luo 2021), and those included in the present study based on recently discovered eukaryotic fossils. For simplicity, we give detailed justifications to only the two calibrations added in the present study (total group Chlorophyta and total group Nematoda), and readers interested in more details of other calibrations are encouraged to read our previous study (Wang and Luo 2021). For nodes where there are disputes in regard to the fossil records or their phylogenetic position, detailed discussions are given for why we favour some over others, and alternative fossil calibrations were also considered in molecular clock analysis (see "Alternatives" below).

**Node:** total-group Florideophyceae (Node 57 in Fig. S2A)
**Used in Wang and Luo 2021?:** Y
**Locality and Stratigraphy level:** Doushantuo Formation, southern China
**Minimum Age:** 550 Ma
**Maximum Age**: 1891 Ma
**Justification:** We followed Parfrey *et al.*, 2011 to use 550 Ma as the minimum age of the total group of Florideophyceae, which was based on the florideophyte fossils found at Doushantuo Formation (Xiao et al. 2004). We followed Morris et al., 2018 and Betts *et al.*, 2018 to set the soft maximum as 1891 Ma. This is based on the earliest fossil of simple eukaryotes (Zhongying 1986; Lamb et al. 2009; Peng et al. 2009), when no evidence of any organisms as complex as multicellular algae is reported to our knowledge (Betts et al. 2018; Morris et al. 2018). The formation's maximum age is based on the rocks the fossils overlie, dated at 1823 Ma ± 68 Ma (Lu et al. 1996).

**Node:** crown group Rhodophyta (total group Bangiophyceae) (Node 56 in Fig. S2A)
**Used in Wang and Luo 2021?:** Y
**Locality and Stratigraphy level:** Angmaat Formation, Bylot Supergroup of Baffin Island, arctic Canada
**Minimum Age:** 1047 Ma
**Maximum Age:** 1891 Ma
**Justification:** The fossils of *Bangiomorpha pubescens* are the oldest unambiguous fossil that can be confidently assigned to a major eukaryotic clade (Butterfield 2000; Gibson et al. 2018). We followed the study (Knoll 2011) to describe it as the calibration point of the total

group of Bangiophyceae, thus the crown group of red algae, as done in other studies (Parfrey et al. 2011; Yang et al. 2016). The minimum time bound was once established 1092 ± 59 Ma (Turner and Kamber 2012), but was recently revised to 1047 ± 0.032 Ma based on the Re-Os isotopic dates from sedimentary rocks stratigraphically bracketing the appearance of *Bangiomorpha pubescens* in the Bylot Supergroup of Baffin Island (Gibson et al. 2018). Following Morris et al., 2018 and Betts *et al.*, 2018, we set the soft maximum as 1891 Ma (see the node of the total-group Florideophyceae).

**Alternatives:** The oldest-possible fossils that represent red algae are *Ramathallus lobatus* and *Rafatazmia chitrakootia* (Bengtson et al. 2009; Bengtson et al. 2017), found in the Vindhyan Supergroup of central India, which were interpreted as red algae based on the presence of pyrenoids and pit plugs, respectively. However, the interpretation of them as red algae was questioned in several later studies (Betts et al. 2018; Gibson et al. 2018; Mills et al. 2022). Specifically, the claim of pyrenoids was difficult because such liquid phase structures often decay quickly and are difficult to be fossilized, and the pit-plug interpretation was also unlikely based on taphonomy experiments (Carlisle et al. 2021). Despite the controversy of *R. lobatus* and *R. chitrakootia*, we alternatively used the age of their fossils at 1600 Ma in the scheme (*Euk_fossil1* in Data S2). Further, since some scholars argued that the features thought to be characteristic of *Bangiomorpha* are also present in other red algae (Betts et al. 2018), we placed the calibration at the total-group red algae as alternative schemes (*Euk_redAlgaeTotalGrp* and *Euk_Betts* in Data S2).

**Node**: total-group eudicots (Node 55 in Fig. S2A)
**Used in Wang and Luo 2021?:** Y
**Locality and Stratigraphy level**: Cowleaze Chine Member, Isle of White
**Minimum Age**: 125 Ma
**Maximum Age**: 250 Ma
**Justification:** Tricolpate pollen represents the most ancient evidence of angiosperms. Following (Clarke et al. 2011a), we assigned the pollen to the Cowleaze Chine Member of the Vectis Formation, thus a minimum time of 126.3 ± 0.4 Ma. The soft maximum time constraint was based on sediments devoid of pollen resembling angiosperms below their first report in the Middle Triassic, approximately 247.1 Ma ± 0.2 Ma (Ogg 2012).

**Node:** total-group bryophytes (crown-group Embryophyta) (Node 54 in Fig. S2A)
**Used in Wang and Luo 2021?:** Y

**Locality and Stratigraphy level:** Qusaiba-1 core from the Quasim formation of northern Saudi Arabia

**Minimum Age:** 450 Ma

**Maximum Age:** 509 Ma

**Justification**: We followed Clarke *et al.*, 2011 to date trilete spores, which are the oldest evidence of embryophytes, to 450 Ma. The soft maximum constraint was placed at the Bright Angel Shale of the Tonto Group of Arizona, dated at 507.2-509 Ma (Baldwin et al. 2004).

**Alternatives:** Considering the debates on the maximum age of the crown group of land plants (Hedges et al. 2018), we followed (Su et al. 2021) to replace the maximum time bound 509 Ma by a more conservative time estimate at 1042 Ma which represents a sampled Precambrian locality where no plant-like spores were discovered (Turnbull et al. 1996) in the alternative dating scheme *Euk_fossil3* (Data S2).

**Node:** total-group Chlorophyta (Node 53 in Fig. S2A)

**Used in Wang and Luo 2021?:** N

**Locality and Stratigraphy level**: Nanfen Formation, Liaoning Province, northern China

**Minimum Age**: 947 Ma

**Maximum Age**: 1891 Ma

**Justification:** Chlorophyta is a major clade of green algae, the other being the recently proposed Prasinodermophyta (Li et al. 2020). The oldest fossils that represent Chlorophyta are those of *Proterocladus antiquus*. They were suggested to represent new benthic siphonocladalean chlorophytes in a recent study (Tang et al. 2020), which, according to the same study, display a suite of morphological features that collectively place them within the crown group Chlorophyta. These features include multicellularity and newly described characters such as cell differentiation, inferred siphonocladous construction, branching filaments, and a holdfast structure. However, the crown-group chlorophyte affinity of the fossils was questioned because it is difficult to justify, based on the original evidence, any interpretations more refined than total-group Chlorophyta (Mills et al. 2022). Hence, we conservatively placed them at the total group of Chlorophyta.

As to the age, according to the study (Tang et al. 2020), while no reliable radiometric ages that can be directly derived from the Nanfen Formation have been available, two pieces of indirect evidence can be used to constrain the age of the Nanfen Formation. This is established by the most recent population of detrital zircons derived from the underlying Diaoyutai Formation, dated at 1056 ± 22 Ma (Yang et al. 2012), and a diabase sill placed in the Qiaotou Formation it overlies, dated at 947.8 ± 7.4 Ma (Zhao et al. 2020). Because the Nanfen Formation is sandwiched between the overlying Qiaotou Formation and the underlying Diaoyutai Formation, the depositional age of the Nanfen Formation could be established as between 1056 and 947 Ma (Tang et al. 2020). Accordingly, we used 947 Ma as a conservative estimate of the minimum time bound for total-group chlorophytes.

**Node:** crown-group Amniota (total-group mammals) (Node 41 in Fig. S2A)
**Used in Wang and Luo 2021?:** Y
**Locality and Stratigraphy level:** Joggins Formation of Nova Scotia, Canada
**Minimum age:** 318 Ma
**Maximum age:** 332 Ma
**Justification:** The minimum age was set to be 318 Ma based on the date of the fossils of *Hylonomus lyelli* Dawson. We followed the study (Benton et al. 2015) to set the maximum as 332 Ma, the age of the fossiliferous Little Cliff Shale of the East Kirkton locality where no fossils representing any reptilians or mammals have been found.

**Node:** crown-group Chordata (total-group Cephalochordata) (Node 40 in Fig. S2A)
**Used in Wang and Luo 2021?:** Y
Locality and Stratigraphy level: Haikou [Yuanshan Fm (formerly Qiongzhusi)], China
**Minimum Age:** 520 Ma
**Maximum Age:** 636 Ma
**Justification:** We followed Benton *et al.*, 2015 to set the minimum based on the fossils of *Haikouichthys ercaicunensis* found in the Chengjiang Biota (Shu et al. 1999). The soft maximum was set based on the absence of Chordata fossils (and even eumetazoan fossils) in Lantian Biota, dated at 636 Ma (Yuan et al. 2011).

**Node:** total-group Nematoda (Arthropoda-Nematoda split) (Node 38 in Fig. S2A)
**Used in Wang and Luo 2021?:** N
**Locality and Stratigraphy level**: Yu'anshan Formation at Xiaotan section, Yongshan, southern China
**Minimum Age**: 514 Ma
**Maximum Age**: 636 Ma
**Justification:** This calibration was added in the molecular clock analysis of the present study because we included two arthropods (*Drosophila melanogaster* and *Daphnia pulex*), and one

nematode (*Caenorhabditis elegans*) for RTC-based dating involving symbionts of insects. Here, we chose to base the minimum time bound on *Yicaris dianensis*, an unequivocal member of crown group Crustacea based on the presence of a number of limb-based characters which are indicative of entomostracan and malacostracan affinity (Zhang et al. 2007; Benton et al. 2015). *Yicaris dianensis* was recovered from the Yu'anshan Formation at Xiaotan section, belonging to the *Eoredlichia-Wutingaspis* Biozone (Zhang et al., 2007). Chinese Cambrian stratigraphy, however, has been revised such that the Eoredlichia-Wutingaspis Biozone might not be recognized anymore (Peng 2009). Nevertheless, *Eoredlichia* is believed to co-occur with *Hupeidiscus* (early Cambrian eodiscoid trilobite), which is diagnostic of the *Hupeidiscus-Sinodiscus* Biozone, the second biozone of the Nangaoan Stage of the Qiandongian Series of the Cambrian of China. The Nangaoan is the proposed third stage of the Cambrian System for the International Geologic Timescale (Peng et al., 2012). Thus, we followed the study (Benton et al. 2015) to set a minimum bound based on the age estimate of the Nangaoan, dated at 514 Ma (Peng et al. 2012).

A soft maximum was set based on the maximum age interpretation of the Lantian Biota (Yuan et al. 2011), which together with the Doushantuo Biota (Luo 2005), provides a series of Lagerstätten (sedimentary deposits that show extraordinary fossils with exceptional preservation). However, none of them preserves any fossil that could be confidently described as even a total group eumetazoan. Therefore, the appearance time of arthropods and nematodes should not be earlier than Lantian Biota's maximum age, estimated to be 635.5 Ma ± 0.6 Mar (Condon et al. 2005). Accordingly, we followed the study (Benton et al. 2015) to set the soft maximum at 636 Ma.

**Alternatives:** It is also possible to establish a minimum time bound based on *Rusophycus* (Benton et al. 2015), dated at 528 Ma (Peng et al. 2012). *Rusophycus* fossils are widely believed to be produced by arthropod-grade animals, showing evidence of segmented limbs, an apomorphy of arthropods, as well as bilateral symmetry (Budd and Jensen 2000). As expected, the very small time difference between fossils of *Rusophycus* and of *Yicaris dianensis* resulted in highly similar time estimates (scheme *Euk_fossil4* Data S2).

**Node:** crown-group Metazoa (total-group Porifera) (Node 36 in Fig. S2A)
**Used in Wang and Luo 2021?:** Y
**Locality and Stratigraphy level:** White Sea Formation, Russia
**Minimum Age:** 550 Ma
**Maximum Age:** 833 Ma

**Justification:** The fossils of *Kimberella quadrata* are the oldest fossils of Bilateria, dated as 550 Ma. Following prior studies (Dos Reis et al. 2015; Betts et al. 2018), we establish a soft maximum of 833 Ma based on the Bitter Springs Formation of central Australia (Schopf 1968), dated at 827 ± 6 Ma, and the Svanbergfjellet Formation of Spitsbergen (Butterfield et al. 1994), both of which preserve various eukaryotic fossils but show no evidence for total group animals.

**Alternatives:** As shown in most phylogenetics studies (Pisani et al. 2015; Feuda et al. 2017; Kapli and Telford 2020; Redmond and McLysaght 2021), sponges are presumably the earliest-split lineage of animals [but see (Whelan et al. 2015; Li et al. 2021)]. Some scholars, however, based the minimum time of the crown group Metazoa on the biomarkers 26-methylstigmastane and 24-isopropylcholestane, which are dated at around 630 Ma (Parfrey et al. 2011). These fossil lipids were once considered to be produced by only sponges, but were later found to be common in rhizaria (a species-rich supergroup of unicellular eukaryotes), refuting the use of these biomarkers to indicate sponges (Nettersheim et al. 2019). A recent study reported putative sponges occurring in deep Proterozoic rocks (Turner 2021), dated at ~890 Ma (Van Acken et al. 2013). This was based on the presence of vermiform structures, originally defined as "in which narrow, sinuous, pale-coloured areas (usually of sparry carbonate) are surrounded by darker, usually fine-grained areas (usually carbonate)" (Walter 1972). However, the interpretation of the vermiform structure as spongin fibres of keratosan sponges, as made by the original study (Turner 2021), remains controversial. For example, Kris and McMenamin 2021 reinterpreted the same fossil as metazoan burrows formed by tiny worm-like animals instead of sponges. Further, the claim made by Turner 2021 was based on tiny holes in a piece of rock which may need further tomographic analysis to confirm. We alternatively calibrated the minimum age of the crown group animals based on the 890 Ma-old sponge as an alternative molecular clock analysis in the scheme *Euk_fossil2* (Data S2) to examine its impact on the molecular dating.

**Node:** total-group Fungi (crown-group Opisthokonta) (Node 34 in Fig. S2A)

**Used in Wang and Luo 2021?:** Y

Locality and Stratigraphy level: Brock Inlierm, the Northwest Territories, Canada

**Minimum age:** 890 Ma

**Maximum age:** 1891 Ma

**Justification:** The oldest fungal fossils are those of *Ourasphaira giraldae*, which came from the shale of Grassy Bay Formation (Shaler Supergroup, Arctic Canada), dated at 1010-890

Ma (Loron et al. 2019). Following Morris et al., 2018 and Betts *et al.*, 2018, we set the soft maximum as 1891 Ma (see the node of the total-group Florideophyceae).

**Node:** crown-group Dikarya (Basidiomycota-Ascomycota split) (Node 43 in Fig. S2A)
**Used in Wang and Luo 2021?:** Y
**Locality and Stratigraphy level:** Rhynie, Aberdeenshire, Scotland, Lower Devonian
**Minimum age:** 400 Ma
**Maximum age:** 1891 Ma
**Justification:** The most ancient uncontroversial fossils belonging to Dikarya are those of *Paleopyrenomycites devonicus*, which display clear characteristics of Ascomycota. The estimated date is ~400 Ma (Schoene et al. 2010; Mark et al. 2011), based on the age of the Rhynie Chert system. Following Morris et al., 2018 and Betts *et al.*, 2018, we set the soft maximum as 1891 Ma (see the node of the total-group Florideophyceae).

#### 3.2.2 Time calibrations within bacteria

In our previous study (Wang and Luo 2021), three cyanobacteria calibrations (total-group Pleurocapsales, total-group Nostocales, and total-group cyanobacteria) were adopted in the so-called "Cyanobacteria fossil-based approach", which dates α-proteobacteria evolution based on only cyanobacteria calibrations. In the present study, we additionally included another two, namely total-group Chromatiaceae and total-group Chlorobi based on biomarkers specific to Chromatiaceae and Chlorobi, respectively (Brocks et al. 2005). These five calibrations have been widely used to bacterial evolution in many studies (Battistuzzi and Hedges 2009; Louca et al. 2018; Boden et al. 2021; Liao et al. 2022). Note that none of these calibrations is associated with a well-established maximum time constraint (Zhang et al. 2021; Liao et al. 2022). Hence, following our previous study (Wang and Luo 2021), we set a soft maximum based on the age of Earth at ~4.5 Ga to all of them (alternative calibrations using the hard maximum were also tested and showed no difference in time estimates; *rootMaxHard* in Fig. S11).

**Node:** total-group Nostocales (Node a in Fig. S8)
**Used in Wang and Luo 2021?:** Y
**Minimum age:** 1600 Ma
**Maximum age:** 4500 Ma

**Justification:** We determined the minimum age of this calibration point based on ~1600 Ma-old fossils interpreted as akinetes in McArthur Group of Northern Australia (Tomitani et al. 2006). Akinetes are widely accepted as characteristic of Nostocales (de Tezanos Pinto et al. 2016). Following our previous study (Wang and Luo 2021), we set the maximum bound according to the age of Earth, which is ~4.5 Ga.
**Alternatives:** We also tried an alternative minimum bound, based on the 1200 Ma-old fossils recovered from the Middle Proterozoic Dismal Lakes Groups, Arctic Canada (Horodyski and Allan Donaldson 1980), as used in (Wolfe and Fournier 2018). Another alternative minimum bound was established based on the ~2000 Ma-old rod-like resting cell which might resemble akinetes found in the Franceville Group, Gabon (Amard and Bertrand-Sarfati 1997). However, its validity has been questioned (Butterfield 2015). The above two alternatives were used in the dating schemes *CyanoNos1200* and *CyanoNos2000* (Data S3), respectively. Further, we alternatively removed this calibration, treated the minimum time bound as a soft bound (2.5% probability to be beyond the bound), and calibrated the time priors with truncated Cauchy distribution instead of uniform distribution in the dating schemes *noCyanoFossil*, *CyanoSoft*, *CyanoCauchy* respectively (Data S3).

**Node:** total-group Pleurocapsales (Node b in Fig. S8)
**Used in Wang and Luo 2021?:** Y
**Minimum age:** 1700 Ma
**Maximum age:** 4500 Ma
**Justification:** The minimum time bound was based on the estimated age of the microfossils of Pleurocapsales recovered in Hebei Province, China, dated at ~1700 Ma, as used in some studies (Sánchez-Baracaldo 2015; Sánchez-Baracaldo et al. 2017). It is important to point out that these studies also used a maximum constraint based on coccoid and filamentous cell fossils, which, however, overlooks the time gap between the first occurrence of an apomorphic character and its first fossilization (Marshall 2019). As a result, this could lead to a false impression of precision as comprehensively discussed in (Zhang et al. 2021). Hence, following our previous study (Wang and Luo 2021), we set the maximum bound according to the age of Earth, which is ~4.5 Ga.
**Alternatives:** We alternatively removed this calibration, treated the minimum time bound as a soft bound (2.5% probability to be beyond the bound), and calibrated the time priors with truncated Cauchy distribution instead of uniform distribution in the dating schemes *noCyanoFossil*, *CyanoSoft*, *CyanoCauchy* respectively (Data S3).

**Node:** total-group oxygenic Cyanobacteria (Node c in Fig. S8)
**Minimum age:** 2320 Ma
**Maximum age:** 4500 Ma
**Justification:** The Great Oxidation Event (GOE), when $O_2$ started to accumulate in the Earth's atmosphere, profoundly altered Earth surface environments (Kump 2008). This is typically believed as a result of oxygenic photosynthesis conducted by oxygenic cyanobacteria (Kopp et al. 2005; Schirrmeister et al. 2013). Therefore, as done in prior studies (Zhang et al. 2021; Liao et al. 2022), the minimum bound for the total group of oxygenic cyanobacteria was set as the estimate time of the GOE at ~2320 Ma (Bekker et al. 2004). Note that some studies assigned the GOE-based calibration at the crown group of oxygenic cyanobacteria (Sánchez-Baracaldo et al. 2014; Sánchez-Baracaldo et al. 2017). This is likely because it was not until recent years that non-oxygenic lineages sister to oxygenic cyanobacteria, now classified as Vampirovibrionia (formerly known as Melainabacteria) and Sericytochromatia (Di Rienzi et al. 2013; Soo et al. 2017), were well recognized in genomics studies (see Zhang et al. 2021 for a more detailed critical discussion). Nonetheless, placing the 2320 Ma-minimum bound to the crown group of oxygenic cyanobacteria is inaccurate because it is possible that all modern oxygenic cyanobacterial lineages and thus the crown group of oxygenic cyanobacteria evolved after GOE (Soo et al. 2017; Fournier et al. 2021).
**Alternatives:** We alternatively removed this calibration in the dating scheme S and treated the minimum time bound as a soft bound (2.5% probability to be beyond the bound) in the dating schemes *noGOE* and *GOE_soft* (Data S3).

**Node:** total-group Chlorobi (Chlorobi-Bacteroidetes split) (Node d in Fig. S8)
**Minimum age:** 1640 Ma
**Maximum age:** 4500 Ma
**Justification:** We determined the minimum age of this calibration point based on the 1.64 Ga-old basin in northern Australia, the same as for Chromatiaceae (Brocks et al. 2005). The derivatives of aromatic carotenoid isorenieratane I and chlorobactane II are typically interpreted as biomarkers for green sulphur bacteria (Chlorobiaceae), implying euxinic conditions in the photic zone of the water column. For the BCF, renieratane, β-renierapurpurane and renierapurpurane are commonly interpreted as biomarkers for Chromatiaceae but note a potential contribution of Cyanobacterial synechoxanthin to the renierapurpurane pool (Brocks and Schaeffer 2008). While in principle isorenieratane could

be derived from Actinomycetes, these biomarkers in the BCF almost certainly derive from Chlorobiaceae for the following two reasons. First, Actinomycetes are only occasionally abundant in marine habitats (Ward and Bora 2006). Second, Actinomycetes contain only isorenieratene and its derivatives, which are among the least abundant aromatic carotenoids in the BCF, but not the precursors of chlorobactane II or any other aromatic carotenoids found in the BCF, which are very abundant in the BCF (Brocks and Schaeffer 2008). The above has been used to calibrate the minimum time bound of Chlorobi-Bacteroidetes split in several bacteria molecular dating studies (Battistuzzi and Hedges 2009; Louca et al. 2018; Hugoson et al. 2022). We set the maximum bound according to the age of Earth, which is ~4.5 Ga.

**Alternatives:** Further, we alternatively removed this calibration, treated the minimum time bound as a soft bound (2.5% probability to be beyond the bound), and calibrated the time priors with truncated Cauchy distribution instead of uniform distribution in the dating schemes *noBiomarker*, *BiomarkerSoft*, *BiomarkerCauchy* respectively (Data S3).

**Node:** total-group Chromatiaceae (Node e in Fig. S8)

**Minimum age:** 1640 Ma

**Maximum age:** 4500 Ma

**Justification:** Carbonates preserved in the 1.64 Ga-old Barney Creek Formation (BCF) of the McArthur Group, northern Australia (Page and Sweet 1998), were found to contain more than 22 types of $C_{40}$ carotenoid derivatives, e.g., β-carotane, γ-carotane, chlorobactane, lycopane, isorenieratane, β-isorenieratane, renieratane, β-renierapurpurane, and okenane. This greatly extends the geological record of the derivatives of carotenoid by more than 1 Ga. Specifically, okenane is believed to be exclusively produced by several genera of phototrophic purple sulphur bacteria (Chromatiaceae, γ-Proteobacteria), and accordingly, it has been interpreted as a biomarker for Chromatiaceae (Imhoff 2006; Brocks and Schaeffer 2008). The above has been used to calibrate the minimum time bound of the total-group Chromatiaceae in several bacteria molecular dating studies (Battistuzzi and Hedges 2009; Louca et al. 2018; Hugoson et al. 2022). We set the maximum bound according to the age of Earth, which is ~4.5 Ga.

**Alternatives:** Further, we alternatively removed this calibration, treated the minimum time bound as a soft bound (2.5% probability to be beyond the bound), and calibrated the time priors with truncated Cauchy distribution instead of uniform distribution in the dating schemes *noBiomarker*, *BiomarkerSoft*, *BiomarkerCauchy* respectively (Data S3).

1276

### Supplementary Note 4. Simulations to evaluate the usefulness of the bootstrap approach to incorporating Cxx models in dating

#### 4.1 Generation of simulated datasets

We tested the performance of the bootstrap-based approach to incorporating the site-heterogeneous profile mixture models Cxx (C10-C60) (Quang et al. 2008) on simulated data reflecting deep-time evolution. For each of the three substitution models used to simulate sequence evolution (see below), we employed TreeSim (Stadler 2011) to generate 30 timetrees each with 30 tips under a birth-death model with the birth and death rate respectively as 0.4 and 0.2 lineages per 100 Ma, as estimated by a previous study for bacterial evolution (Scholl and Wiens 2016), and taxon sampling proportion as 10%. The simulated trees were generated with a true root age of 1.0, 2.0, 3.0, and 4.0 Ga (close to Earth' age), corresponding to evolution on different time scales.

For each timetree, branch-wise substitution rates were drawn from a lognormal distribution (the independent rate clock model in MCMCTree) with the parameters $\mu = -3.7$ and $\sigma = 0.2$. In other words, the log-transformed values of branch rates follow a normal distribution parameterized by $\mu = -3.7$ and $\sigma = 0.2$. Thus, the mean of the lognormal distribution is $\exp(\mu + \frac{\sigma^2}{2}) \approx 0.25$ substitutions per site per Ga and variance of $[\exp(\sigma^2) - 1]\exp(2\mu + \sigma^2) \approx 0.005$ (Taboga 2022). This corresponds to the observation that tip-to-root distance in phylogenomic tree of bacteria is around 1.0 amino acid substitution per site (Moody et al. 2022), assuming a 4 Ga-old LCA of bacteria ($0.25 \times 4 = 1$). Alignments of 1000 amino acids were generated using AliSim (Ly-Trong et al. 2022) implemented in IQ-Tree under different models. Note that C60 was not used in simulation to save time.

i) LG+G{1.0}: LG substitution model and across-site rate variation under a four-category discrete gamma distribution with the shape parameter α=1.0, thus Gamma(1.0,1.0), meaning that the mean and variance of the relative rates across site are $\frac{1}{1} = 1$ and $\frac{1}{1^2} = 1$ respectively (Yang 1994).

ii) LG+G{1.0}+C40: LG substitution model plus a mixture of amino acid site frequency profiles C40 and across-site rate variation under a four-category discrete gamma distribution with α=1.0.

iii) LG+G{0.5}+C40: LG substitution model plus a mixture of amino acid site frequency profiles C40 and across-site rate variation under a four-category discrete gamma distribution with α=0.5, which reflects more among-site rate variation than using α=1.0.

### 4.2 Calibrations used in MCMCTree analysis on simulated datasets

We ran MCMCTree analyses on simulated sets with a single calibration at the root, or with a root calibration plus two internal calibrations. The two internal calibrations were chosen as the nodes at the 1/3 and 2/3 quantiles of the ages in the "true" timetree (Fig. S5). The time priors of all calibrated nodes were set to be uniform within the interval [true_age - (true_age/5), true_age + (true_age/5)] with soft bounds on both upper and lower time bounds, thus a probability of 2.5% (by default) that the age is outside the bound.

### 4.3 Substitution models to test on simulated datasets

We tested the performance of using different substitution models in time estimation using MCMCTree's approximate likelihood method (dos Reis and Yang 2011) on simulated datasets generated described as above. The following four ways in estimating the branch length and the hessian matrix (second-order derivatives of the log likelihood) were compared.

i) LG+G (MCMCTree): branch lengths and hessian both directly calculated by MCMCTree under LG+G.
ii) LG+G (bs): branch lengths calculated by IQ-Tree's LG+G and hessian approximated by bootstrapping.
iii) LG+G+C20 (bs): branch lengths calculated by IQ-Tree's LG+G+C20 and hessian approximated by bootstrapping.
iv) LG+G+C40 (bs): branch lengths calculated by IQ-Tree's LG+G+C40 and hessian approximated by bootstrapping.

In summary, we simulated 30 alignments under each of the above three substitution models. Four root ages were tried respectively in simulating each timetree. This added up to $30 \times 3 \times 4 = 360$ alignments. As to MCMCTree analysis, it was run on each alignment with or without internal calibrations, and under each of the above four ways in estimating branch lengths and the hessian matrix. Collectively, this means $360 \times 2 \times 4 = 2880$ MCMCTree analyses.

### 4.4 Comparing the performance of different substitution models on simulated datasets

Two indices were used to test the performance of different substitution models by comparing the branch lengths (i.e., time spanned between the two adjacent nodes) obtained by MCMCTree with a given substitution model, to those used in simulation (true values).

i) Branch score distance (BSD): defined in the study (Kuhner and Felsenstein 1994) as $\sqrt{\sum_i (b_i^m - b_i^{true})^2}$, where $i$ traverses all branches, and $b_i^m$ and $b_i^{true}$ denote the average time spanned between the two adjacent nodes connected by branch $i$ in the posterior timetrees inferred by MCMCTree under substitution model $m$, and those used in simulation (true values), respectively. The lower the value is, the more accurate the time estimates using the substitution model $m$ are.

ii) Relative difference: calculated as $mean\left(\frac{|b_i^m - b_i^{true}|}{\max(b_i^m, b_i^{true})}\right)$, where $b_i^m$ and $b_i^{true}$ are defined in the same way as defined in the BSD. The lower the value is, the more accurate the time estimates using the substitution model $m$ are.

We first compared the time estimation between using MCMCTree's default method to calculate the hessian matrix, and our approach to approximating the hessian matrix by bootstrapping, under the same substitution model LG+G, i.e., *LG+G (MCMCTree)* vs. *LG+G (bs)* in Fig. S9. In all analyses, the time estimates by these two were highly similar as indicated by the above two indices ($P$-value > 0.05 Wilcoxon test). This indicates that our bootstrap-based approach can well replace those directly calculated by MCMCTree in time estimates (but see Note 2.5 for limitations).

Second, we compared the performance in time estimation between using LG+G and LG+G+C40. In case the true root age of the simulated timetree was 1.0 or 2.0 Ga, or if the model used to simulate sequence evolution was LG+G, the results estimated using both approaches yielded highly similar results (Fig. S9). However, when the simulation substitution model was LG+G+C40 and the true root age was set as 3.0 or 4.0 Ga, LG+G+C40 obtained significantly more accurate time estimates than LG+G if no internal calibration was provided (red boxplots in Fig. S6; LG+G vs. LG+G+C40). In case of two internal calibrations, the improvement in time estimation accuracy by using LG+G+C40 was reduced, but was still statistically different and notable (green boxplots in Fig. S9; LG+G vs. LG+G+C40). Moreover, LG+G+C20 and LG+G+C40 performed equally well even when the simulated alignment was generated by LG+G+C40.

Third, as to the estimate of the absolute rate (unit: substitutions/site/Ga), for sequences simulated under LG+G, LG+G obtained more accurate estimates of the rate than LG+G+Cxx, while LG+G+Cxx tended to overestimate the rate. On the other side, for sequences simulated under LG+G+Cxx, LG+G+Cxx showed accurate estimates of the rate, while LG+G likely underestimated the rate.

The above simulation analysis implies that the site-heterogeneous model LG+G+Cxx outperformed MCMCTree's default LG+G for dating deep phylogenies of bacteria, which plausibly dates back to more than 3.0 Ga and where the sequence evolution might be under more complex substitution models than LG+G (Gouy et al. 2015; Martijn et al. 2018; Wang et al. 2019; Moody et al. 2022). It needs to be emphasized that we do not mean that the "true" model is LG+G+Cxx, but that LG+G+Cxx is closer to the "true" model and better fits the data as indicated by AIC or BIC. In other words, for deep-time evolution, the sequence is often (much) more likely to be generated by LG+G+Cxx than LG+G. Further, that the difference in time estimates between LG+G and LG+G+Cxx became smaller in case of a few reliable internal calibrations indicates the usefulness of internal calibrations (green boxplots in Fig. S9). This is likely because underestimation of branch lengths by too simple models (e.g., LG+G) can be balanced by multiple reliable calibrations (Groussin et al. 2011; Tao et al. 2020) but at the cost of underestimates of the evolutionary rate.

### 4.5 Technical comments on the bootstrap-based approach to incorporating Cxx model in MCMCTree

Note that to generate the hessian matrix used by the approximate likelihood method of MCMCTree, our bootstrap method involves calculating the inverse matrix of the covariance matrix of branch length estimates. This may fail if the covariance matrix is invertible, or in other words, if its determinant equals zero (see any linear algebra textbook). This happens particularly if organisms involved in the analysis are phylogenetically closely related such that their sequences are extremely similar. In such cases, the MLE of their branch length might be very close to zero, and the phylogenetics program must assign the branch length a very small value which in IQ-Tree is $10^{-6}$ by default. Accordingly, all values of the corresponding row in the covariance matrix equal zero. As illustrated by the following matrix: assume that the $i^{th}$ branch has the same branch length across all bootstraps, so the covariance between the branch length estimate $\hat{\theta}_i$ and any others equals zero. Accordingly, the determinant of the bootstrap covariance matrix is zero, and the matrix is therefore invertible.

$$Var(\hat{\theta}) = \begin{bmatrix} Cov(\hat{\theta}_1, \hat{\theta}_1) & \cdots & Cov(\hat{\theta}_1, \hat{\theta}_{i-1}) & 0 & Cov(\hat{\theta}_1, \hat{\theta}_{i+1}) & \cdots & Cov(\hat{\theta}_1, \hat{\theta}_n) \\ \vdots & \ddots & \vdots & \vdots & \vdots & \ddots & \vdots \\ Cov(\hat{\theta}_1, \hat{\theta}_{i-1}) & \cdots & Cov(\hat{\theta}_{i-1}, \hat{\theta}_{i-1}) & 0 & Cov(\hat{\theta}_{i-1}, \hat{\theta}_{i+1}) & \cdots & Cov(\hat{\theta}_{i-1}, \hat{\theta}_n) \\ 0 & \cdots & 0 & 0 & 0 & \cdots & 0 \\ Cov(\hat{\theta}_1, \hat{\theta}_{i+1}) & \cdots & Cov(\hat{\theta}_{i-1}, \hat{\theta}_{i+1}) & 0 & Cov(\hat{\theta}_{i+1}, \hat{\theta}_{i+1}) & \cdots & Cov(\hat{\theta}_{i+1}, \hat{\theta}_n) \\ \vdots & \ddots & \vdots & \vdots & \vdots & \ddots & \vdots \\ Cov(\hat{\theta}_1, \hat{\theta}_n) & \cdots & Cov(\hat{\theta}_{i-1}, \hat{\theta}_n) & 0 & Cov(\hat{\theta}_{i+1}, \hat{\theta}_n) & \cdots & Cov(\hat{\theta}_n, \hat{\theta}_n) \end{bmatrix}$$

Further, the assumption that all gradients equal zero adopted in our bootstrap-based approach may be violated if any branch takes zero length (for details see Reis and Yang 2011). Fortunately, in terms of building deep phylogenies, this phenomenon should be uncommon because closely related strains are less likely to be involved. However, it is still advised to carefully select the sequences and organisms to avoid extremely short branch before using this method. Similarly, increasing the number of bootstraps may also help since running with a small number of bootstraps can easily cause the lengths of certain branches to be the same in all bootstrap trees. For our dataset, 1000 bootstraps appeared enough.

Although increrasingly appraciated phylogenetic reconstruction, to our knowledge, the profile mixture model (CAT and Cxx) has been implemented in only one molecualr clock software PhyloBayes (Lartillot et al. 2009). However, PhyloBayes implements only the exact likelihood method in molecular clock analysis which is much slower than the MCMCTree's approximate likelihood method, and thus may not meet the need for molecular dating in the genomics era where genome-scale data becomes increasingly available (Dos Reis et al. 2016). Specifically, MCMCTree's approximate likelihood method provides a 1000× speed-up (dos Reis and Yang 2011) and ~100× less $CO_2$ emission (Álvarez-Carretero et al. 2022) over traditional MCMC molecular dating software while obtaining similar accuracy. Hence, the bootstrap-based approach to estimating the hessian matrix used in MCMCTree's approximate method developed in the present study provides an easy and flexible way to integrate profile mixture model in molecular dating, thereby allowing more accurate, faster, and greener computation in deep-time molecular clock analysis using genome-scale datasets.

### Supplementary Note 5: Additional discussion

#### 5.1 An inferred evolutionary timeline of eukaryotes

Our estimate of an origin of crown-group eukaryotes at ~1600 Ma is consistent with many previous time estimates (Parfrey et al. 2011; Eme et al. 2014; Betts et al. 2018). A recent study dated a much earlier origin of eukaryotes at ~2100 Ma (Strassert et al. 2021). However, note that this study used a relatively new calibration that was not used as calibration in most previous molecular clock studies. Specifically, this calibration calibrates the minimum time of the crown group of red algae (more specifically Rhodophytina, as subdivision of red algae) to be 1600 Ma based on the fossil of *Rafatazmia* (Bengtson et al. 2017). This is roughly 400 Ma (Parfrey et al. 2011; Yang et al. 2016) to 550 Ma (Betts et al. 2018; Gibson et al. 2018; Wang and Luo 2021) earlier than used in most prior studies based on the fossil of *Bangiomorpha pubescens* (Butterfield 2000). As shown in Fig. S3 (*Euk_fossil1*), when we kept all other calibrations unchanged but calibrated the minimum bound of crown-group red algae to be 1600 Ma instead of 1047 Ma as used in the focal molecular clock analysis, the posterior ages of most (deep) eukaryote nodes shifted toward the past, and the root age increased to ~2100 Ma. The new time estimate was similar to those reported in Strassert et al. 2021. This suggests that the use of this calibration, which is the oldest among all calibrations used in the present as well as many other (Parfrey et al. 2011; Betts et al. 2018; Wang and Luo 2021) molecular clock analyses for eukaryotes, might explain the different time estimates obtained in different studies.

Note that our molecular clock analysis does not invalidate the alternative calibration based on the 1600 Ma-old fossil *Rafatazmia*. To say the least, the posterior ages of eukaryotes estimated in the first-step sequential analysis using a 1600 Ma-old minimum bound for red algae crown group had little impact on the time estimates of bacteria (*Euk1* in Fig. 3), suggesting that our time estimates of the bacterial tree were robust to the use of alternative eukaryotic fossils in general.

#### 5.2 Caveats in reconstruction of ancestral lifestyle for modern bacterial symbionts

There are several caveats in our ASR analysis. First, our model assumes the ancestral bacterium took only one host or lifestyle at one time. While it is straightforward to assume so, it is possible that some symbionts adapt to different lifestyles at different life stages or under different environmental conditions. This is epitomized by rhizobia, which can form $N_2$-fixing nodules with legumes but which can also live in soils in the absence of compatible legume host. Further, while bacterial symbionts isolated from animals are typically classified as animal-associated, some might actually be symbionts of the protists that live in the gut or other parts of the animals (Collingro et al. 2020). One example is *Elusimicrobia*, which are identified as endosymbionts of various flagellated protists dominating the gut community of termites and ants (see also Note S3.1). Hence, in the present study we confidently classified those found from animal samples as protist-associated symbionts for *Elusimicrobia*. While such cases may also be present in other bacterial clades, further evidence is needed. Additionally, some bacteria found in animal organs like gut may be transient free-living bacteria, as these kinds of organs are connected to external environments.

In sum, mis-classification of bacterial lifestyles may indeed occur, which could affect ASR. Nonetheless, note that the internal nodes that served as the RTCs were selected from classical "symbiont clades" based on not only ASR, but comprehensive literature review (Note S3.1). Also, the consistent results obtained when alternative ways of lifestyle classification and parameters in ASR were used suggest that the above caveats likely did not have a large impact on the posterior date estimate in the RTC framework. This is likely because of the many RTCs included such that any change in a few of them unlikely affected the overall pattern.

2028

2029
